# Learning epistatic polygenic phenotypes with Boolean interactions

**DOI:** 10.1101/2020.11.24.396846

**Authors:** Merle Behr, Karl Kumbier, Aldo Cordova-Palomera, Matthew Aguire, Omer Ronen, Chengzhong Ye, Euan Ashley, Atul J. Butte, Rima Arnaout, Ben Brown, James Priest, Bin Yu

**Affiliations:** Department of Statistics, University of California at Berkeley, Berkeley, CA, USA; Department of Pharmaceutical Chemistry, University of California, San Francisco, CA, USA; Department of Pediatrics, Stanford Medicine, Stanford, CA, USA; Department of Biomedical Data Science, Stanford Medicine, Stanford, CA, USA; Division of Cardiovascular Medicine, Stanford Medicine, Stanford, CA, USA; Bakar Computational Health Sciences Institute, University of California, San Francisco, CA, USA; Division of Cardiology, Department of Medicine, University of California, San Francisco, CA, USA; Biosciences Area, Lawrence Berkeley National Laboratory, Berkeley, CA, USA; Tenaya Therapeutics, San Francisco, CA, USA; Department of Electrical Engineering and Computer Sciences and Center for Computational Biology, University of California at Berkeley, Berkeley, CA, USA; Faculty of Informatics and Data Science, University of Regensburg, Regensburg, Germany

## Abstract

Detecting epistatic drivers of human phenotypes is a considerable challenge. Traditional approaches use regression to sequentially test multiplicative interaction terms involving pairs of genetic variants. For higher-order interactions and genome-wide large-scale data, this strategy is computationally intractable. Moreover, multiplicative terms used in regression modeling may not capture the form of biological interactions. Building on the Predictability, Computability, Stability (PCS) framework, we introduce the epiTree pipeline to extract higher-order interactions from genomic data using tree-based models. The epiTree pipeline first selects a set of variants derived from tissue-specific estimates of gene expression. Next, it uses iterative random forests (iRF) to search training data for candidate Boolean interactions (pairwise and higher-order). We derive significance tests for interactions, based on a stabilized likelihood ratio test, by simulating Boolean tree-structured null (no epistasis) and alternative (epistasis) distributions on hold-out test data. Finally, our pipeline computes PCS epistasis p-values that probabilisticly quantify improvement in prediction accuracy via bootstrap sampling on the test set. We validate the epiTree pipeline in two case studies using data from the UK Biobank: predicting red hair and multiple sclerosis (MS). In the case of predicting red hair, epiTree recovers known epistatic interactions surrounding *MC1R* and novel interactions, representing non-linearities not captured by logistic regression models. In the case of predicting MS, a more complex phenotype than red hair, epiTree rankings prioritize novel interactions surrounding *HLA-DRB1*, a variant previously associated with MS in several populations. Taken together, these results highlight the potential for epiTree rankings to help reduce the design space for follow up experiments.

## Introduction

Epistasis between genetic alleles describes a non-additive relationship among different loci governing a single trait [1]. While epistatic interactions have been hypothesized to play an important role in regulating phenotypes [2, 3], most large-scale studies on polygenic contributions have focused on additive effects. Discovering epistatic interactions involved in human phenotypes has been a slow, small-scale process with relatively modest or minimal evidence from large-scale studies [4, 5].

The most commonly accepted statistical definition of epistasis dates back to Fisher [6]: the “deviation from the addition of superimposed effects (…) between different Mendelian factors.” However, Fisher’s definition does not correspond to a well-defined statistical model for either the null (no-epistasis) or alternative (epistasis) hypotheses. Additivity depends on the scaling of a response (e.g., penetrance for a binary trait). For example, a multiplicative function becomes additive on a log-scale.

Moreover, for a fixed scaling there are many ways to write an additive model of individual components. For example, taking the inverse normal or other transformation of each feature before running logistic regression. Although these issues have been noted repeatedly in the literature [7–11], their impact on statistical results are often not highlighted.

While there is no unique model of Fisherian epistasis, traditional approaches evaluate epistasis through linear (continuous response) or logistic (binary response) regression with two genetic variants (possibly incorporating covariates for population structure and genetic-linkage) [12–15]. In this setting, the null (no-epistasis) model has linear additive components for the two involved genes and the alternative (epistasis) model has an additional multiplicative interaction term. However, complex phenotypes can involve more than two genes and the functional relationship between genes and phenotype can be highly complex. Interactions identified as highly significant by brute-force, pairwise searches with logistic/linear regression may not always provide a good fit to the data, and may fail to generalize to new/unobserved samples. Moreover, statistical hypothesis testing based on heavily mis-specified null or alternative models are often unstable and can lead to irreproducible results [16–19].

Beyond the challenges of modeling complex interaction forms, the massive search space of genomic interactions presents several barriers to classical approaches. First, brute force searches can become computationally intractable for genome-scale data. This is a particular issue when considering high-order (i.e., beyond pairwise) interactions. Second, the standard approach of modeling a single interaction at a time may lead to model misspecification by failing to take the entire genetic background into account. Finally, analyses typically limit their focus to marginally important variants due to high-dimensional genomic data. Interactions involving variants with weak marginal effects cannot be detected in this setting.

Here, we propose the epiTree pipeline, which makes four key contributions to the challenge of detecting polygenic, epistatic interactions in genome-scale data. First, epiTree makes a conceptual contribution: using data splits to separate the discovery of interactions, performed on training data, from statistical inference, performed on testing data. Second, epiTree uses a two-step, biologically inspired dimension reduction to tractably search for Boolean, epistatic interactions of arbitrary size, considerably reducing the search space of high-order interactions. Third, epiTree fits tree-based models to learn data-adaptive representations of interactions and evaluate deviations from additivity directly on the penetrance scale. Fourth, epiTree builds on the recently proposed predictability, computability, and stability (PCS) framework [20] to derive an inference procedure, epiTree test. epiTree test is used to obtain p-values for interactions that demonstrate accurate prediction on the hold-out test set without relying on chi-square distributional approximations, which can suffer from poor distributional tail approximations under model misspecification [21].

We validate the epiTree pipeline in two case studies of UK Biobank data: (i) predicting red hair and (ii) predicting mutliple sclerosis (MS). In the red hair case study, pairwise epistatic interactions among unlinked genes surrounding *MC1R* have been previously reported [15]. We rediscover *MC1R*-related interactions and discover new interactions, for example pairwise interactions among genes that were not previously associated with red hair (e.g. *UPF3A*, *SIAH2*); pairwise interactions between genes individually associated with hair color but not previously reported as interacting; as well as high-order (up to order-4) interactions which other methods cannot typically detect. In the MS case study, we discover several pairwise and one order-3 interaction among unlinked genes surrounding *HLA-DRB1*, which has been associated with MS across several populations [22–24], and involving genes associated with biological processes implicated in MS. We note that epistatic interactions in the MS case study are less-studied, see however [25–27]. The ranked epistastic interactions by our PCS p-values serve as a predictive and stable source of evidence for designing follow-up experiments to confirm such interactions. We believe the proposed epiTree pipeline reduces the design space of such experiments compared to standard analyses of epistasis.

## Materials and methods

### Ethics statement

This research was covered by the UK Biobank ethics agreement. UK Biobank has approval from the North West Multi-centre Research Ethics Committee (MREC). Complete details of all UK Biobank procedures are available at www.ukbiobank.ac.uk. All participants provided written informed consent.

### Phenotype and genotype data from the UK Biobank

As positive control case study for epiTree, we analyzed epistatic interactions in UK Biobank data associated with a previously-studied, genetically-driven phenotype: red hair [15]. We considered a total of 337,535 unrelated white British individuals and their self-reported hair color (Data field 1747; 15,326 with `Red` hair: positive cases and 322,209 individuals with `Blonde`, `Light Brown`, `Dark Brown`, `Black`, or `Other` hair color: controls). We searched for interactions in a balanced, random sample of cases and controls (15,000 red hair individuals and 15,000 controls). To assess the generalizability of our results to unobserved data, we performed a random sample split into training and test sets with 26K training samples and 4K test samples. We note that, in general, one might prefer non-randomly to randomly sampled test sets—e.g., a data from an external experiment. Such an external data set was non available to us in this setting.

We modeled hair color using *∼* 15, 000, 000 common variants imputed to the Haplotype Reference Consortium (HRC) and UK10K reference panels from *∼* 800, 000 directly genotyped variants, which were obtained by UK Biobank using one of two similar arrays [28]. Details regarding the ascertainment and quality control of these genotypes have been previously described [15, 28]. In brief, genotyped variants were subject to outlier-based filtration on effects due to batch, plate, sex, array, as well as discordance across control replicates. Samples with excess heterozygosity or missingness were excluded from the data release. Imputed variants were further subject to filtration based on Hardy-Weinburg equilibrium, missingness in white British individuals, minor allele frequency (*>* 10*^-^*^4^), and imputation quality. Respective details on phenotype and genotype data for the MS case study can be found in SI Section S1.8.

### Two-step interaction screening with biologically inspired dimension reduction

We developed a two-step procedure for screening interactions to improve computational efficiency and help stabilize of results. First, we aggregated SNP-level information by gene to search for gene-level interactions associated with the target phenotype. This initial step (i) substantially reduced dimensionality of the data and (ii) detected interactions at the level of a biologically interpretable unit. Second, variants surrounding putatively interacting genes were selected to search for SNP-level interactions.

#### Gene-level interactions

For the red hair case study we estimated gene expression levels in skin tissue (see SI Section S1.8 for respective details on the MS case study), which captures melanocyte biology and pigmentation, from the individual SNP data using PrediXcan [29]. In brief, PrediXcan predicts tissue-specific gene expression levels using elastic net models trained on GTEx v7 data [30]. From a statistical perspective, this mapping serves as dimensionality reduction step, reducing the number of genetic features from *∼* 10^7^ variants to *∼* 10^4^ genes. We then searched for non-linear interactions between estimated expression levels and the target phenotype using iterative random forests (iRF) [31] (see below).

#### SNP-level interactions

We selected variants within +/- 1MB of putatively interacting genes and re-ran iRF over this reduced subset to search for candidate SNP interactions. The selection of 1MB is a trade-off between (i) being large enough to include variants surrounding a gene that may also be associated with the response and (ii) being small enough to result in a reasonable absolute number of SNP features. From a statistical perspective, it would be sufficient to only select variants included in putatively interacting genes’ PrediXcan models. However, it is possible that variants within a gene region but not included in the PrediXcan model exhibit stronger interactive effects.

The overall two-step procedure is illustrated in Fig 1. Our procedure identified variant- and gene-level interactions that capture nonlinear dependencies over arbitrary sets of variants/genes. The ability to model interactions among any set of genes arises directly from our use of iRF, which models the joint contribution of all genes simultaneously. We also explored a complementary one-step approach to directly search for variant-level interactions, by pre-filtering variants based on iRF applied to variant-sub-batches, which is discussed in further detail in the Supplementary Information (SI).

**Fig 1.**
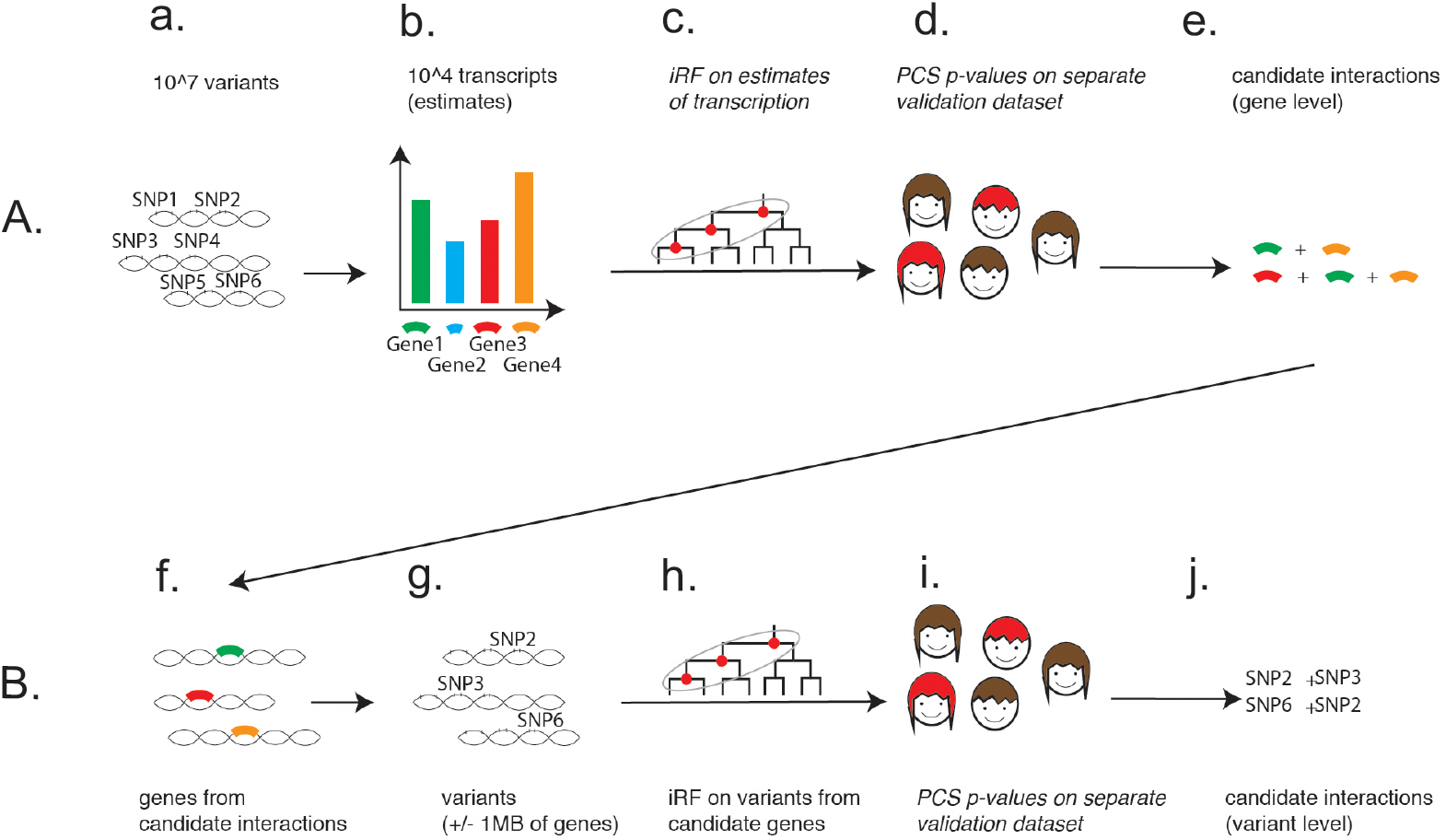
Illustration of the two-step procedure, from gene level analysis (A.) to variant level analysis (B.), for extraction of candidate interactions. A. First row from left to right: from approx. 10^7^ variants (a.) tissue specific transcripts for approx. 10^4^ genes (b.) are imputed using the software PrediXcan. Then the epiTree pipeline is applied to extract interactions for the gene expression features. B. Second row from left to right: for all genes that appear in interactions from the first step (f.), variants within 1MB of the start or end of an interacting gene are extracted (g.). Then the epiTree pipeline is applied to extract interactions for the variant features. For both, gene and variant level analysis, the epiTree pipeline first applies iRF (c./h.) to extract candidate interactions and then calculates PCS p-values for these candidate interactions on hold-out test data (d./i.). This results in the final set of selected gene-level interactions (e.) and variant level interactions (j.).

It is important to note that individual SNP weights in PrediXcan models are based on each SNP’s association with a gene’s expression levels. Our proposed approach is therefore explicitly focused on detecting interactions where changes in gene expression levels are non-additively associated with the target phenotype. By filtering to these gene-level interactions in the first stage, we expect to detect SNP-level interactions among variants that are also associated with changes in gene. Variants that result in functional changes to a protein without influencing its expression level may be missed by our approach.

### iRF non-linear model selection

Given a set of features (either estimated gene expression or individual genetic variants), we applied iRF [31, 32] as a non-linear model selection step to extract candidate interactions from the training data. The iRF algorithm fits a series of feature-weighted random forests (RF) [33] to iteratively stabilize feature selection along the decision paths of trees in the forest. After the final iteration, iRF identifies stable candidate interactions by searching for features that frequently co-occur on decision tree paths using random intersection trees (RIT) [34]. In addition to the interactions explicitly identified by iRF, we further expanded our set of candidate interactions for PCS epistasis inference (see below) by taking all inter-chromosome pairwise interactions among the top 50 iRF genes (with respect to Gini importance).

A key benefit of iRF is that the computational complexity of searching for interactions does not grow exponentially with the size of an interaction. This allowed us to identify high-order (i.e. beyond pairwise) Boolean interactions in a computationally tractable manner. Moreover, accuracy of iRF predictions on the hold-out test data provides a measure of generalizability for our non-linear model selection method, and correspondingly, the identified interactions. Finally, iRF detects known interactions in both simulation and biological case studies [31, 32, 35, 36] and analytically tractable versions of iRF yield provably consistent interaction discovery [37].

### Predictability, Computability, Stability (PCS) inference for epistatic interactions based on Boolean models

The significance of a candidate interaction for a binary phenotype is traditionally evaluated through logistic regression (see SI Section S1.1.2 for a recap). However, the model imposed by logistic regression—including a response rescaling, transformations of gene expression data, and the polynomial interaction term—does not arise from a known biological observation or mechanism. Indeed, additive or multiplicative forms may not capture the stochastic nature of living systems or threshold effects commonly observed in molecular and cellular phenomena [38–41]. A mathematically inaccurate representation of the biological mechanism of action between multiple genetic loci may result in unstable and irreproducible results that provide a poor fit of the data.

To address these issues, we developed a PCS epistasis p-value to evaluate. The PCS framework, recently proposed in [20], is based on the three core principles of data science: Predictability, Computability and Stability (PCS). It unifies and expands ideas from machine learning and statistics by using predictivity as a universal reality (or model) check, appropriate computational strategies for every step of an analysis process including simulating realistic reference/null distributions, and stability analysis to assess reproducibility at every stage of an analysis—from problem formulation/data collection, to modeling including inference, and to post-hoc model interpretations. PCS epistasis p-values are based on Boolean models of non-epistasis (null) and epistasis (alternative), learned adaptively from training data, and evaluated on hold-out test data. This approach has the advantage of being able to learn flexible interaction forms from the training data, rather than assuming multiplicative interactions that lack biological justification [8, 9, 11, 42].

#### No epistasis (null) model

Consider an interaction between genes A and B, (or a collection of interacting features in the set). We define the non-epistasis (null) model as an additive combination of decision trees fit to individual features using the training set samples and combined with backfitting:

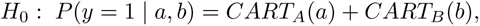

where CART denotes a “Classification And Regression Tree” (see [43]). We stress that the null model *H*_0_ also covers the case where the response *P* (*y* =1 *| a, b*) only depends on one of the genes A or B, or on neither of them, with *CART_A_* and/or *CART_B_* equal to a constant.

For higher-order interactions, say of the feature set *I* with *|I| >* 2, the null model can be defined in a similar way. In this case however, lower-order interactions among features *I^0^* (*I* must also be taken into account. We address this by defining the null model of no-epistasis based on disjoint partitions of features *I* into two groups *I*_1_ and *I*_2_, which can be considered as two disconnected graphs. Features within each group (subgraph) can interact with each other via a decision tree model *CART_I__i_*, with *i* = 1, 2, but features between the two groups (subgraphs) cannot. The overall no-epistasis null model for the epiTree p-values then corresponds to the sum of the two individual trees, such that features between the two groups do not interact with each other. More precisely, for a dth-order interaction with *d* features *I* = {*A*_1_*,..., A_d_*}, we consider

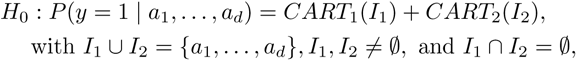

where *I*_1_ and *I*_2_ correspond to the partition which yield the highest prediction accuracy on the test data. We give details on the construction of p-values for higher-order interactions in the SI (see SI Section S1.1.9).

#### Epistasis (alternative) model

The epistasis (alternative) model is based on a single decision tree fit on training set samples and all features in an interaction. That is, the alternative model (no-epistasis) for a pairwise interaction is of the form

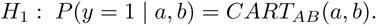

For a higher-order interaction of size d the alternative model corresponds in an analog way to a single decision tree taking all d features as input. E.g., for an interaction of size 4 with genes A,B,C, and D, the alternative model takes the form *H*_1_ : *P* (*y* =1 *| a, b, c, d*)= *CART_ABCD_*(*a, b, c, d*). See SI Section S1.1.9 for further details. An example from the red hair case study for fitted null and alternative models is shown in Fig 2 for the two genetic features A = *ASIP* and B = *DEF8*, see also the SI Fig S9 for response surfaces that show penetrance as a function of interacting genes.

**Fig 2.**
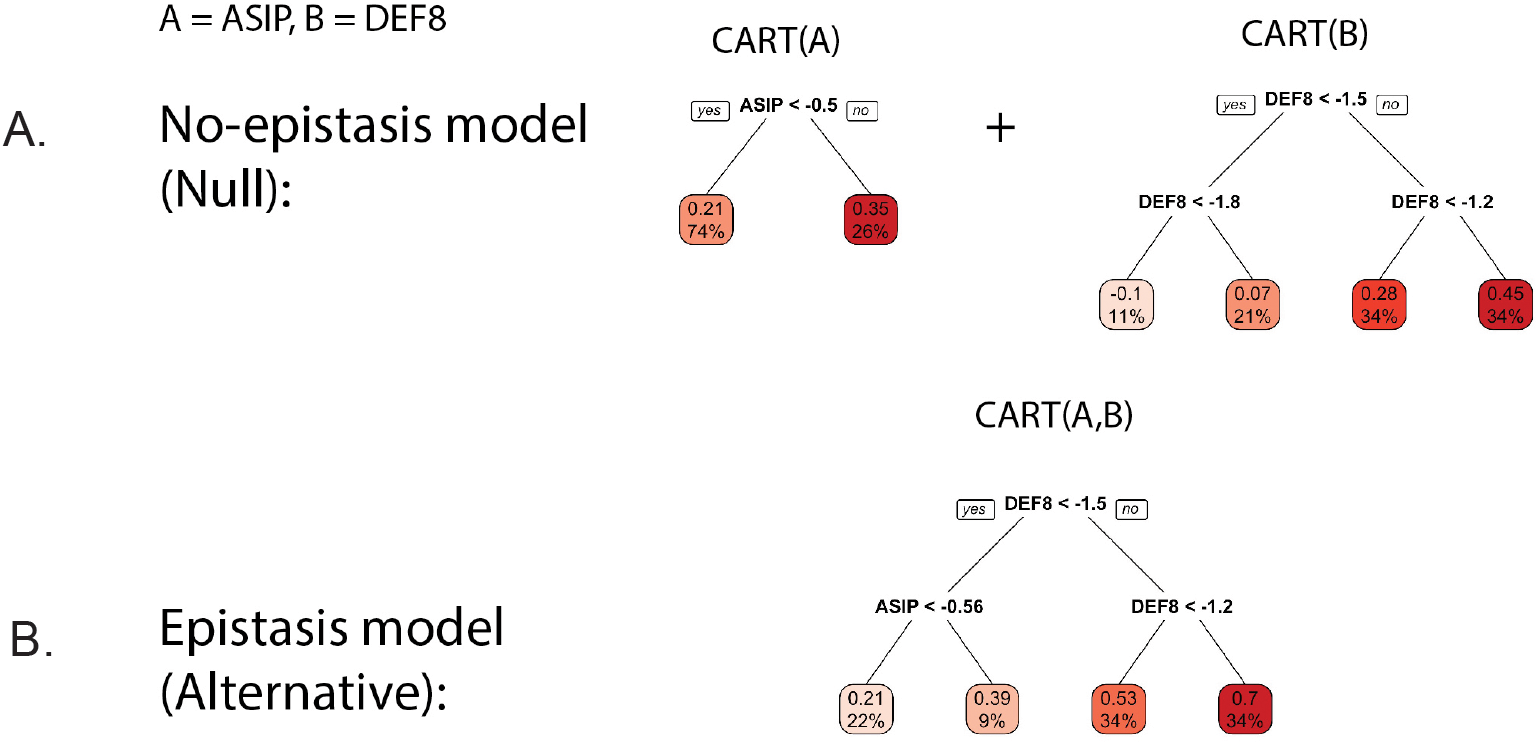
Example for the decision tree-based null model (no-epistasis) shown in top row (A.) and alternative model (epistasis) shown in bottom row (B.), for gene expression features A = *ASIP* and B = *DEF8*, which are tested for via the PCS p-values. The models where fitted using the CART algorithm [33] on the training data. The decimal digits at the tip nodes correspond to the predicted probability of red hair. The percentage at the tip nodes corresponds to the percentage of training observations falling into this tip node.

#### Tuning parameter selection

The CART algorithm, which is used to fit the models *CART_A_, CART_B_, CART_AB_* on the training data, depends on tuning parameters that influence the depth of these trees. We select these tuning parameters for each candidate interaction separately to ensure that the interaction model *CART_AB_* is as shallow as possible, while still capturing interactions between genes *A* and *B*—i.e., both genes *A* and *B* appear in the CART model. This depth constraint helps prevent overfitting, while guaranteeing that the complexity of all trees, *CART_AB_, CART_A_, CART_B_*, is comparable.

We note that our approach to tuning parameter selection results in different tuning parameters for each candidate interaction (i.e., decision tree model). If one selected a single (fixed) parameter for all interactions, then CART models would not necessarily contain all gene features involved in an interaction. More details on how we select the tuning parameters are given in SI Section S1.4, where we demonstrate that interactions with significant PCS p-values using shallow trees remain significant with deeper trees (i.e., our results are stable with respect to the tuning parameters of CART). Finally, we stress that by choosing shallow trees for the interaction model, our results also become more interpretable.

#### PCS inference

For an order-d interaction, PCS p-values are computed by comparing prediction error of the epistasis and the no-epistasis models. Prediction error is quantified using the log-likelihood, equivalent to cross-entropy for binary responses, and evaluated on hold-out test data. The epistasis and no-epistasis models are fitted on training data. When the epistasis model has a worse (or equal) prediction error compared to the no-epistasis model, we report no significant finding by formally setting the PCS p-value equal to one. That is, prediction on the test set is used as a screening as stipulated by PCS inference in [20]. Otherwise, we compute PCS p-values using a modified version of the classical likelihood ratio test, which is designed to take finite sample variability into account.

Specifically, for an interaction of size *d* with gene features *A*_1_*,..., A_d_*and binary responses *Y* (e.g., red hair yes/no), we consider *b* = 1*,..., B* bootstrap samples from the test datasets. We use the no-epistasis (i.e., additive tree) model, *f*_0_(*A*_1_*,..., A_d_*)= *CART*_1_(*I*_1_)+ *CART*_2_(*I*_2_), with *I*_1_ *[I*_2_ = {*A*_1_*,..., A_d_*}, to estimate each observation’s response probability *P*_0_*|_b_*. We then sample *null perturbation* responses *Y*_0_*|_b_ ∼* Bernoulli(*P*_0_*|_b_*). Thus, for each bootstrap sample we have a pair of vectors, *Y*_0_*|_b_* and *Y |_b_*, representing simulated (from the null perturbation) and true responses, respectively.

We use the likelihood ratio statistic,

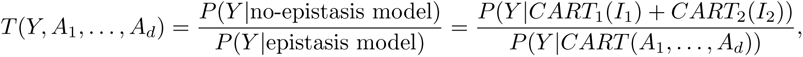

to quantify the improvement in prediction error of the epistasis model relative to the no-epistasis model. We extend the classical test for finite sample variability by asking whether the improvement, as quantified by *T*, is greater under the observed response compared with the null-perturbation response. That is, we evaluate *T* (*Y |_b_, A|_b_*) *> T* (*Y*_0_*|_b_, A|_b_*). The PCS p-value is given as

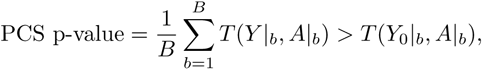

a stabilized version of the likelihood ratio p-value. We provide further details on the derivation of the PCS p-value in SI Section S1.1. In practice, one should choose B sufficiently large such that the PCS p-value converges. A simple derivation (see SI Section S1.1.8 for details), shows that it is easy to approximate the PCS p-value (conditioned on the data) as B tends to infinity, which we have done for the red hair analysis.

We call this new test the epiTree test. As a result, the quality of an interaction is directly related to its prediction accuracy on hold-out test data. A detailed description of underlying assumptions and derivation of the PCS p-value is presented in the Supplementary Information, see SI Section S1.1. We also provide an illustrative example for PCS p-values, which provides some additional intuition about PCS p-values in SI Section S1.6.

The results presented next are PCS epistasis p-values for gene level interactions found by iRF. We also present comparisons to the p-values obtained by a standard logistic regression analysis, see e.g., Figs S15 and S16 in the SI. We note that PCS inference could also be used to assess the significance of interaction terms in a logistic regression model, but we do not consider such an analysis in this paper.

## Results

### Red hair

To assess how well genotype/phenotype relationships learned by iRF (e.g., predicting ’red hair’ phenotype from the gene and variant features, respectively) generalized to new samples, we evaluated prediction accuracy on the hold-out test set. Fig 3 reports ROC curves for the prediction accuracy of iRF and competitors on the gene level (left) and variant level (right) (all fit using training data): penalized logistic regression with L1 penalty term and random forests (RF). With an area under the ROC curve (AUROC) of 0.93 (95% bootstrap confidence interval was [0.922, 0.936]) on hold-out test data, iRF demonstrates high prediction accuracy and outperforms its closest competitor (penalized logistic regression AUROC = 0.9, 95% bootstrap confidence interval of [0.894, 0.913]; ranger AUROC = 0.88, 95% bootstrap confidence interval of [0.871, 0.892]).

**Fig 3.**
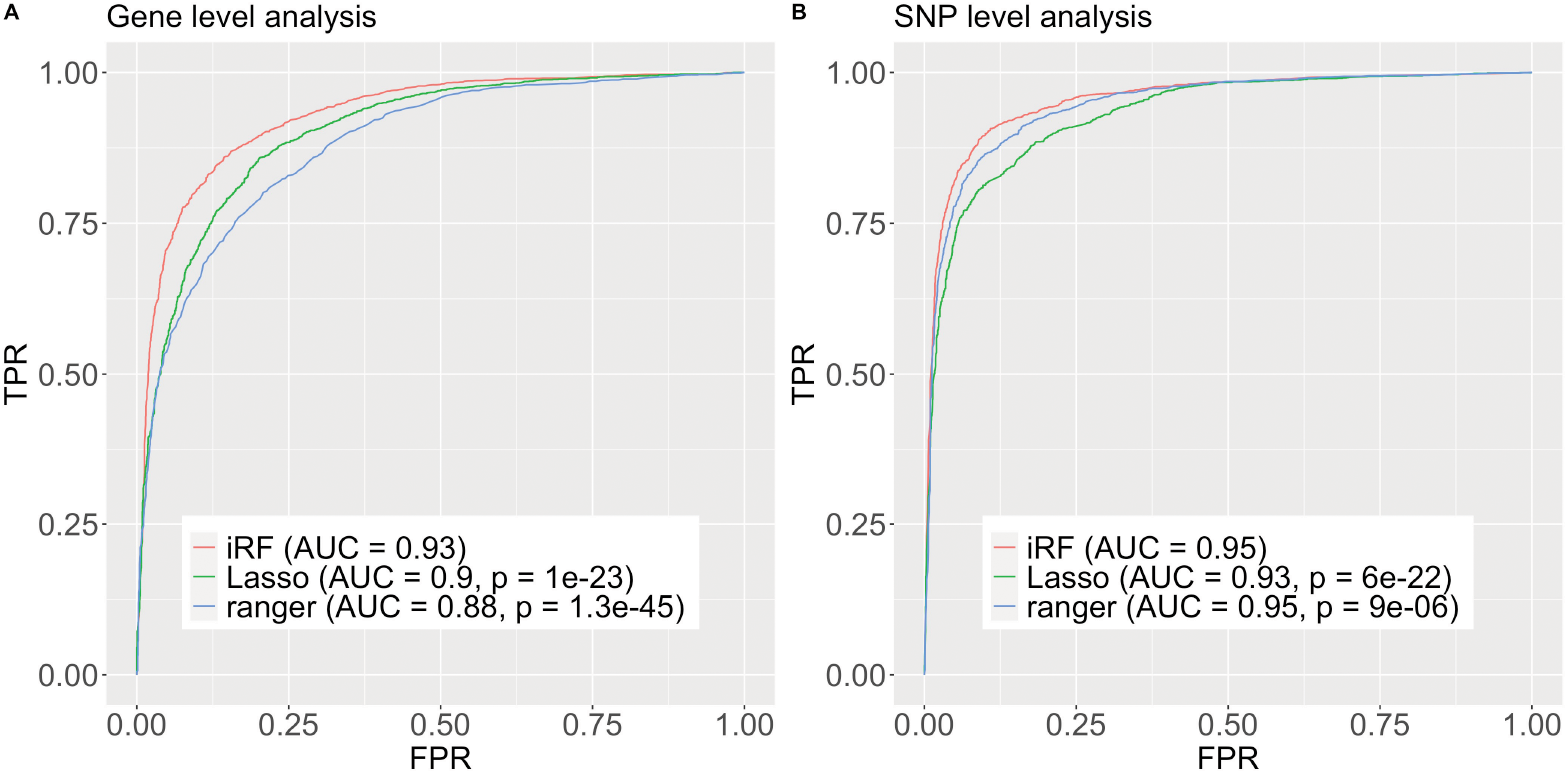
ROC curves of iRF prediction model and competitors on hold out test data. “Lasso” stands for a logistic regression model with an additive L1 penalty term on the parameter vector, i.e., a lasso type estimator. The lambda tuning parameter was selected via cross validation using the cv.glmnet R function from the glmnet R package. The “ranger” competitor corresponds to the random forest implementation of the R package ranger with default parameters. Left (A.): using the gene expression features to train a model which predicts ’red hair’. Right (B.): using the variant features to train a model which predicts ’red hair’. P-values testing for a difference between iRF’s ROC curve and the respective competitors are computed using the R package `pROC` and function `roc.test`, using DeLong’s Algorithm [63].

Before conducting inference on interactions recovered by iRF, we applied two filtering steps. First, we performed `n.bootstrap` = 50 bootstrap replicates of the iRF search and removed interactions appearing in fewer than 50% of these replicates. Second, we removed interactions among genes/variants that were in high linkage disequilibrium (LD) as it has been well-documented that LD can lead to spurious estimates of epistasis [19, 44, 45]. We note that many of the gene-level interactions we detected were among genes in the vicinity of MC1R (SI, Fig S23; LD *R*^2^ of at least 40% for all removed interactions); our LD filtering was equivalent to restricting results to interactions among genes/variants on different chromosomes. This resulted in 10 order-2, 6 order-3, and 2 order-4 interactions (gene-level analysis, Fig 5, see also Fig 4) as well as 25 order-2 and 3 order-3 interactions (SNP-level analysis, Fig 6), representing a massive reduction in the *∼* 10^16^ possible order-4 interactions among *∼* 10^4^ genes. See also SI Fig S23, which shows the number of discovered interactions per interaction-order within the individual screening steps. Interactions primarily centered around genes in the *MC1R* and *ASIP* regions, which corroborates previously reported epistatic interactions surrounding red hair, see [15]. We stress that while other works *a priori* restrict their search to interactions involving *MC1R*, our approach does not require such pre-selection.

**Fig 4.**
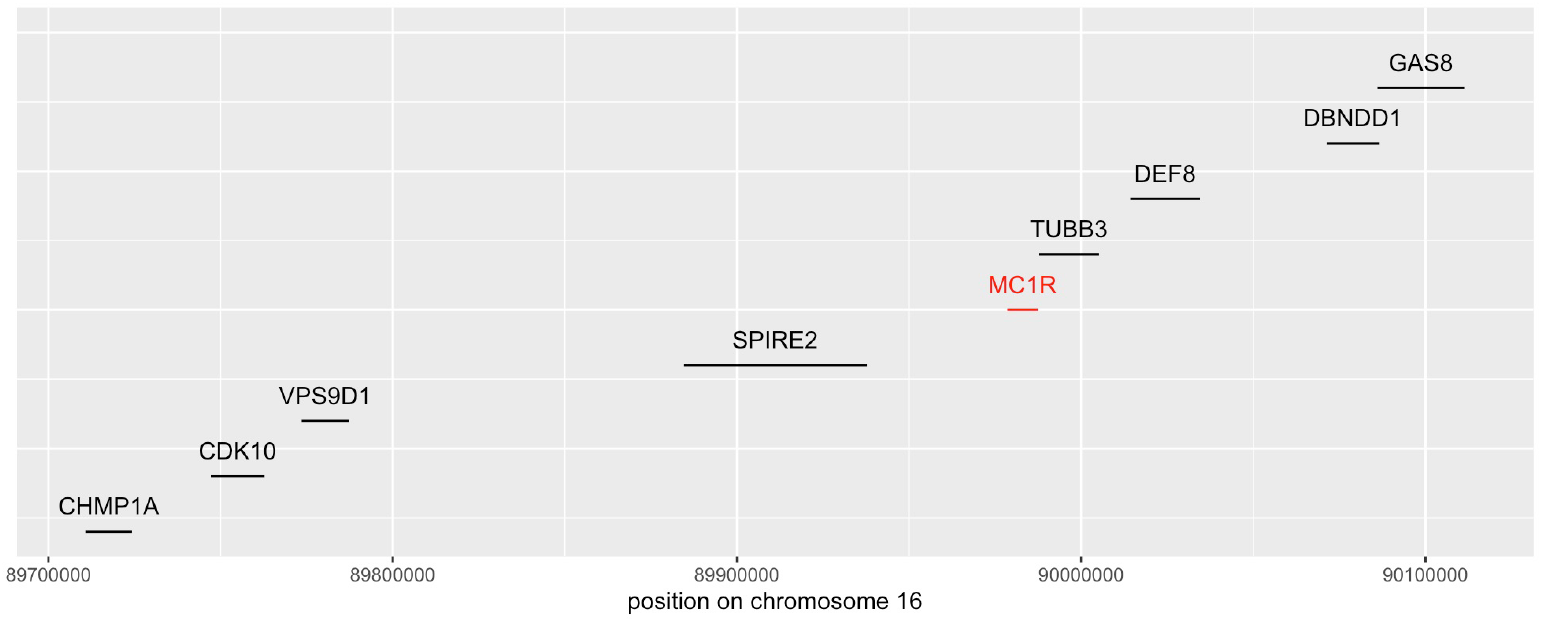
Location of the coding region for the 8 chromosome 16 genes which appear in the stable gene level interactions found by iRF, together with the location of the MC1R gene.

**Fig 5.**
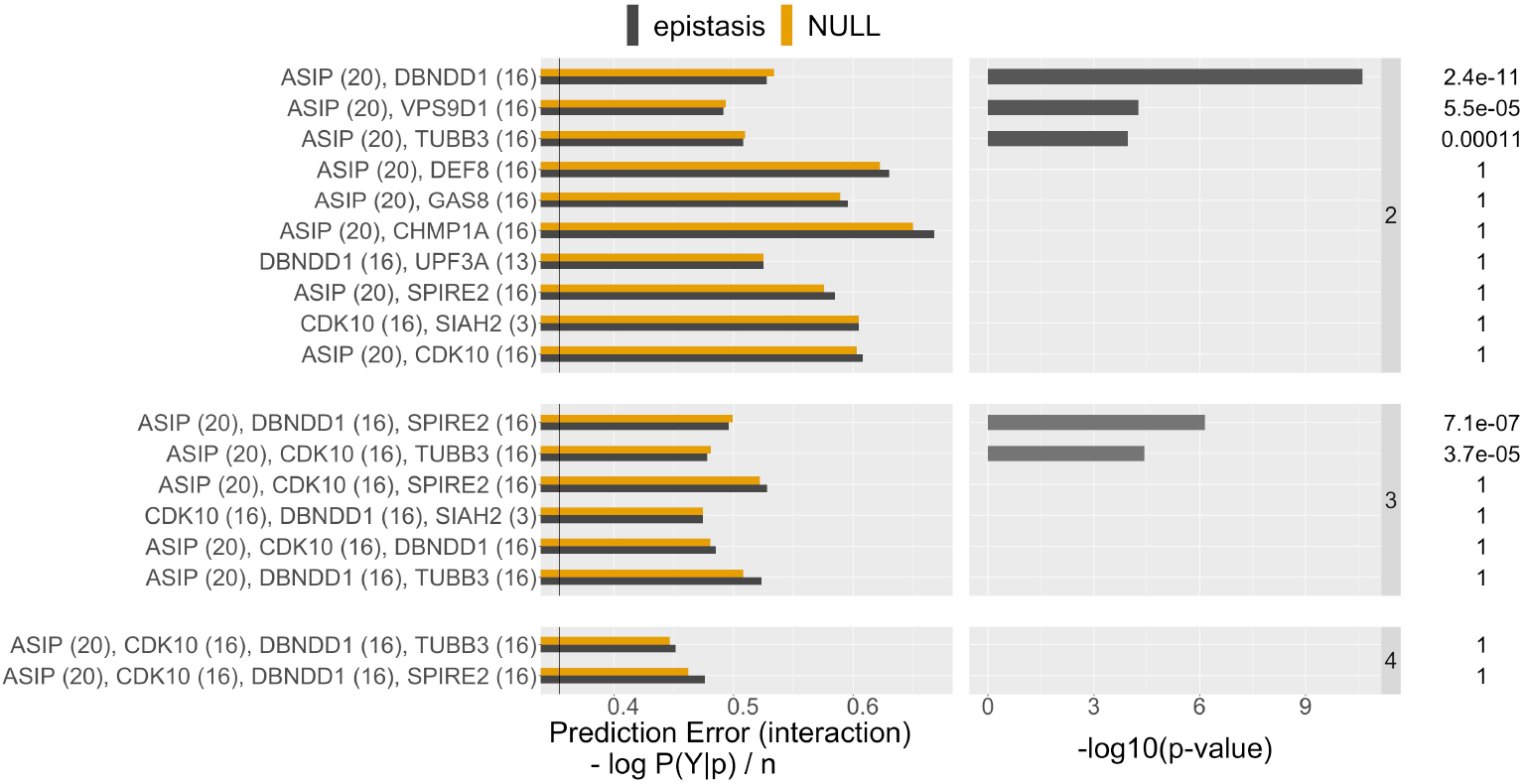
List of stable gene level interactions found by iRF (stability score *>* 0.5). The fist column shows the prediction error (defined as cross-entropy, see supplementary Section S1.1.6) on the test data of the learned CART models for both, no-epistasis (NULL, orange) and epistasis (alternative, gray). The second column shows the PCS p-value on a -log10 scale and the numeric value is shown on the very right, up to two significant digits. The black vertical line in the first column shows the prediction error achieved by iRF using all the gene features simultaneously. The chromosome of the respective gene is shown in parenthesis. The 8 genes on chromosome 16 all have their coding region in the vicinity of the MC1R gene, as shown in Fig 4. Note that the prediction error of the Null model can be less than the prediction error of the alternative model, as they are evaluated on hold-out test data. Whenever this happens the PCS p-values is 1 by construction.

**Fig 6.**
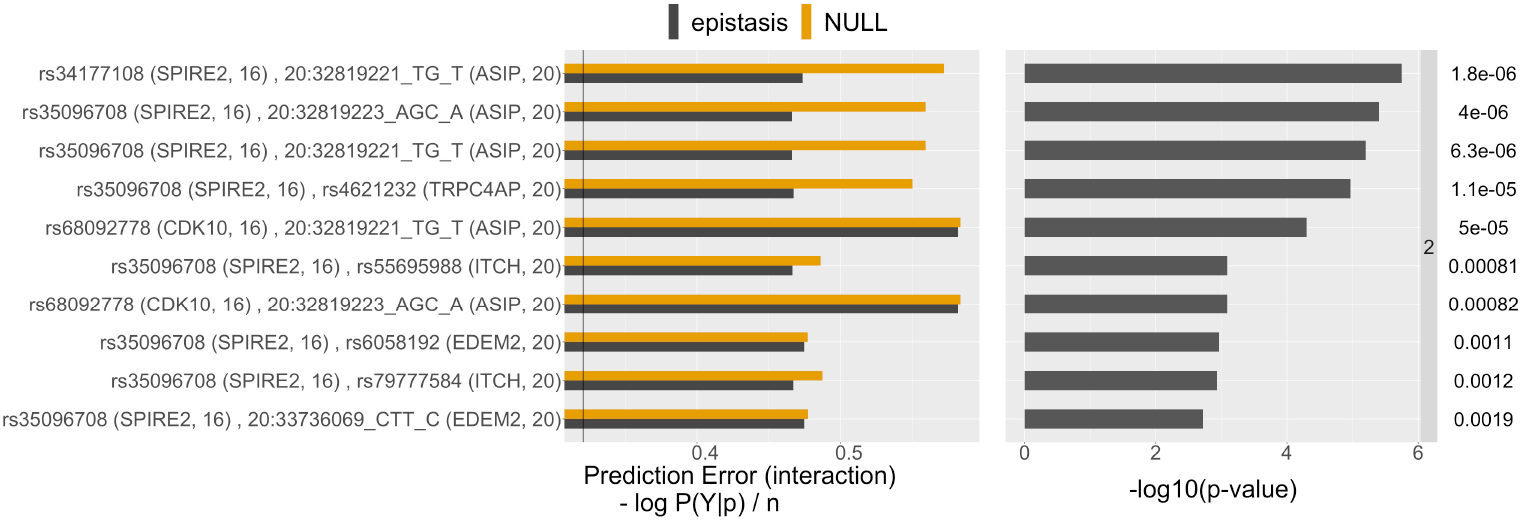
Same as Fig 5 for the top 10 order two variant level interactions.

We evaluated the significance of the individual candidate interactions using PCS epistasis inference (Figs. 5 and 6; comparisons with logistic regression reported in SI Figs S15-16). Interactions with significant PCS p-value (< 0.05 with Bonferroni correction) correspond to those for which the epistasis model results in a stable decrease in prediction error relative to the no epistasis model. For several of these interactions, CART tree-based models show a considerable improvement in prediction error relative to logistic regression (e.g. *ASIP*, *TUBB3* ; *ASIP*, *VPS9D1* ; Figs S15-16). In contrast, prediction accuracy of the epistasis model shows little improvement over the no epistasis model for interactions that are significant with respect to the logistic regression p-value. This finding suggests that the thresholding, Boolean form of interactions used by CART is a more accurate model of interaction behavior between genes comapred with the polynomial interaction model used by logistic regression. We hypothesize that this is a result of thresholding behavior that has been previously observed in biomolecular interactions [38–41].

### Multiple sclerosis

We further evaluated the epiTree pipeline in the context of multiple sclerosis (MS), a chronic, autoimmune disorder of the central nervous system. Since MS is known to be more prevalent in women [46], we trained separate ”Male” and ”Female” models both when searching for interactions and conducting inference. MS is believed to be influenced by both genetic and environmental factors [46], making it an inherently more complex phenotype to study compared with red hair. There are a range of genetic variants and biological processes which have been associated with the disorder [22–24, 47–51]. The most prominent ones correspond to genes in the MHC class II region, in particular surrounding HLA-DRB1, [26]. There exists evidence obtained from family data for epistatic interactions within the HLA-DRB1 region [25–27].

However, as noted above, for general cohort data as considered here, high linkage between variants / genes can result in spurious estimates of epistasis [19, 44, 45]. We are not aware of any results for epistatic interactions for the MS phenotype among genetically unlinked genes. In light of this complexity, we sought to determine whether epiTree could identify plausible interaction hypotheses among unlinked genes—surrounding variants and biological processes previously implicated in MS. Here, epiTree rankings can be viewed as narrowing the search space of possible interactions. We found that of the seven top-ranked interactions with non-significant (with Bonferroni correction) but small PCS p-value (< 0.1), five involve genes that have all been previously associated with MS or related processes. These five gene interactions are HLA-DRB1 + ZNF771 and HLA-DRB1 + SLC1A6 (in male subjects) and HLA-DRB1 + NDUFS2, HLA-DRB1 + STEAP3, and HLA-DRB5 + WDR18 + POC5 (in female subjects). Further details are provided in the ’Biological findings’ section below.

### PCS detects different types of epistasis

PCS epistasis and logistic regression p-values can differ substantially. This reflects the fact that each captures a different form of interaction, and thus, different genotype/phenotype relationships. The PCS p-values, based on CART models, operate directly on the penetrance scale and capture thresholding-type behavior. Logistic regression considers epistasis on a logit scale and is restricted to polynomial interaction terms. A key advantage of having CART as the building block for PCS p-values in our epiTree test is that interaction behavior is interpretable and easily visualized via the respective decision trees (Fig 2 for an example). We highlight two examples from the red hair case study below.

First, we considered interactions where PCS epistasis p-values were significant and logistic regression p-values were not. For example, the interaction between *ASIP* and *TUBB3* shows the greatest difference between PCS and classical p-values (see SI Fig S15 and Fig 5). Fig 7 reports response surfaces, indicating penetrance (*P* (red hair)) as a function of interacting features (top/bottom rows: logistic regression/CART, left/right columns: non-epistasis/epistasis models). For comparison, we report the smoothed distribution of test data as ground truth (Fig 7e). Here, the polynomial interaction term assumed by logistic regression fails to capture the non-monotic relationship between *TUBB3* and *P* (red hair). On the other hand, the CART model more accurately captures this relationship, which is confirmed by the improvement in prediction accuracy relative to logistic regression (SI Fig S15). In other words, the PCS p-value detects a non-additive, epistatic relation between the two genes that cannot be captured by the logistic regression model. Such non-monotonic behaviour, as it is observed in Fig 7, is common among PrediXcan estimated gene expression features.

**Fig 7.**
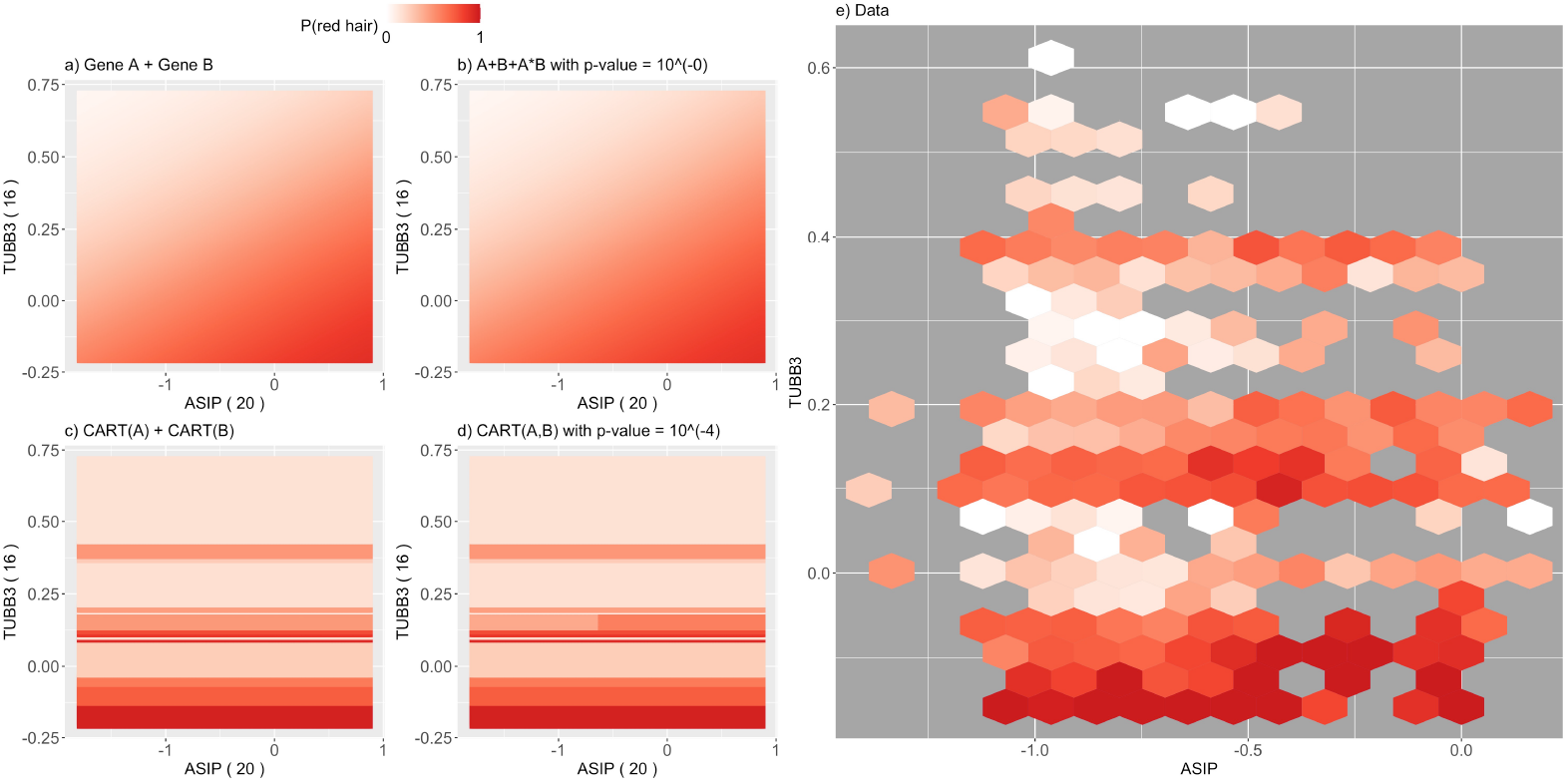
Response surface for *ASIP* - *TUBB3*, right plot (e.): smoothed test data is shown with colored hexagons providing percentage of red hair as a color code for the test data in specific hexagon (hexagons are drops when they contain less than 5 data points); left plot (a. - d.): response surfaces for fitted models; top (a./b.): logistic regression model, bottom (c./d.): CART based model, right (b./d.): epistasis model, left (a./c.): non-epistasis model.

This is due to the fact that estimated expression of a single gene corresponds to a weighted linear combination of discrete variants. When a small number of variants are strongly associated with the response and have large PrediXcan weights for a given gene, the response surface exhibits marked transitions that correspond to different values of the highly weighted variants. This is exactly the case for *TUBB3*, with 25% of its weight mass on the variant rs8048449, which shows strong marginal association with red hair. A similar example for the interaction *ASIP - DBNDD1* is shown in Fig 8.

**Fig 8.**
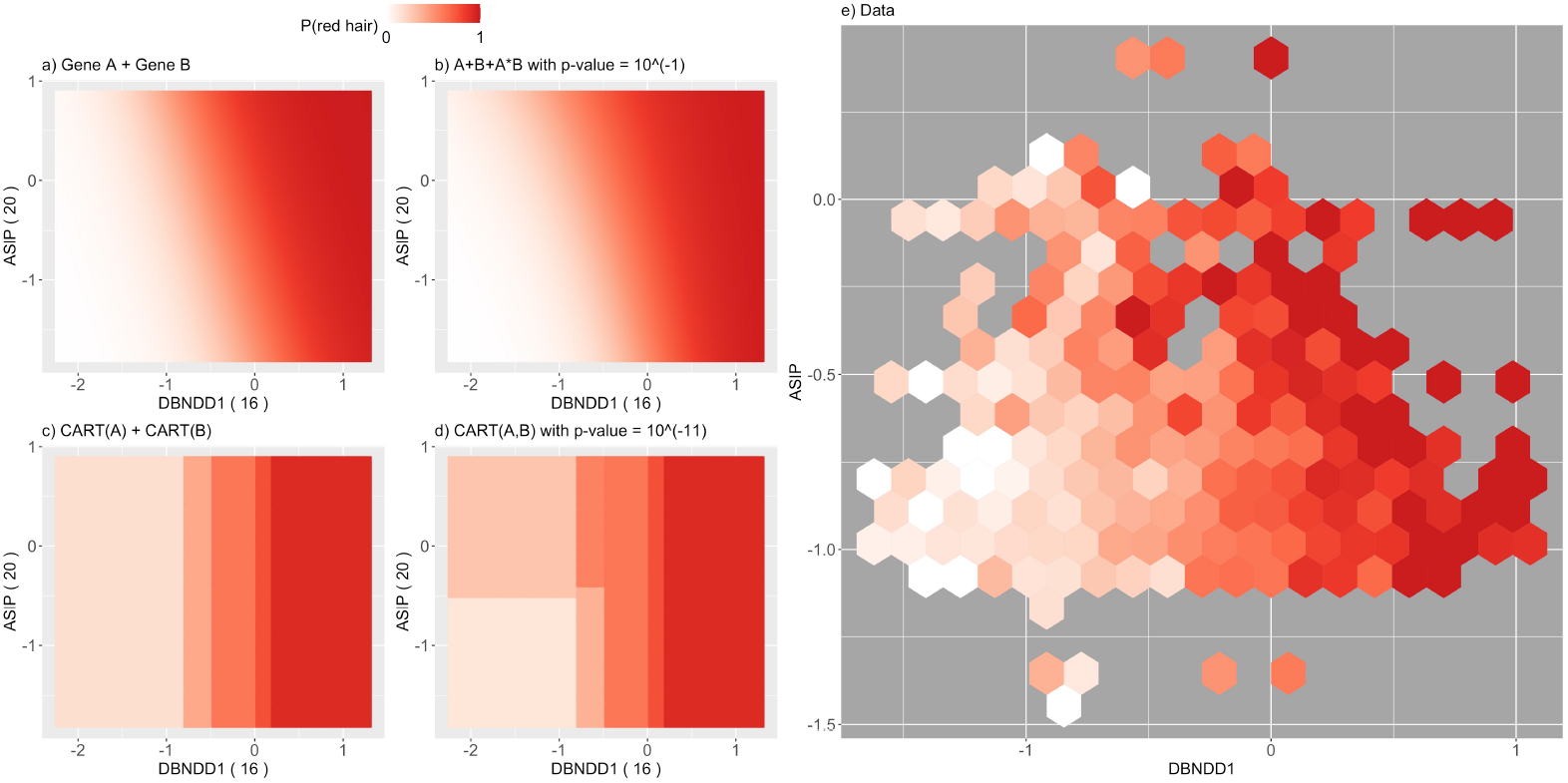
Response surface for *ASIP* - *DBNDD1*, otherwise as Fig 7

Second, we considered interactions where PCS epistasis p-values were not significant and logistic regression p-values were. For example, the interaction between *DEF8* and *ASIP* has highly significant logistic regression p-value (SI). The response surfaces indicate that these models differ most in the region where *DEF8* is small and *ASIP* is large. While the logistic regression model estimates a high probability of red hair in this region, there is limited data to support how responses behave here. Indeed, all four models (logistic regression non-epistasis, logistic regression epistasis, CART non-epistasis, CART epistasis) achieve nearly identical prediction accuracy (with average cross entropies of 0.624, 0.620, 0.622, 0.630, see SI Fig S15 and Fig 5). In other words, there is no evidence with respect to prediction accuracy that a particular model should be preferred, despite the significance of the classical logistic regression p-value. SI Fig S12 shows another example for the interaction *ZNF276-RPL36P4* where the PCS p-value is not significant but the logistic regression p-value is. Here, the data clearly shows a non-linear relationship in one of the variables, which cannot be captured by the linear components of the null model for logistic regression. CART can capture such a relation well and outperforms logistic regression with a simple additive relationship. That is, the logistic regression significant p-values appear to be driven by a main effect that is not captured by the linear form of main effect in logistic regression.

We provide a similar discussion for the remaining three pairwise interactions for which either the PCS p-value or the logistic regression p-value was significant (with Bonferroni correction) in SI Section S1.2.

## Biological findings

### EpiTree interactions recapitulate epistasis between *MC1R* and *ASIP*

For the red hair case study, the epiTree pipeline recovered several epistatic interactions between the *MC1R* and *ASIP* region, both on the gene and variant levels (Figs 5 and 6), that have previously been linked to red hair [15]. The *MC1R* gene on chromosome 16 is an evolutionarily conserved regulator of pigmentation with a strong association with the red hair phenotype in humans. As a result, other works, studying epistasis related to red-hair, restrict to interactions between variants in the *MC1R* region and other marginally associated regions as a pre-filtering step to reduce the overall search space for interactions, see [15]. We stress that our pipeline does not rely on any such *a priori* knowledge and still recovers the *MC1R* and *ASIP* interaction, making it particularly valuable for phenotypes that have not been studied extensively.

### EpiTree identifies novel higher-order interactions associated with red hair

The epiTree pipeline recovered several higher-order interactions, which is not computationally feasible in most other epistasis pipelines with a few exceptions where order three interactions are considered, e.g., in [52]. For example, the top order three interaction between *ASIP*, *DBNDD1*, and *SPIRE2* had significant PCS p-value with Bonferroni correction. Moreover, the respective CART interaction model showed a strong increase in prediction accuracy, compared to the additive CART model (no-epistasis), as well as both logistic regression models (epistasis and non-epistasis). We note that *DBNDD1* and *SPIRE2* are both genes on chromosome 16 which are just 200 kb apart. Thus, in this case, we cannot rule out the effect of strong genetic linkage which may result in spurious detection of epistasis.

### EpiTree suggests interactions among genes not previously associated with red hair

In addition to recapitulating well established epistatic interactions between *MC1R* and *ASIP*, our pipeline epiTree also provides new insights for the red hair phenotype. The iRF model selection step identified two genes that were not previously associated with red hair: *UPF3A* and *SIAH2*, both involved in interactions with genes neighboring *MC1R* (*DBNDD1* + *UPF3A* and *CDK10* + *SIAH2*). However, neither the PCS p-values nor the logistic regression p-values were significant for these interactions and also the subsequent iRF variant interaction filtering did not result in any epistasis related to these genes. Consequently, we cannot report those interactions as significant findings. Nonetheless, we do see some further indication for potential association of these genes with red hair. In the pairwise gene level search of the top iRF genes, both, *UPF3A* and *SIAH2*, appear among the top interactions with respect to logistic regression p-value, with logistic regression p-value < 0.01, see SI Fig S20. We note however, that with Bonferroni correction the respective p-values are not significant. Moreover, we note that the PCS p-value, while providing comparable or even higher prediction accuracy, was not significant for the same interactions. In summary, with the current data at hand, we cannot report any strong statistical evidence for the association of *UPF3A* and *SIAH2* with red hair, but these are suggestive of further investigations with new data.

### EpiTree identifies evidence of epistasis among genes previously associated with red hair

For the pairwise, inter-chromosome gene-level analysis of the top 50 iRF genes, we found that an interaction between *PKHD1* (chr 6) and *XPOTP1* (chr 20)

was among the top interactions in terms of PCS p-value, with the CART model from PCS also showing some improvement in terms of prediction accuracy compared to the logistic regression models. The respective response surface and interaction CART model, shown in SI Figs S13 and S14, suggests a thresholding of gene expression levels with the red hair phenotype. In particular, one observes a decrease in red hair penetrance when gene expression of both, *PKHD1* and *XPOTP1*, are small and an increase when both are large. Otherwise, when one is small and the other is large, one observes an average red hair penetrance of around 50% (recall that we consider a balanced sample in this analysis). This interaction behavior is well described by the tree-based model used in the PCS p-value and results in a PCS p-value of *∼* 10*^-^*^3^, although we note that this p-value is not significant after Bonferroni correction for all pairs of genes that were tested and thus, we cannot report this interaction as a statistically significant finding. We note that both, *PKHD1* and *XPOTP1*, have been associated with hair color previously. In particular, in the recent work [15] both genes had reported epistasis with *MC1R*. However, epistasis between the two genes has not been previously described.

### Top-ranked interactions associated with MS surround *HLA-DRB1*

In the MS case study, iRF models obtained AUROC of 0.58 and 0.63 for male and female samples respectively, see SI Fig S24. As in the red hair case study, we considered stable interactions (stability *>* 0.5) with genes from different chromosomes to avoid spurious interactions from high linkage. iRF detected 209 and 60 such interactions for male and female subjects, respectively. After applying the epiTree test to these interactions, we found that most of them had PCS p-value equal to 1 and none of them were significant after Bonferroni correction. However, for the male subjects we found that 2 and for the female subjects 5 interactions had a PCS p-value < 0.1 (without Bonferroni correction). These interactions are displayed in SI Fig S25. In both male and female samples, all of these interactions involved HLA-DRB1, which has been associated with MS across multiple populations [22–24], or HLA-DRB5, which is in high LD with HLA-DRB1. Of the two interactions surrounding HLD-DRB1 detected in males, one involves ZNF771—a gene that has been associated with male MS in transcriptomic meta analysis [47], while the other involves SLC1A6—a gene involved in regulation of the neurotransmitter glutamate [53], which is present in excess levels in MS patients [48]. In females, 3 / 5 interactions surrounding HLA-DRB1 involve genes that are either explicitly associated with MS or involved in processes known to be dysregulated in MS. NDUFS2, which is involved in Mitochondiral Complex I, has been previously associated with MS [51]. STEAP and WDR18 are involved in ion transport and RNA metabolism, processes that have been previously reported as dysregulated in MS [49,50], while mutations in POC5 have been associated with increased BMI [50], a known risk factor for MS [54]. While there is no gold standard for genetic interactions associated with MS, these results point to interesting hypotheses—the majority of which involve genes that are individually linked to MS and/or processes believed to be involved in its etiology.

### PCS epistasis p-values detect diverse interaction forms in simulation studies

To evaluate the performance of epistasis inference methods, we conducted a series of data-based simulations comparing (i) epiTree PCS epistasis p-values (ii) logistic regression p-values (iii) permutation random forest (pRF) p-values [55]. We generated responses using four different models, each corresponding to a different form of interaction, applied to the gene expression data from the red hair case study. For each response generating model, we generated a set of candidate interactions using iRF and computed p-values for each inference method. For three of the four response generating models, epiTree both consistently recovered true interaction(s) (along with subsets of interacting features) and ranked these interactions as most significant across all candidates. In contrast, pRF consistently recovered true interactions in 2/4 response generating models. However, pRF tended to rank any candidate interaction that included the true interaction as highly significant (even if the candidate included inactive features). Finally, logistic regression recovered the true interaction in 2/4 response generating models, though at a lower significance level and with greater variability. For a detailed description of our simulation study and results, see SI Section S1.7.

## Discussion

### Summary of findings

Here we introduce the epiTree pipeline, building on a novel tree-based method and PCS framework, to detect epistasis. epiTree is grounded in a biologically relevant dimensional-reduction schema derived from tissue-specific gene expression to derive variant-level interactions, and additionally, may detect interactions of between more than two loci. A particular advantage of our methodology is that it allow for flexible forms of Boolean interactions and additive components. For many interactions, we find that this flexibility provides better fit to the data compared with multiplicative terms used in logistic regression. Moreover, by modeling responses directly on the penetrance scale, our approach avoids the need for transforming responses as, e.g., performed in logistic regression.

Without prior knowledge of the phenotype of red-hair coloration, our approach detects known interactions between *MC1R* (and nearby genes related to pigmentation) and *ASIP*, as well as novel interactions between *CDK10* and *SIAH2*. Our findings may suggest a previously unrecognized relationship between *SIAH2* and the red hair phenotype. Interestingly, *SIAH2* is known to be associated with breast cancer risk and cellular response to hypoxia -a phenotype that has been associated with melanoma and pigmentation in general [56]. The red hair phenotype, which was considered in this manuscript, mainly served as an illustrative example for the epiTree pipeline. We selected this phenotype because of its well established epistasis that provides a positive control for interaction discovery. We stress that for further validation regarding new epistasis findings for the red hair phenotype independent data will be needed. We also consider another case study for the more complex phenotype of multiple sclerosis (MS). While epiTree detects interactions among genes related to processes dysregulated in MS, we are not aware of previously reported findings on gene-level interactions among unlinked genes associated with MS to benchmark against.

### Related work

Previous approaches typically screen variants based on main effects due to computational constraints [15, 52, 57, 58], which may miss interacting genetic variants with weak main effects. The tree-based approach presented here does not limit analysis to variants with a strong main effect, however we note that decision tree-based methods have been proposed before in epistasis, e.g., [52, 57, 59, 60]. In [57] and [52] they are used to obtain marginal feature importance for the individual variant features from which candidate interactions are extracted either via a brute force search or with the help of phasing, in particular, independent of the phenotype under consideration. In contrast, going beyond main effect genes, iRF explicitly exploits the structure of the trees in a forest to extract Boolean interactions by interpreting frequently co-occurring features along the paths of trees as interactions. Similar to iRF, [59] and [60] take the joint appearance of features on a decision path as a measure of interaction. [59] interpret every path in a collection of boosted decision trees as a candidate interaction. To ensure the set of reported interactions is a manageable size, the authors restrict their search to interactions among at most d variants (with d = 5 being the default value). In contrast, iRF employs the stability principle [20, 61]—reporting feature combinations that frequently co-occur rather than the entire set of decision paths—to filter the set of candidate interactions with no restriction on size, after soft-dimensionality reduction through iterations of RF. [60], on the other hand, extract a measure of interaction strength from an RF for a pre-selected interaction. Moreover, their approach does not provide a way to extract candidate interactions from the trees as is done in iRF via the RIT algorithm; thus, it also requires a brute force search in practice. We are not aware of any other tree-based method for epistasis that incorporates stability driven bootstrap sub-sampling into its pipeline for screening interactions. More importantly, all these previous works do not develop a significance test or p-value that is based on tree models to evaluate the found interactions.

### Limitations and directions for future work

Our application of iRF is based upon biologically-relevant dimensional reduction in the form of estimated tissue-specific expression values derived from GTeX via PrediXcan [29]. While this approach detects both known and previously undescribed epistasis, it depends on the accuracy of gene expression estimates derived from bulk RNA sequencing on heterogeneous cell types. We make the assumption that tissue-specific data is sufficiently representative of the biology from which the phenotype of interest is derived. This assumption of representativeness is not likely to hold for traits deriving from rare cell types or from tissues and cell-types, which are not sampled in GTeX in the first place. Additionally, we note that the number of individuals in GTeX is not currently representative of all population strata, which serves to limit the application of our approach for detection of epistasis to human phenotypes by excluding diverse genetic populations. Finally, our use of PrediXcan for dimensionality reduction models epistatic relationships at the level of transcription, which may miss variants that are not associated with changes in gene expression levels. Nevertheless, our epiTree test is a stand-alone test for epistasis of any given interaction relative to a phenotype that arises from other knowledge rather than the PrediXcan.

In principle, the epiTree pipeline can detect interactions of arbitrary order. However, the maximal detectable interaction order depends on tree depth—for the iRF screening step an interaction corresponds to subsets of features on individual decision paths. In general, the depths of a decision tree learned from data scales logarithmically with the number of samples. This implies that, in general, we cannot detect interactions of order higher than *O*(ln(*n*)). From a practical perspective, this limitation is fairly minor as even order-3 interactions remain unexplored in many biological settings. A corollary of this limitation is that the number of samples falling into an individual leaf node splitting on *d* features is *O*(*n/*2*^d^*) (assuming balanced splits), where *n* is the sample size. Thus for an interaction represented by a single leaf node, *n/*2*^d^* needs to be large enough to detect deviation from additivity. Thus, detection of higher-order interactions becomes exponentially more difficult as the size of the interaction increases. This problem applies not just to the epiTree pipeline, but to interaction discovery in general, where the size of all possible candidate interactions of order *d* grows exponentially. However, while traditional methods that screen through all possible interactions are typically restricted to pairwise interaction due to computational reasons, the epiTree pipeline can detect interactions of higher order if the interaction signal is strong enough. More mathematical details on how the order of an interaction enters the complexity of the interaction discovery problem for Boolean interaction recovery with a theoretically tractable version of iRF can be found in [37].

Another potential limitation of the iRF interaction screening is that it does not explicitly take the genomic location of individual features into account. Even on the gene expression level, features that are close-by on the genome are typically highly correlated. Such features might appear exchangeably in interactions, as observed for the red hair phenotype with *MC1R*, see Fig 5. More generally, it is well known that for random forest based algorithms, correlated features can lead to masking effects, where the effect of a feature might be hidden by another dominating, correlated feature [62]. Finally, as discussed earlier, genetic features that are close-by on the genome might show artificial epistasis that is caused by hidden linkage with unobserved causal variants. In fact, for the red hair phenotype iRF reported various interactions among genes in the vicitinity of MC1R (recall Results section) that we filtered out subsequently. One idea to overcome these limitations, would be to analyze by haplotypes of merged close-by genes into a single hyper-feature. However, two correlated genetic features do not necessarily show the same epistatic behaviour. Thus, there is a general tradeoff. Aggregating features in this way may increase the ability to detect epistasis by concentrating the forest on a single feature. On the other hand, aggregation may lead to loss of specificity that decreases the chance of detecting epistasis. For the red hair analysis, all interactions were either among genes between different chromosomes, in which case there is no correlation or linkage between them, or among genes that were all in the vicinity of MC1R and contained variants that were highly linked to each other (recall Results Section). Therefore, we followed the most simple approach for the red hair analysis and filtered out interactions on the same chromosome. In general, we note that more advanced approaches are possible by taking drop-off of linkage explicitly into account.

Although, the CART tree-based model that we propose for the PCS p-value (see Materials and Method and Supplementary Information) can describe significantly more flexible additive and non-additive relationships for the epistasis and non-epistasis model than a single linear or multiplicative term, we also emphasize that not every functional relationship might be well described by a relatively simple CART model. In particular, when a relationship is truly linear or multiplicative, clearly, working with a linear model will be more powerful than a CART approximation. The response surface plots, as shown in e.g., in Fig 7 and discussed in the Results section, are a powerful tool to investigate whether CART or linear models might be more appropriate for the given data at hand. Moreover, we stress that PCS p-values are, in principle, not restricted to CART models, but could be combined with any learning algorithm that appears appropriate for the data. As with any machine learning algorithms there is typically a tradeoff between interpretability and prediction accuracy. We find the CART tree-based models provide a good balance between both, being sufficiently flexible and very straight-forward to interpret, while capturing the thresholding behavior of interactions among many biomolecules.

The epiTree pipeline involves several screening steps in order to result in a set of candidate interactions, such as estimating gene expression features from the SNP data, applying iRF, and fitting individual tree-based null and alternative models. We stress, however, that all these screening steps are based on the training data only, while the final evaluation of the PCS p-values is based on the hold-out test data. With this sample-split approach, which is well accepted as a post-selection tool, we obtain valid inference that avoids overfitting from the screening procedure.

Here, we obtained our training and test data sets from a random sample split. While such an approach results in valid inference, we recognize that using an independent or external data set as test data will be preferred. At the same time, we also recognize that in many cases such an independent data set will not be available (also for our analysis an independent data set for the red hair phenotype was not available to us). Therefore, our PCS p-values are designed in such a way that they explore the full data distribution via bootstrap sampling of the test-data, which makes them more stable to the somewhat arbitrary choice of the random sample split.

The sample splitting paradigm that we follow in our approach holds the advantage of separating the discovery from the inference stage, which prevents overfitting and thus, makes findings more reliable, in general. We note, however, that compared to classical p-values, the need of a separate training set, will generally imply that there is less data available to evaluate the p-values, thus, a decrease in power. However, we note that due to the iRF interaction selection step, in general our approach requires the testing of orders of magnitude fewer hypotheses than any brute-force p-value approach. Thus, multiplicity correction of p-values will be much less of a problem in our pipeline, i.e., less stringent significance thresholds for p-values apply, which can compensate for this effect.

## Conclusion

Our new methodology, the epiTree pipeline, provides an approach to identify predictive and stable Boolean, epistatic genetic relationships. epiTree goes well beyond multiplicative interaction, providing flexibility to represent a more diverse set of biological interactions, and is capable of detecting predictive and stable genetic interactions of order three and higher. The two case studies substantiate the promise of epiTree for epistatic interaction discovery, especially for complex phenotypes such as MS, where epiTree ranking provides a useful source of info to reduce design space for follow-up experiments

## Acknowledgments

This work was carried out under UK Biobank study number 15860.

## Data and code availability

The datasets and code generated during this study are available at https://github.com/merlebehr/epiTree. The raw genetic and phenotype data are available from UK Biobank.

## References

1. Bateson W. Mendel’s Principles of Heredity. Cambridge Univ. Press; 1909.

2. Ritchie MD. Finding the Epistasis Needles in the Genome-Wide Haystack. In: Epistasis. Methods in Molecular Biology (Methods and Protocols). vol. 1253. New York: Humana Press; 2015. p. 19–33.

3. Bell JT, Timpson NJ, Rayner NW, Zeggini E, Frayling TM, Hattersley AT, et al. Genome-Wide Association Scan Allowing for Epistasis in Type 2 Diabetes: 2D GWA Scan of Type 2 Diabetes. Annals of Human Genetics. 2011;75(1):10–19.

4. Van Steen K, Moore JH. How to Increase Our Belief in Discovered Statistical Interactions via Large-Scale Association Studies? Human Genetics. 2019;138(4):293–305.

5. Nag A, McCarthy MI, Mahajan A. Large-Scale Analyses Provide No Evidence for Gene-Gene Interactions Influencing Type 2 Diabetes Risk. Diabetes. 2020;69(11):2518–2522.

6. Fisher RA. The Correlation between Relatives on the Supposition of Mendelian Inheritance. Transactions of the Royal Society of Edinburgh. 1919;52(2):399–433.

7. Wade MJ, Winther RG, Agrawal AF, Goodnight CJ. Alternative Definitions of Epistasis: Dependence and Interaction. Trends in Ecology & Evolution. 2001;16(9):498–504.

8. Cordell HJ. Epistasis: What It Means, What It Doesn’t Mean, and Statistical Methods to Detect It in Humans. Human Molecular Genetics. 2002;11(20):2463–2468.

9. North BV, Curtis D, Sham PC. Application of Logistic Regression to Case-Control Association Studies Involving Two Causative Loci. Human Heredity. 2005;59(2):79–87.

10. Phillips PC. Epistasis—the Essential Role of Gene Interactions in the Structure and Evolution of Genetic Systems. Nature reviews Genetics. 2008;9(11):855–867.

11. Sailer ZR, Harms MJ. Detecting High-Order Epistasis in Nonlinear Genotype-Phenotype Maps. Genetics. 2017;205(3):1079–1088.

12. Wu X, Dong H, Luo L, Zhu Y, Peng G, Reveille JD, et al. A Novel Statistic for Genome-Wide Interaction Analysis. PLoS Genetics. 2010;6(9):e1001131.

13. Ueki M, Cordell HJ. Improved Statistics for Genome-Wide Interaction Analysis. PLOS Genetics. 2012;8(4):e1002625.

14. Huang Y, Wuchty S, Przytycka TM. eQTL Epistasis – Challenges and Computational Approaches. Frontiers in Genetics. 2013;4:51.

15. Morgan MD, Pairo-Castineira E, Rawlik K, Canela-Xandri O, Rees J, Sims D, et al. Genome-Wide Study of Hair Colour in UK Biobank Explains Most of the SNP Heritability. Nature Communications. 2018;9:5271.

16. Wasserstein RL, Lazar NA. The ASA Statement on P-Values: Context, Process, and Purpose. The American Statistician. 2016;70(2):129–133.

17. McShane BB, Gal D, Gelman A, Robert C, Tackett JL. Abandon Statistical Significance. The American Statistician. 2019;73(sup1):235–245.

18. Kim K. Massive False-Positive Gene–Gene Interactions by Rothman’s Additive Model. Annals of the Rheumatic Diseases. 2019;78(3):437–439.

19. de los Campos G, Sorensen DA, Toro MA. Imperfect Linkage Disequilibrium Generates Phantom Epistasis (and Perils of Big Data). G3: Genes, Genomes, Genetics. 2019;9(5):1429–1436.

20. Yu B, Kumbier K. Veridical Data Science. Proceedings of the National Academy of Sciences. 2020;117(8):3920–3929.

21. Santosh Bangalore S, Wang J, Allison DB. How Accurate Are the Extremely Small -Values Used in Genomic Research: An Evaluation of Numerical Libraries. Computational Statistics & Data Analysis. 2009;53(7):2446–2452.

22. Alcina A, Abad-Grau MdM, Fedetz M, Izquierdo G, Lucas M, Fernandez O, et al. Multiple sclerosis risk variant HLA-DRB1* 1501 associates with high expression of DRB1 gene in different human populations. PloS one. 2012;7(1):e29819.

23. Fogdell A, Hillert J, Sachs C, Olerup O. The multiple sclerosis-and narcolepsy-associated HLA class II haplotype includes the DRB5* 0101 allele. Tissue antigens. 1995;46(4):333–336.

24. McElroy JP, Cree BA, Caillier SJ, Gregersen PK, Herbert J, Khan OA, et al. Refining the association of MHC with multiple sclerosis in African Americans. Human molecular genetics. 2010;19(15):3080–3088.

25. Gregersen JW, Kranc KR, Ke X, Svendsen P, Madsen LS, Thomsen AR, et al. Functional epistasis on a common MHC haplotype associated with multiple sclerosis. Nature. 2006;443(7111):574–577. doi:10.1038/nature05133.

26. Lincoln MR, Ramagopalan SV, Chao MJ, Herrera BM, DeLuca GC, Orton SM, et al. Epistasis among *HLA-DRB1, HLA-DQA1*, and *HLA-DQB1* loci determines multiple sclerosis susceptibility. Proceedings of the National Academy of Sciences. 2009;106(18):7542–7547. doi:10.1073/pnas.0812664106.

27. Ramagopalan SV, Julian C K, George C E. Multiple sclerosis and the major histocompatibility complex. Current opinion in neurology. 2009;22(3):219–225.

28. Bycroft C, Freeman C, Petkova D, Band G, Elliott LT, Sharp K, et al. The UK Biobank Resource with Deep Phenotyping and Genomic Data. Nature. 2018;562(7726):203–209.

29. Gamazon ER, Wheeler HE, Shah KP, Mozaffari SV, Aquino-Michaels K, Carroll RJ, et al. A Gene-Based Association Method for Mapping Traits Using Reference Transcriptome Data. Nature Genetics. 2015;47(9):1091–1098.

30. Lonsdale J, Thomas J, Salvatore M, Phillips R, Lo E, Shad S, et al. The Genotype-Tissue Expression (GTEx) Project. Nature Genetics. 2013;45(6):580–585.

31. Basu S, Kumbier K, Brown JB, Yu B. Iterative Random Forests to Discover Predictive and Stable High-Order Interactions. Proceedings of the National Academy of Sciences. 2018;115(8):1943–1948.

32. Kumbier K, Basu S, Brown JB, Celniker S, Yu B. Refining Interaction Search through Signed Iterative Random Forests. bioRxiv:467498. 2018;.

33. Breiman L. Random Forests. Machine Learning. 2001;45:5–32.

34. Shah RD, Meinshausen N. Random Intersection Trees. The Journal of Machine Learning Research. 2014;15(1):629–654.

35. Wang Q, Tang TM, Youlton N, Weldy CS, Kenney AM, Ronen O, et al. Epistasis regulates genetic control of cardiac hypertrophy. medRxiv. 2023;doi:10.1101/2023.11.06.23297858.

36. Cliff A, Romero J, Kainer D, Walker A, Furches A, Jacobson D. A High-Performance Computing Implementation of Iterative Random Forest for the Creation of Predictive Expression Networks. Genes. 2019;10(12):996. doi:10.3390/genes10120996.

37. Behr M, Wang Y, Li X, Yu B. Provable Boolean interaction recovery from tree ensemble obtained via random forests. Proceedings of the National Academy of Sciences. 2022;119(22):e2118636119. doi:10.1073/pnas.2118636119.

38. Little JW, Shepley DP, Wert DW. Robustness of a Gene Regulatory Circuit. The EMBO Journal. 1999;18(15):4299–4307.

39. Kobiler O, Rokney A, Friedman N, Court DL, Stavans J, Oppenheim AB. Quantitative Kinetic Analysis of the Bacteriophage Genetic Network. Proceedings of the National Academy of Sciences. 2005;102(12):4470–4475.

40. Little JW. Threshold Effects in Gene Regulation: When Some Is Not Enough. Proceedings of the National Academy of Sciences. 2005;102(15):5310–5311.

41. Levine E, Hwa T. Small RNAs Establish Gene Expression Thresholds. Current Opinion in Microbiology. 2008;11(6):574–579.

42. Cordell HJ, Todd JA, Hill NJ, Lord CJ, Lyons PA, Peterson LB, et al. Statistical Modeling of Interlocus Interactions in a Complex Disease: Rejection of the Multiplicative Model of Epistasis in Type 1 Diabetes. Genetics. 2001;158(1):357–367.

43. Breiman L, Friedman JH, Stone CJ, Olshen RA. Classification and Regression Trees. New York: Chapman and Hall; 1984.

44. Wood AR, Tuke MA, Nalls MA, Hernandez DG, Bandinelli S, Singleton AB, et al. Another Explanation for Apparent Epistasis. Nature. 2014;514(7520):E3–E5.

45. Zan Y, Forsberg SKG, Carlborg Ö. On the Relationship between High-Order Linkage Disequilibrium and Epistasis. G3: Genes, Genomes, Genetics. 2018;8(8):2817–2824.

46. Goldenberg MM. Multiple sclerosis review. Pharmacy and therapeutics. 2012;37(3):175.

47. Català-Senent JF, Andreu Z, Hidalgo MR, Soler-Sáez I, Roig FJ, Yanguas-Casás N, et al. A deep transcriptome meta-analysis reveals sex differences in multiple sclerosis. Neurobiology of Disease. 2023;181:106113.

48. Levite M. Glutamate, T cells and multiple sclerosis. Journal of Neural Transmission. 2017;124(7):775–798.

49. Williams R, Buchheit CL, Berman NE, LeVine SM. Pathogenic implications of iron accumulation in multiple sclerosis. Journal of neurochemistry. 2012;120(1):7–25.

50. Hecker M, Ruege A, Putscher E, Boxberger N, Rommer PS, Fitzner B, et al. Aberrant expression of alternative splicing variants in multiple sclerosis–A systematic review. Autoimmunity reviews. 2019;18(7):721–732.

51. Ban M, Elson J, Walton A, Turnbull D, Compston A, Chinnery P, et al. Investigation of the role of mitochondrial DNA in multiple sclerosis susceptibility. PLoS One. 2008;3(8):e2891.

52. Jiang R, Tang W, Wu X, Fu W. A Random Forest Approach to the Detection of Epistatic Interactions in Case-Control Studies. BMC Bioinformatics. 2009;10(Suppl 1):S65.

53. Martinez-Lozada Z, Guillem AM, Robinson MB. Transcriptional regulation of glutamate transporters: from extracellular signals to transcription factors. Advances in pharmacology. 2016;76:103–145.

54. Gianfrancesco MA, Glymour MM, Walter S, Rhead B, Shao X, Shen L, et al. Causal effect of genetic variants associated with body mass index on multiple sclerosis susceptibility. American journal of epidemiology. 2017;185(3):162–171.

55. Li J, Malley JD, Andrew AS, Karagas MR, Moore JH. Detecting gene-gene interactions using a permutation-based random forest method. BioData mining. 2016;9(1):1–17.

56. Bedogni B, Powell MB. Hypoxia, Melanocytes and Melanoma - Survival and Tumor Development in the Permissive Microenvironment of the Skin. Pigment Cell & Melanoma Research. 2009;22(2):166–174.

57. Chen X, Liu CT, Zhang M, Zhang H. A Forest-Based Approach to Identifying Gene and Gene–Gene Interactions. Proceedings of the National Academy of Sciences of the United States of America. 2007;104(49):19199–19203.

58. Cordell HJ. Detecting Gene–Gene Interactions That Underlie Human Diseases. Nature Reviews Genetics. 2009;10(6):392–404.

59. Wan X, Yang C, Yang Q, Xue H, Fan X, Tang NLS, et al. BOOST: A Fast Approach to Detecting Gene-Gene Interactions in Genome-Wide Case-Control Studies. The American Journal of Human Genetics. 2010;87(3):325–340.

60. Yoshida M, Koike A. SNPInterForest: A New Method for Detecting Epistatic Interactions. BMC Bioinformatics. 2011;12:469.

61. Yu B. Stability. Bernoulli. 2013;19(4):1484–1500.

62. Louppe G. Understanding Random Forests: From Theory to Practice. arXiv:14077502. 2015;.

63. Sun X, Xu W. Fast Implementation of DeLong’s Algorithm for Comparing the Areas Under Correlated Receiver Operating Characteristic Curves. IEEE Signal Processing Letters. 2014;21(11):1389–1393. doi:10.1109/LSP.2014.2337313.

