## Supplementary Information for "Learning epistatic polygenic phenotypes with Boolean interactions"

### Supplementary Information to manuscript
### Learning epistatic polygenic phenotypes with Boolean interactions

**S2 Supplemental Figures** **33**

### S1 Supplemental Material and Methods

#### S1.1 epiTree test based on PCS epistasis inference and PCS p-value

Traditional methods of statistical hypothesis testing evaluate uncertainty relative to a null distribution that describes the hypothesized data generating process. Intuitively, these methods address questions of the form: how likely is the observed test statistic if data were drawn from the specified null distribution. A statistic (and hence data) that is not likely under the null provides evidence against the corresponding null model. In practice, null distributions are often used without justification as to why they are a "reasonable" baseline for such comparisons. This can lead to small p-values that provide strong evidence against a "straw-man" null hypothesis — i.e. a null hypothesis that is *a priori* known or widely believed to be a poor description of the data generating process.

To address this issue, the PCS inference framework [34] introduced a prediction screening step that evaluates model accuracy on hold-out test set. This step allows one to filter out "straw-man" null hypotheses and focus inference on those that provide a reasonable fit to the data. In order to evaluate the accuracy of red hair phenotype predictions, we split data into training and hold-out test sets, learn models corresponding to epistasis and non-epistasis on the training set, and carry out statistical hypothesis testing on the hold-out test set. By splitting our data in this way, we introduce an additional layer of uncertainty — inference is defined relative to one of many possible sample splits. We address this extra layer of uncertainty through the PCS inference framework.

Specifically, we build on general PCS inference ideas to assess the strength of an epistatic effect. Intuitively, our inference framework captures both the uncertainty quantified by traditional statistical inference and the uncertainty surrounding data perturbations (sample split). In other words, the PCS p-values we propose consider not only distributional assumptions of a null model — as in traditional statistical inference — but also uncertainty surrounding the range of sample splits that could have been considered but were not. By "inflating" p-values to account for a wider range of uncertainty, our approach helps ensure that evidence for a given interaction is more robust to a broader range of modeling decisions. We call our approach epiTree test, because it is for **epi**stasis discovery and uses decision-**tree**s for both epistasis and no-epistasis models.

##### S1.1.1 Inference setting

For simplicity, we assume that a candidate interaction consists of two features: A and B. We provide details for the analogous case of general higher-order interactions in Section S1.1.9. Features A and B could be either continuous-valued gene expression features or discrete-valued SNP features. In the following, we focus on the gene expression setting. The case of SNP features is similar and described in Section S1.1.10. Recall that in our epiTree pipeline, candidate interactions are obtained by running iRF on the training data. The PCS epistasis inference for these candidate interactions, as outlined below, uses the same training data to obtain individual (for each interaction), tree-based null (no-epistasis) and alternative (epistasis) models. On the separate hold-out test data, our aim is to evaluate evidence (i.e. a p-value) for the null hypothesis

$$H_0 : (A, B) \text{ is not epistatic for phenotype } Y, \tag{1}$$

relative to the alternative hypothesis

$$H_1 : (A, B) \text{ is epistatic for phenotype } Y, \tag{2}$$

using test data consisting of $n$ sample points $Y = (y_1, \ldots, y_n)^\top$, $A = (a_1, \ldots, a_n)^\top$, $B = (b_1, \ldots, b_n)^\top$. The individual response values $y_i$ and features $a_i, b_i$ may be continuous valued or discrete. Here, we consider the binary phenotype of red hair, $y_i \in \{0, 1\}$, where $y_i = 1$ indicates red hair, and continuous-valued gene expression features $a_i, b_i \in \mathbb{R}$.

##### S1.1.2 Recap of standard logistic regression

The classical translation of Fisher's original definition of epistasis for binary phenotypes seeks evidence against null $H_0$ relative to alternative hypothesis $H_1$ under a simple logistic regression model. That is, the data are assumed to have come from a logistic regression model, which is a strong assumption that often leads to unrealistic uncertainty assessments in practice, especially for large samples of data. Models for the null and alternative can be written as,

$$\begin{aligned} H_0 &: \text{logit}(P(y = 1|a, b)) = \beta_A a + \beta_B b + \beta_0, \\ H_1 &: \text{logit}(P(y = 1|a, b)) = \beta_A a + \beta_B b + \beta_{AB} ab + \beta_0, \end{aligned} \tag{3}$$

where $\beta_j$, $j \in \{A, B, AB, 0\}$ represent coefficients describing the contribution of $A$, $B$, the interaction term $AB$, and an intercept respectively. In some cases, correction for population structure is necessary. Employing standard chi-square approximations for the likelihood ratio test of $H_0$ vs. $H_1$, it is straightforward to obtain a p-value for the inference problem in (3). This (or some slight variation) is the standard p-value typically reported as a measure of an interaction's significance, e.g., with p-value $< 0.05$/number of interactions tested, reported as significant (with Bonferroni correction).

##### S1.1.3 Scaling of the response variable

The logit scaling in (3) has a considerable influence on the way an interaction is defined. Clearly, a relation which is multiplicative (non-linear, hence epistatic) on one scale, can be additive (non-epistatic) in another. For instance, a log transform of a multiplicative interaction becomes additive. Hence, Fisher's commonly accepted definition of *epistasis* critically depends on the selected scaling. In fact, the situation is even more extreme. It is demonstrated in [11] that any multivariate, real valued function with compact support can be written as an additive function for some appropriate scaling. In other words, without fixing a response scale, every function can be rewritten in an additive form. Thus, Fisher's definition of epistasis is only well-defined relative to a selected scale.

The intrinsic scaling problem for epistasis has been pointed out by many authors, see e.g., [6, 7, 24, 26]. Out of statistical convenience, logistic regression models still dominate standard analyses. However, there is no biological justification for the logistic scaling. It has even been reported in empirical studies that this scaling does not always reflect biological function [24, 10].

Acknowledging that any epistatic results is only well-defined with respect to a specific scaling, it is natural (see e.g., discussion and references in [6]) to select a canonical scaling for this purpose: the penetrance $P(y = 1 \mid a, b)$ itself (instead of a logit scaling that transforms the penetrance via the function $f(x) = log(x/(1-x))$ as in logistic regression) [12, 7]. We articulate Fisher's definition in the penetrance scale as

$$H_0 : P(y = 1 \mid a, b) = f_A(a) + f_B(b) \quad \text{vs.} \quad H_1 : P(y = 1 \mid a, b) = f_{AB}(a, b), \tag{4}$$

where $f_j : \mathbb{R} \to [0, 1]$, $j \in \{A, B, AB\}$ are functions that describe the relationship between genes or interactions and penetrance.

###### S1.1.4 Form of additive and interaction models using training data

While raw gene expression data are often represented as counts (e.g. RNA-Sequencing), commonly available data are typically preprocessed. For example, PrediXcan estimates/imputes inverse normal transformed gene expression rather than raw expression value. In practice, there are a range of standard transforms and pre-processing steps, such that the actual scaling of gene features $A$ and $B$ can be rather arbitrary (see e.g., [2] for some general discussion). These transforms/pre-processing steps are generally not linear, although they are typically monotonic. As a result, we favor models that are invariant to monotone transformations of the data (resulting from potential pre-processing steps). In addition, we want to allow more flexible mappings beyond linear (as in 3) to account for non-linearities that pervade biological systems. There are many different forms of non-linear functional relationships, which may each be useful in describing different epistatic behavior. A particularly flexible class of functions are decision trees, which have the benefit of being both simple to interpret and invariant to monotone feature transformations. In addition, as mentioned previously, it is well known that many biological processes exhibit thresholding dynamics that are mimicked in the Boolean rules used by decision trees

[21, 16, 20, 18]. Moreover, for the classical epistasis model $H_1$ in (3) the multiplicative functional form lacks biological justification. Collectively, these considerations motivate our use of decision trees in both the additive, null model $H_0$ and non-additive, alternative model $H_1$. All individual decision trees are obtained via a CART [4] fit on the training data, using backfitting for the additive model. In summary, we translate Fisher's epistasis definition into a (non-parametric) hypothesis inference setting as

$$H_0 : P(y = 1 \mid a, b) = CART_A(a) + CART_B(b) \quad \text{vs.} \quad H_1 : P(y = 1 \mid a, b) = CART_{AB}(a, b). \tag{5}$$

Decision trees $CART_A, CART_B, CART_{AB}$ are fit on the training data set (that is, the same data that was used in the iRF candidate selection step) via the CART algorithm (using the R Package rpart). To fit the additive regression model $H_0$ in (5), we applied backfitting [13], where one recursively re-fitts additive CART components on the respective residuals with the remaining components. We stopped the recursion when none of the predicted values changed by more than 1% compared to the previous iteration. We further discuss the parameter choice for the CART algorithm in Section S1.4. Finally, we conduct inference using a hold-out test set, following the prediction principle (see next section).

###### S1.1.5 Predictability principle from the PCS framework

Classical p-values, as obtained via logistic regression-tests for model (3), compare the goodness-of-fit for the null and the alternative hypothesis using all available data. In other words, the null and the alternative models are fit on the same data that are used to evaluate the quality of each fit. In contrast, we learn null and alternative models on one (often randomly sampled) training data set and compare their prediction accuracy on a disjoint (often randomly sampled) set of test or hold-out data. This sample splitting approach is an old idea in statistics and has been used recently to deal with post-selection and so called universal inference problems, see, e.g., [31, 17, 3, 9, 32] and is widely used by the machine learning community to guard against overfitting.

Fitted null and alternative models correspond to hypotheses for non-epistasis (null, additive) and epistasis (alternative, non-additive) respectively, which leads to a simple hypothesis testing problem on the hold-out test data (since we fix both, the null and the alternative model from training data). We define predictions from the non-epistasis and epistasis models on the test data with gene expression (or SNP) values $(a_i, b_i)$, for $i = 1, \ldots, n$, as:

$$p_0(a_i, b_i) = CART_A(a_i) + CART_B(b_i) \quad \text{vs.} \quad p_1(a_i, b_i) = CART_{AB}(a_i, b_i). \tag{6}$$

For simplicity, we denote $p_0, p_1 \in \mathbb{R}^n$ as the $n$-vectors with components $p_{0,i} := p_0(a_i, b_i)$ for and $p_{1,i} := p_1(a_i, b_i)$.

In principle, one could apply a Neyman-Pearson test (with likelihood ratio test statistic) in order to obtain a p-value (conditioned on $p_0$ and $p_1$). However, the models $p_0$ and $p_1$ will generally be (at least slightly) mis-specified. For large sample size this can lead to unrealistic extremely small p-values (e.g., in the red hair data example as small as $10^{-100}$). This astronomically small p-value is due to the fact that the chi-squared asymptotic distributional approximation to the likelihood ratio test-statistics does not take into account model misspecification, which cannot be ignored for sufficiently large sample sizes (as in the UK biobank data).

In the following we use the PCS inference ideas proposed in [34] to explicitly take finite sample variability into account, via a bootstrapping approach on the test data, and to address the misspecification problem through prediction error evaluation on the test set. Instead of using the simple null hypothesis of the Neyman-Pearson test — the underlying penetrance is exactly $p_0$ — we use a more flexible null hypothesis. Specifically, we inflate the $p_0$ null distribution with the (centered) empirical sample distribution of the test statistic (via bootstrap sampling). As a result, our approach to inference takes into account the empirical variation in the likelihood ratio test statistic on the test data.

###### S1.1.6 epiTree test details: computation of CART based PCS p-value with bootstrap sampling of test data

Classically, p-values in a logistic regression model are calculated using a chi-squared approximation of the likelihood ratio test statistic under the null distribution, derived without accounting for model misspecification. Our approach does not assume a classical probabilistic model. Instead, it generates a null hypothesis based on the PCS framework, where the null distribution is approximated directly via bootstrap samples of the test data. To conduct inference using this approach, we need to specify a test statistic and null distribution, which we detail in the following two paragraphs.

**Test statistic.** CART models (5) estimate penetrance for both the no-epistasis (null) and epistasis (alternative) hypotheses (using training data). Given this pair of fitted CART models, we estimate penetrance in the test data as $p_0$ (no-epistasis) and $p_1$ (epistasis) based on test set genotype features. Note that both, $p_0$ and $p_1$, are independent of the observed response $Y$ in the test data used for inference. Given the observed test data $(Y, A, B)$, we consider a canonical test statistic, the log-likelihood ratio

$$T(Y) = T(Y, A, B) = \ln\left(\frac{P(Y \mid p_0(A, B))}{P(Y \mid p_1(A, B))}\right), \tag{7}$$

with

$$P(Y|p_\cdot) = \prod_{i=1}^{n} p_{\cdot i}^{y_i}(1 - p_{\cdot i})^{1-y_i},$$

where $p_\cdot$ can be $p_0$ or $p_1$, and we have dropped dependence on $A, B$ in our notation for simplicity. For given labels $Y$, $T(Y)$ measures how much more likely observed labels are under the null ($p_0$) compared to the alternative ($p_1$). Smaller values of $T(Y)$ correspond to greater evidence for the epistasis model.

**PCS Prediction screening.** Note that both, $p_0$ and $p_1$, were obtained from training data only. Evaluating $P(Y \mid p_0)$ and $P(Y \mid p_1)$ on hold-out test data provides a measure of how accurately the null and alternative models generalize to new samples. When $T(Y) \geq 0$, or equivalently $P(Y \mid p_0) \geq P(Y \mid p_1)$, then the alternative model $p_1$ (epistasis) provides no increase in prediction accuracy — as measured by the likelihood — on the test data compared to the null model $p_0$ (no epistasis). In this case, we conclude that there is not evidence for epistasis and epiTree formally reports a p-value of 1, that is

$$\text{PCS p-value} = 1 \quad \text{if } P(Y \mid p_0) \geq P(Y \mid p_1). \tag{8}$$

When $T(Y) < 1$, i.e. the epistasis model yields an increase in prediction accuracy, we quantify the uncertainty surrounding this improvement by pairing a simulated null data perturbation with bootstrap sub-sampling, as outlined below.

**PCS Null perturbation.** The PCS framework uses *null perturbations* to simulate data that respect known structure and compares observed results to results obtained from these simulated data. Recall that $p_0$ in (6) denotes our estimated penetrance for the test data under the hypothesis of no epistatic interaction. We simulate responses under the no epistasis null hypothesis as

$$Y_0 \sim \text{Bernoulli}(p_0), \tag{9}$$

which we refer to as the null perturbation. Below, we outline our approach for constructing a reference null distribution of the test statistic.

**PCS p-value** To obtain a PCS p-value, we use bootstrap re-sampling over the test data to evaluate

$$\text{PCS p-value} = P_{\text{bootstrap}}(T(Y) > T(Y_0)), \tag{10}$$

where $P_{\text{bootstrap}}$ indicates randomness relative to bootstrap sampling of the test data. More precisely, we generate bootstrap samples $m = 1, \ldots, M$, each of size $n$ (= sample size of the test data) drawn i.i.d. with replacement from the empirical distribution of our observed test data. For each bootstrap sample we obtain $p_0|m \in [0,1]^n$, a vector of estimated penetrance under the null. Using $p_0|m$, we simulate null responses for bootstrap sample $m$ under the null perturbation (9) as

$$Y_0|m \sim \text{Bernoulli}(p_0|m).$$

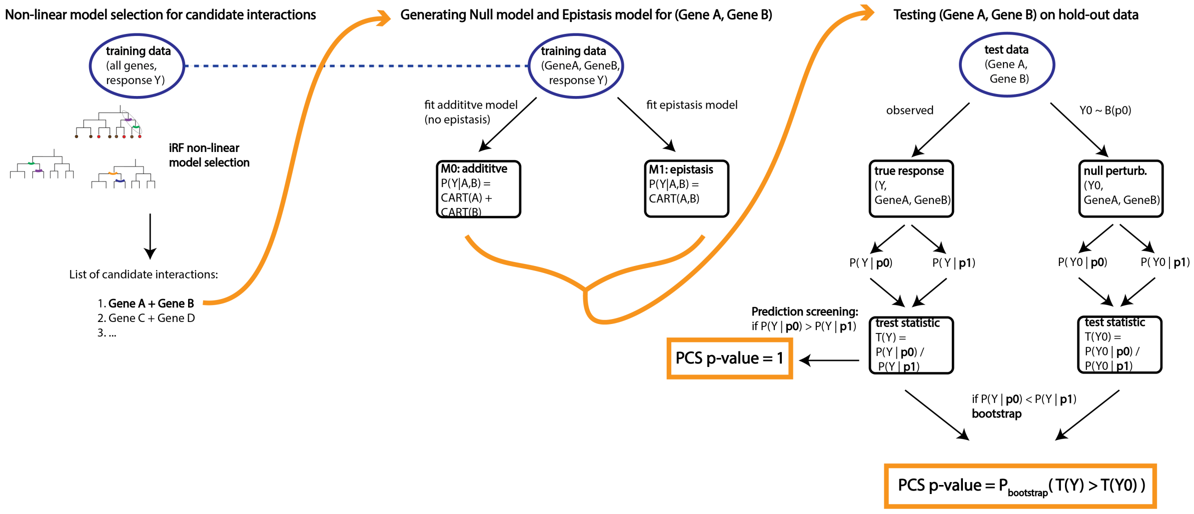

Figure S1: Illustration of epiTree pipeline.

| | CART based PCS p-value | logistic regression p-value |
|---|---|---|
| Scale for response | penetrance scale $P(Y \mid A, B)$ | Logit scale: $\log\left(\frac{P(Y \mid A,B)}{1-P(Y \mid A,B)}\right)$ |
| Null model (no epistasis) | $CART(A) + CART(B)$ | $\beta_A A + \beta_B B + \beta_0$ |
| Alternativ model (epistasis) | $CART(A, B)$ | $\beta_A A + \beta_B B + \beta_{AB} AB + \beta_0$ |
| Data used for model fitting | (separate) training data | full data |
| P-value calculation | Bootstrap from null perturbation | Distributional approximation (chi-square) |

Table S1: High-level summary of comparison between p-value from logistic regression with polynomial model vs. PCS p-value with CART model for epistasis testing.

In addition to the null responses above, we obtain $Y|m \in \{0,1\}^n$, an $n$-vector consisting of the observed responses for the $m$th bootstrap sample. The PCS p-value is then given by

$$\text{PCS p-value} = \frac{1}{M} \sum_{m=1}^{M} \mathbb{1}\{T(Y|m) > T(Y_0|m)\}. \tag{11}$$

In practice, one can obtain a simple analytic approximation of (11) as outlined in Section S1.1.8. By construction, $Y$ and $Y_0$ have the same distribution under the null hypothesis, making our proposed PCS inference a valid p-value (conditioned on $p_0$). We summarize the general test procedure epiTree in terms of the computation of the CART based PCS p-values in Figure S1. Table S1 shows a high level comparison of the CART based PCS p-values and p-values obtained from logistic regression.

##### S1.1.7 Comparison between PCS p-value and classical p-value calculations

In the following, we provide insights into the major differences between calculations of a PCS p-value and a classical p-value. For a given test statistic $T(Y)$, null hypothesis $H_0 : Y \sim F_0$ with null distribution $F_0$, and alternative hypothesis $H_1$ where smaller values of $T$ correspond to stronger evidence against the null (as e.g., for $T$ as in (7) for the epiTree test), a classical p-value is given by

$$\text{p-value } = P_{F_0}(T(Y_0) < t), \tag{12}$$

where $t = T(Y)$ is the observed test statistic and the probability is taken over the randomness of $Y_0$ with distribution $F_0$. The classical p-value in 12 does not take into account empirical fluctuation of $T(Y)$ among different subsamples of data — randomness only enters through $Y_0$.

In contrast, the PCS p-value explicitly takes the empirical variability of the test data $(Y, A, B)$ into account through bootstrap sampling, as in (11). This is necessary because sample splitting leads to a particular test set. Bootstrap samples of the test set mimic the other possible test sets that could have been resulted from a different sample split. More precisely, let $\hat{F}_n$ denote the empirical distribution of the test data. As the number of bootstrap samples $M$ in (11) tends to infinity,

$$\text{PCS p-value } = P_{F_0 \times \hat{F}_n}(T(Y_0) < T(Y)). \tag{13}$$

Thus, asymptotically, when the bootstrap distribution (empirical distribution) converges to the true underlying distribution of $Y$, denoted as $F$, the PCS p-value is equivalent to

$$\text{PCS p-value } \approx P_{F_0 \times F}(T(Y_0) < T(Y)), \tag{14}$$

where $Y$ and $Y_0$ are independent. For the given observed value $t$, as in (12), we can rewrite (14) to obtain

$$\text{PCS p-value } \approx P_{F_0 \times F}(T(Y_0) + (t - T(Y)) < t), \tag{15}$$

where $t$ is fixed and the probability is taken as in (14) jointly over $(Y_0, Y)$ with distribution $F_0 \times F$. Note that the difference between (12) and (15) is that $T(Y_0)$ is replaced by $T(Y_0) + (t - T(Y))$. This means that the effective null distribution of the PCS p-value corresponds to a convolution of the proposed null distribution, $T(Y_0)$, and a centered version of the observed distribution $T(Y)$. In particular, (15) does not just take into account the distribution of the probabilistic null model $F_0$, but also the observed sample distribution of $Y$. Hence the PCS p-value is more stable with respect to the observed data and possibly also model mis-specifications. This centered, convoluted null distribution also prevents artificially small p-values that are often obtained in the classical setting (12) with large $n$ and slightly missspecified null distributions.

For the CART based PCS p-value, as in the epiTree pipeline, the test statistic $T(Y)$ as in (7) corresponds to the log-likelihood ratio statistic for the simple null hypothesis $y_i \sim Bernoulli(p_{0,i})$ against the simple alternative $y_i \sim Bernoulli(p_{1,i})$. A classical p-value in this setting corresponds to a Neyman-

Pearson test for these simple null and alternative hypothesis. In particular, for the classical Neyman-Pearson p-value the variance of the test statistic $T(Y)$ under the null distribution $H_0 : Y \sim F_0$ is completely independent of the actual observed responses $Y$. In contrast, the PCS p-value considers the *inflated* null distribution of $T(Y_0) - T(Y)$, with $Y_0 \sim F_0$ and $Y \sim \hat{F}_n$. Hence, the PCS p-value does not just incorporate the distributional variance of $F_0$ but also the observed empirical variance of the response $Y$. We stress that the general concept of the PCS p-value does not depend on the particular choice of statistic $T(Y)$ in (7). As a result, we can calculate PCS p-values not just for the CART based model that we employed for the epiTree pipeline, but also for any other hypothesis testing problem, with an arbitrary test statistic $T(Y)$ and null distribution $F_0$.

In the supplemental Section S1.6, we provide a detailed toy example of a simple linear regression model without intercept, where analytic forms of a classical p-value, as in (12), and PCS p-value, as in (14), can easily be derived analytically. Although this setup is overly simplistic, it provides concrete analytical insights into situations when a PCS p-value is beneficial compared with the a classical p-value. As shown in Section S1.6, the classical p-value has higher power when the data are exactly generated from the hypothetical alternative distribution $H_1$. In this case, the PCS p-value's lower detection power originates from the additional uncertainty quantification in the bootstrap sub-sampling — PCS p-value explicitly takes the empirical variation of the observed data sample into account. However, in arguably most real data applications (and certainly for epistasis testing, as stressed throughout this paper), the data generating process cannot be specified exactly for either the null or the alternative model. In such settings, the PCS p-value can be more robust towards such misspecification. Specifically, we show (both, analytically and in simulations) that classical p-values can result in severe false positive rates when the data are generated from a slight variation of the hypothesised null distribution. On the other hand, the PCS p-value is generally more robust and, in contrast to the classical p-value, does not result in an increased type 1 error. Moreover, we also provide an explicit example in Section S1.6 where the data are generated from a slight variation of the hypothesised alternative distribution. In this case, we observe that the PCS p-value can have a higher detection power compared to the classical p-value. This originates form the fact that the PCS p-value explores the full empirical distribution of the observed responses, and is therefore able to detect deviations from the hypothesized null distribution of the statistic $T(Y)$, which typically cannot be detected from the a single observed $t$, as in the classical p-value. In summary, while the standard p-values work well in settings where everything is specified correctly, when models are misspecified PCS p-values can be favorable – they provide more stable type 1 error control and can have a higher detection power in the presence of outliers.

##### S1.1.8 Analytic approximation of PCS p-value in the epiTree test

In order to obtain an analytic approximation of (11) note that

$$\begin{aligned} T(Y) - T(Y_0) =& \log\left(\frac{P(Y|p_0)}{P(Y|p_1)}\right) - \log\left(\frac{P(Y_0|p_0)}{P(Y_0|p_1)}\right) \\ =& \sum_{i=1}^{n}(y_i - y_{0,i})(\log(p_{1,i}) - \log(1-p_{1,i}) - \log(p_{0,i}) + \log(1-p_{0,i})) \\ =& \sum_{i=1}^{n}\delta_i, \end{aligned}$$

where $p_{1,i}$ and $p_{0,i}$ denote the $i$th component of the vectors $p_1$ and $p_0$, respectively, and $y_{0,i}$ denotes the $i$th component of the $n$-vector $Y_0$, and

$$\delta_i := (y_i - y_{0,i})(\log(p_{1,i}) - \log(1-p_{1,i}) - \log(p_{0,i}) + \log(1-p_{0,i})). \tag{16}$$

For each bootstrap sample $m$, we draw $n$ i.i.d. indexes $I(m)_1, \ldots, I(m)_n$ uniformly from the set $\{1, \ldots, n\}$. Thus we can write

$$\mathbb{1}\{T(Y|m) > T(Y_0|m)\} = \mathbb{1}\left\{\sum_{i=1}^{n}\delta_{I(m)_i} > 0\right\}. \tag{17}$$

Conditioned on the data (with the only randomness coming from the bootstrap sampling via the random indexes $I(m)_1, \ldots, I(m)_n$), the $M$ different terms in equation (11) are independent and identically distributed Bernoulli random variables. Hence, by the law of large numbers and using equation (17), we get that for infinitely many bootstrap samples ($M \to \infty$)

$$\text{PCS p-value} = P_{(I_1,\ldots,I_n)}\left(\sum_{i=1}^{n}\delta_{I_i} > 0\right), \tag{18}$$

where randomness is over the $n$ i.i.d. random indexes $I_1, \ldots, I_n$ drawn uniformly at random from the set $\{1, \ldots, n\}$. Again, conditioned on the data, we have that the $n$ random variables $\delta_{I_1}, \ldots, \delta_{I_n}$ are independent and identically distributed, each following a uniform distribution on the set $\{\delta_1, \ldots, \delta_n\}$. Thus, they have mean

$$E_{I_1}(\delta_{I_1}) = \frac{1}{n}\sum_{i=1}^{n}\delta_i =: \mu \tag{19}$$

and variance

$$Var(\delta_{I_1}) = E_{I_1}((\delta_{I_1} - \mu)^2) = \frac{1}{n}\sum_{i=1}^{n}(\delta_i - \mu)^2 =: \sigma^2, \tag{20}$$

where $E_{I_1}(\cdot)$ mean that we take the expectation w.r.t. the random bootstrap sample. Hence, it follows from the central limit theorem that $X := \sqrt{n}(\frac{1}{n}\sum_{i=1}^{n}\delta_{I_i} - \mu)/\sigma$ converges in distribution to a standard Gaussian as $n \to \infty$ (in our case $n = 4$K). Thus,

$$\text{PCS p-value} = P(X > -\sqrt{n}\mu/\sigma), \tag{21}$$

$$\text{with } |P(X > -\sqrt{n}\mu/\sigma) - \Phi(\sqrt{n}\mu/\sigma)| \to 0, \text{ as } n \to \infty, \tag{22}$$

where $\Phi$ denotes the cumulative distribution function of the standard normal distribution. Note that it follows from (21) that PCS p-value $< 0.5$ if any only if $\mu < 0$, which is equivalent to $T(Y) < T(Y_0)$. Thus the PCS p-value can only be significant when the improvement in prediction (of the epistasis model over the non-epistasis model) for the observed data are greater than for the null perturbation. Further, note one can upper bound the rate of convergence in (21) using the nonuniform Berry–Esseen theorem:

**Theorem 1.** *[23, Theorem 3] Let $Z_1, \ldots, Z_n$ be i.i.d. random variables with $E(Z_1) = 0$, $Var(Z_1) = 1$, and $c_3 := E(|Z_1|^3) < \infty$. Then there exists some constant $L$ such that for all $z \geq 0$*

$$\left| P\left(\sum_{i=1}^{n} Z_i > z\sqrt{n}\right) - (1 - \Phi(z)) \right| \leq \frac{Lc_3}{\sqrt{n}(1 + |z|^3)}.$$

Therefore, from (18) and Theorem 1 we get that when $\mu < 0$

$$|\text{PCS p-value} - \Phi(\sqrt{n}\mu/\sigma)| = \left| P\left(\sum_{i=1}^{n}(\delta_{I_i} - \mu)/\sigma > n(-\mu/\sigma)\right) - (1 - \Phi(\sqrt{n}(-\mu)/\sigma)) \right| \leq \frac{C}{n^2}, \tag{23}$$

with $C = \frac{L \cdot c_3}{|\mu/\sigma|^3} < \infty$ and $c_3 = \frac{1}{n} \cdot \sum_{i=1}^{n} |\delta_i - \mu|^3$. Thus, the approximation error from the CLT for the PCS p-value decreases of order $n^{-2}$. In our case $n = 4,000$, i.e., $n^{-2}$ is of order $10^{-7}$. We note, however, that one also has to take the exact value of $C$ into account (which depends on the data).

**Averaging over null perturbation** For the PCS p-value approximation above, we conditioned on both the observed responses $Y$ as well as the (simulated) null perturbation $Y_0$ (see definition of $\delta_i$ in (16)). In practice, the reported PCS p-value should be independent of any randomness introduced by $Y_0$. Below, we detail two different ways to achieve this:

1. Average the PCS p-value over infinitely many random realizations of $Y_0$. In other words, when computing the expectation and standard deviation of $\delta_{I_i}$ in equation (19) and (20), we average over the randomness of $Y_0$, which is equivalent to replacing the $y_{0,i}$ term with its expectation $p_{0,i}$. This gives

$$E_{I_1,Y_0}(\delta_{I_1}) = \frac{1}{n}\sum_{i=1}^{n} E_{y_{i,0}}(\delta_i) = \frac{1}{n}\sum_{i=1}^{n}\overline{\delta}_i =: \overline{\mu},$$

with

$$\overline{\delta}_i = (y_i - p_{0,i})(\log(p_{1,i}) - \log(1 - p_{1,i}) - \log(p_{0,i}) + \log(1 - p_{0,i})), \tag{24}$$

and

$$Var(\delta_{I_1}) = E_{I_1,Y_0}((\delta_{I_1} - \overline{\mu})^2) = \frac{1}{n}\sum_{i=1}^{n} E_{y_{i,0}}((\delta_i - \overline{\mu})^2) = \frac{1}{n}\sum_{i=1}^{n}\left((\overline{\delta}_i - \overline{\mu})^2 + \omega_i^2 p_{i,0}(1 - p_{i,0})\right) = \overline{\sigma}^2,$$

with

$$\omega_i := \log(p_{1,i}) - \log(1 - p_{1,i}) - \log(p_{0,i}) + \log(1 - p_{0,i}),$$

where $E_{I_1,Y_0}(\cdot)$ means that we take the expectation w.r.t. the random bootstrap index $I_1$ and the random null perturbation $Y_0$. The PCS p-value approximation is then given by $\Phi(\sqrt{n}\overline{\mu}/\overline{\sigma})$.

2. Alternatively, for a given bootstrap sample $m$ in (17), instead of comparing $T(Y|m)$ to $T(Y_0|m)$ for some particular random realization of $Y_0|m$, one can compare $T(Y|m)$ to the average realization of $T(Y_0|m)$, that is, $E_{Y_0}(T(Y_0|m))$. This is equivalent to replacing $\delta_i$ by $\overline{\delta}_i$ from (24) when computing the mean $\mu$ and standard deviation $\sigma$ in (19) and (20). Note that the only different between $\delta_i$ and $\overline{\delta}_i$ is that the random quantity $y_{0,i}$ gets replaced by its expectation $p_{0,i}$. In particular, conditioned on the observed data $(Y, A, B)$, the quantity $\delta_i$ is random, as it depends on the random null perturbation $y_{0,i}$, and $\overline{\delta}_i$ is not random.

Note that the only difference between the two different approaches is in the variance term $\sigma^2$, where the former approach results in a slightly larger variance via the additional $\frac{1}{n}\sum_{i=1}^{n}(\omega_i^2 p_{i,0}(1 - p_{i,0}))$ term. For the red hair analysis, we found that both approaches give the same magnitude for the p-values. The p-values presented in the results section for the red hair analysis correspond to the latter approximation.

###### S1.1.9 Higher order interactions

In the following, we discuss how to extend inference from pairwise to higher-order interactions. Inference for an interaction in the logistic regression model is based on the respective coefficients in a polynomial interaction model. Thus, for higher-order interactions one typically adds higher-order polynomial terms to the model. For example, to test an order-three interaction among genes $A, B, C$, the logistic regression hypothesis problem in equation (3) translates to

$$H_0 : \text{logit}(P(y = 1 \mid a, b, c)) = \beta_A a + \beta_B b + \beta_C c + \beta_{BC} bc + \beta_{AC} ac + \beta_{AB} ab + \beta_0, \tag{25}$$

$$H_1 : \text{logit}(P(y = 1 \mid a, b, c)) = \beta_A a + \beta_B b + \beta_C c + \beta_{BC} bc + \beta_{AC} ac + \beta_{AB} ab + \beta_{ABC} abc + \beta_0. \tag{26}$$

If we replace the polynomial terms in (25) with CART interaction terms and replace the logistic scale by the penetrance scale, we obtain

$$\begin{aligned} H_0 : P(y = 1 \mid a, b, c) &= CART_A(a) + CART_B(b) + CART_C(c) \\ &\quad + CART_{BC}(b, c) + CART_{AC}(a, c) + CART_{AB}(a, b) \\ H_1 : P(y = 1 \mid a, b, c) &= CART_A(a) + CART_B(b) + CART_C(c) \\ &\quad + CART_{BC}(b, c) + CART_{AC}(a, c) + CART_{AB}(a, b) + CART_{ABC}(a, b, c) \end{aligned} \tag{27}$$

Note, however, that in (27), without loss of generality, the lower order CART terms can be incorporated into the respective higher-order CART terms[1]. Hence, for the CART model the direct comparison to (25) is to rewrite equation (5) as

$$\begin{aligned} H_0 &: P(y = 1 \mid a, b, c) = CART_{AB}(a, b) + CART_{BC}(b, c) + CART_{AC}(a, c), \\ H_1 &: P(y = 1 \mid a, b, c) = CART_{ABC}(a, b, c). \end{aligned} \tag{28}$$

More generally, the null model $H_0$ to test for an order-$d$ interaction can be written as an additive model of $d$ CARTs each taking a subset of size $d - 1$ and the alternative epistasis model can be written as a single CART taking all the $d$ features as input.

From a biological perspective, it is not clear whether $H_0$ in (28) — where each of the genes interact with every other gene — should be interpreted as no $d^{th}$ order epistasis. Across the subcellular, cellular, tissue and organismal levels, biological processes are often found to be subdivided into multiple component proteins, such as an enzyme complex or signaling cascade [25]. Considering the genetic basis of human phenotypes, the linkage of multiple genetic variants spanning protein subunits but influencing a specific pathway is an increasingly common finding [8, 28, 14]. Modeling the relationship between all $d$ components in $H_0$ of (28) yields the opportunity to more directly capture the redundancy and non-linear behavior of biological pathways and protein signaling which characterizes observations of epistasis in complex organisms [27, 29]. To illustrate this, consider the relationship between the different features as a graph, where the nodes of the graph correspond to the individual features and edges between features correspond to pairwise interactions. For the null model (no $d^{th}$ order epistasis) $H_0$ in (28), each pair of features is connected via an edge in the graph, as illustrated in Figure S2. From the perspective of a biological pathways, S2 implies that all genes belong to a shared pathway, which arguably represents a $d^{th}$ order biological interaction one may be interested in detecting.

By framing inference w.r.t. to prediction, the epiTree p-values have greater flexibility derived from learning null and alternative models on separate training data. In particular, for the epiTree p-value

[1] To see this, note that any additive CART model of the form $CART_A(a) + CART_{AB}(a, b)$ can be rewritten as a single CART term $\tilde{CART}_{AB}(a, b)$, where $\tilde{CART}_{AB}(a, b)$ is the same as $CART_{AB}(a, b)$ but with additional splits at the tip nodes that correspond to the splits of $CART_A(a)$. The same holds true for higher order interaction CART models.

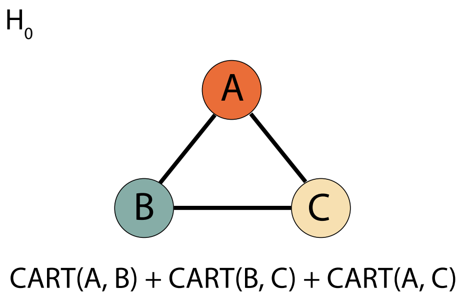

Figure S2: Illustration of the null model $H_0$ in (28).

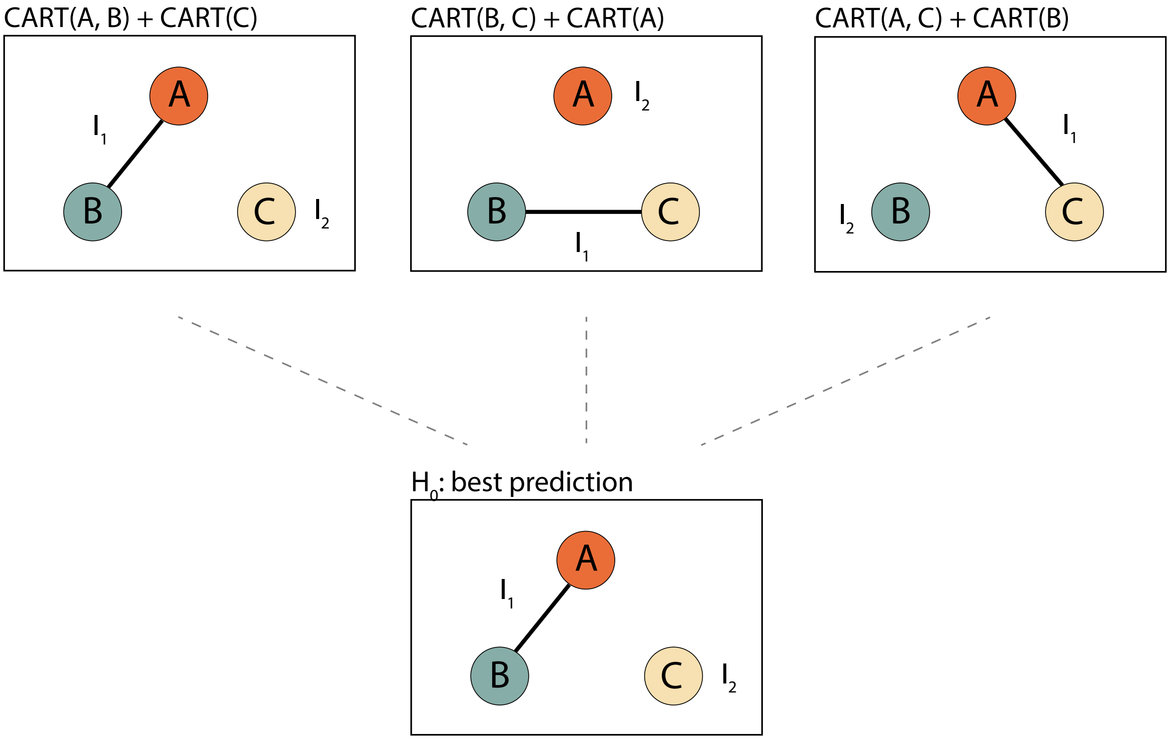

Figure S3: Illustration of the null model $H_0$ in (29), as it is considered for the epiTree p-value.

null model, instead of considering a null model as in (28) and Figure S2, we can learn a partition of the tested interaction features, such that features between different groups do not interact. More precisely, we consider all partitions of the features into two disjoint sets (corresponding to two unconnected graphs), such that features within each set can interact, but features between the two sets can not. The final null model $H_0$ is then selected via a prediction screening, i.e., the partition with the highest prediction accuracy on the test data is selected as the null model $H_0$ which is tested against the interaction model $H_1$ of d-th order epistasis. More precisely, we consider the CART null and alternative model to test a 3rd-order interaction as follows

$$\begin{aligned} &H_0 : P(y = 1 \mid a, b, c) = CART_1(I_1) + CART_2(I_2), \text{ with } I_1 \cup I_2 = \{a, b, c\}, I_1, I_2 \neq \emptyset, \text{ and } I_1 \cap I_2 = \emptyset, \\ &H_1 : P(y = 1 \mid a, b, c) = CART_{ABC}(a, b, c). \end{aligned} \tag{29}$$

We illustrate this case of 3rd-order epistasis null model $H_0$ in Figure S3 and 4th-order epistasis null model $H_0$ in Table S2. Similarly, for a dth-order interaction with features $A_1, \ldots, A_d$, we consider

$$\begin{aligned} H_0 &: P(y = 1 \mid a_1, \ldots, a_d) = CART_1(I_1) + CART_2(I_2), \\ &\quad \text{with } I_1 \cup I_2 = \{a_1, \ldots, a_d\}, I_1, I_2 \neq \emptyset, \text{ and } I_1 \cap I_2 = \emptyset, \\ H_1 &: P(y = 1 \mid a_1, \ldots, a_d) = CART_{A_1 \ldots A_d}(a_1, \ldots, a_d), \end{aligned}$$

where $I_1$ and $I_2$ correspond to the partition which yield the highest prediction accuracy on the test data.[2] Finally, note that in the case of pairwise interactions (i.e. $S = \{A, B\}$) the definitions (28) and (29) are the same.

[2]We evaluate prediction accurarcy analog as in the prediction screening step (8) via the likelihood.

| Candidates for the 4th order no-epistasis $H_0$ model | Final 4th order no-epistasis null model $H_0$ | Alternative 4th order epistasis model $H_1$ |
|---|---|---|
| $M_1 : CART_A(a) + CART_{BCD}(b, c, d)$ | | |
| $M_2 : CART_{A,B}(a, b) + CART_{CD}(c, d)$ | | |
| $M_3 : CART_{A,B,C}(a, b, c) + CART_D(d)$ | $H_0 : M_i$ with | $H_1 : CART_{A,B,C,D}(a, b, c, d)$ |
| $M_4 : CART_B(b) + CART_{ACD}(a, c, d)$ | $P(\text{test data}\|M_i) > P(\text{test data}\|M_j)$ | |
| $M_5 : CART_{BC}(b, c)) + CART_{AD}(a, d)$ | for all $j \neq i$ | |
| $M_6 : CART_{BD}(b, d) + CART_{AC}(a, c)$ | | |
| $M_7 : CART_C(c) + CART_{ABD}(a, b, d)$ | | |
| $M_8 : CART_D(d) + CART_{ABC}(a, b, c)$ | | |

Table S2: Illustration of the null model $H_0$ in (29), but for a 4th order interaction.

##### S1.1.10 SNP features

Due to the discreteness of SNP features, there is no need to consider different scaling functions $f_A$, $f_B$, $f_{AB}$ as in equation (4). This is because any (bijective) scaling will simply map the three different possible SNP values $\{0, 1, 2\}$ to some other distinct values. Thus, equation (3) becomes

$$\begin{aligned} H_0 : \text{logit}(P(y = 1 \mid a, b)) =& \beta_0 + \sum_{i=1}^{2} \beta_{Ai} \mathbb{1}_{a=i} + \sum_{j=1}^{2} \beta_{Bj} \mathbb{1}_{b=j} \\ H_1 : \text{logit}(P(y = 1 \mid a, b)) =& \beta_0 + \sum_{i=1}^{2} \beta_{Ai} \mathbb{1}_{a=i} + \sum_{j=1}^{2} \beta_{Bj} \mathbb{1}_{b=j} + \sum_{i=1}^{2} \sum_{j=1}^{2} \beta_{ABij} \mathbb{1}_{a=i,b=j} \end{aligned}$$

and equation (5) becomes

$$\begin{aligned} H_0 : \text{logit}(P(y = 1 \mid a, b)) =& \beta_0 + \sum_{i=1}^{2} \beta_{Ai} \mathbb{1}_{a=i} + \sum_{j=1}^{2} \beta_{Bj} \mathbb{1}_{b=j} \\ H_1 : \text{logit}(P(y = 1 \mid a, b)) =& \beta_0 + \sum_{i=1}^{2} \beta_{Ai} \mathbb{1}_{a=i} + \sum_{j=1}^{2} \beta_{Bj} \mathbb{1}_{b=j} + \sum_{i=1}^{2} \sum_{j=1}^{2} \beta_{ABij} \mathbb{1}_{a=i,b=j} \end{aligned}$$

For more than two genes the situation is analog as in Section S1.1.9, where the null model allows for interactions up to order $K-1$, when an interaction of order $K$ gets tested. That means, for an interaction of order $K$ the alternative model (epistasis) has $3^K$ parameters and the null model (no-epistasis) has $3^K - 2^K$ parameters.

#### S1.2 Further discussion on comparison between CART models and logistic regression for specific red hair interactions

In the results section of the main text we investigated the difference between CART based models and logistic regression on the response surfaces of the null (no-epistasis) and alternative (epistasis) models for two exemplary cases: *ASIP* - *TUBB3* and *ASIP-DEF8*. Here we provide a similar discussion for the other three pairwise interactions with significant p-values from either PCS or logistic regression.

The response surface for a putative interaction among *ASIP* - *DBNDD1* is shown in Figure **??**. Here, the PCS p-value ($= 10^{-11}$) is much smaller than the logistic regression p-value ($= 10^{-1}$). However, the prediction error for the CART alternative model (epistasis) (cross entropy of 0.528) is slightly worse than the prediction error of the logistic regression null model (no-epistasis) (cross entropy of 0.516). Looking at the response surface, we find that the thresholding behavior observed for many of the other interactions (recall the results section in the main text) is less present for these features. As a consequence the smooth response surface of the additive logistic regression model provides a similar data fit as the non-additive CART interaction model, such that overall the interaction behavior is less clear in this case and would require further investigation on the SNP level. Figure S10 shows the response surface for *ASIP* - *VS9D1*. Here, the data show a clear nonlinear thresholding behavior, which is reflected in the smaller prediction error of the CART models (cross entropy at around 0.5) compared to the logistic regression model (cross entropy at around 0.7). The final response surface shown in Figure S11 considers an interaction among *ASIP* - *GAS8*. Here, the logistic regression p-value ($= 10^{-7}$) is much smaller than the PCS p-value. However, all four models (CART null and alternative, logistic regression null and alternative) have a similar prediction error (cross entropy at 0.589, 0.596, 0.589, 0.585, respectively). From this, we conclude that there is less evidence for epistasis in this case, as the additive CART model can explain the data just as well as a multiplicative interaction term.

#### S1.3 An alternative approach to imputed gene expression dimension reduction via sub-batching

The extremely high dimensionality of the SNP features prohibited us from running iRF on the SNP data directly. Therefore, in the epiTree pipeline we followed a biologically inspired dimension reduction step via imputed gene expression and then first searched for interactions on the gene level, before going back to the SNP level data. In the following, we describe an alternative approach which we also explored.

We performed an initial screen for important variants across sub-batches of SNPs. First, we partitioned the full dataset into batches of 10,000 SNPs and fit a RF to predict the red hair phenotype from each batch. We used the RF feature importance – mean decrease in Gini impurity (MDI) – to rank SNPs and fitted separate iRFs using the top: 100, 500, 1000, and 2500 SNPs. This strategy parallels marginal screening approaches, which filter a large set of variants based on associations between a single gene and the target response. However, by using RF feature importance to screen SNPs, we maintain the possibility of including interacting variants with weak main effects since decision paths reflect the importance of a feature conditioned on previous splits in the path.

In principle, one advantage of the sub-batch over gene expression screening by iRF is that important SNPs that do not influence gene expression are more likely to pass the initial screening procedure. That is, SNPs that are not important in the PrediXcan model have a better chance of being picked up by iRF. However, we found that regions with particularly strong effects (like MC1R for red hair) tend to mask SNPs from regions with weaker effects. Aggregating SNPs at the gene level gives weaker SNPs a higher chance to pass the initial screen. Interpreting interactions directly on the gene level also has practical advantages. Follow up experiments are much easier to be performed on the gene level then on the SNP level. Therefore, we focused on the gene-expression approach in our analysis.

#### S1.4 Choice of tuning parameters to fit the CART interactions

The CART algorithm, which is applied to fit the tree components of $H_0$ and $H_1$ in (5) has several tuning parameters, such as the complexity parameter and the minimal number of observations required at each leave node, which control how deep the tree is grown. In general, the PCS p-value depends on the particular trees which are tested in (5) and thus, depends on the CART tuning parameters. Our aim was to select the trees in (5) as simple as possible, but deep enough such that they capture the potential interaction behaviour of the considered genes. To this end, we controlled the depth of the tree using the complexity parameter (`cp` in `rpart`), which controls the minimum improvement in the model needed at each node. The larger `cp` is, the shallower is the fitted tree. If `cp` is very large, the resulting interaction tree $CART_{AB}$ in (5) might split only on one of the two genes $A$ or $B$. In that case, it is clearly not possible for $CART_{AB}$ to capture interaction behaviour of $A$ and $B$. Analog holds for higher-order interactions (recall Section S1.1.9), if the resulting tree does not contain all genes of that interaction. Therefore, we selected the complexity parameter (`cp` in `rpart`) for the interaction model $CART_{AB}$ adaptively, namely, just small enough such that the interaction model included all genes (but not smaller than 0.01, which is the default parameter in the `rpart` implementation). More precisely, we start with `cp` = 0.01 and then, as long as not all features get split on in the tree, we replace the current `cp` value by `cp /1.1`. For the additive components, $CART_A$ and $CART_B$, the complexity parameter is chosen to be the same as for the interaction component. In this way it is guaranteed that the tree for the interaction model, $CART_{AB}$, is sufficiently deep to capture potential interaction behavior. Moreover, we prevent overfitting by not making `cp` smaller than necessary for all genes to appear in the tree and generally not smaller than 0.01. At the same time, this guarantees that the complexity of all trees, $CART_{AB}, CART_A, CART_B$, is comparable.

We analyzed the stability of the PCS p-value with respect to this choice of tuning parameter in the CART algorithm. This is illustrated for the candidate interaction DBNDD1 and ASIP in Figure S4. The top plot shows on the x-axis the complexity parameter of the CART algorithm which was used to fit the models in (5) and on the y-axis it shows the respective p-value. When the complexity parameter is too large (left side of the plot), the resulting interaction tree does not contain any splits on ASIP (as illustrated in Figure S4) and thus, cannot capture interaction behaviour between ASIP and DBNDD1. The dashed line corresponds to the largest complexity parameter, such that the tree splits on both DBNDD1 and ASIP. This is the tree which we considered as interaction model for the PCS p-value (see the dashed line in Figure S4 and the tree marked with a red arrow), leading to a highly significant PCS p-value in this case. As the complexity parameter decreases, the tree is grown deeper (as illustrated in Figure S4) and the p-values changes, as can be seen in the jumps in Figure S4). However, we note that although the trees change, the p-values remains significant for a wide range of smaller complexity parameters. Figure S4 shows the same plots for all other candidate interactions which were detected by the iRF algorithm (recall Figure ??). In general, we observe that when the PCS p-value is significant, then it also remains significant over a wide range of complexity parameters. Figure S6 summarizes this observations, where the x-axis shows the PCS p-value of the different candidate interactions and the y-axis shows the fraction of complexity parameters[3] where the resulting p-value is smaller than 0.01. This implies that significant PCS p-values appear to be robust to reasonable changes of tuning parameters of the CART algorithm. On the other hand, we note that for those interactions where the PCS p-value was not significant, there are still some interactions where smaller complexity parameters consistently result in significant p-values. In practice, one might consider these candidate interactions for further investigation, but we stress that such a screening through complexity parameters does not account for multiple testing.

#### S1.5 Details on parameter choices for iRF, ranger, and penalized logistic regression

In the following, we provide further specific details on the choice of implementation and software parameters made in our analysis of the red hair phenotype for UK Biobank data with the epiTree pipeline.

**Penalized logistic regression:** The penalized logistic regression model was fit with the R package `glmnet`, using default parameters for classification. The tuning parameter `lambda` was selected via cross validation using the `cv.glmnet` function of the `glmnet` R package.

**Random forest:** The RF model was implemented in the R package `ranger` with default parameters for classification. We note that our iRF model was also fit using `ranger`. As a result, differences between iRF and RF can be directly attributed to iterative feature re-weighting, which is a form of "soft" regularization (see more details below).

**iRF:** For the iRF model, for $k = 10, 50, 100, 500, 1000, 2500$, we ran the first iteration with default parameters for classification. We then applied hard thresholding on the top $k$ features, with $k$ selected by minimizing out-of-bag error, followed by three iterations of iterative re-weighting (i.e. soft-thresholding). For the gene level analysis $k = 50$, while the subsequent SNP level analysis used $k = 1,000$. After the final iteration, we then searched for interactions using RIT, where we grew many shallow trees, in order to obtain a large set of candidate interactions ($ntree = 5,000, depth = 3, child = 5$). To evaluate stability of candidate interactions, we performed $n.bootstrap = 50$ bootstrap replicates and only included candidate interactions that were consistently filtered in at least 50% of the replicates.

[3]Considering `cp` values on a log-scale up to one unit smaller than considered in the PCS p-value, that is, the range of `cp` values shown right of the dashed line in Figure S5.

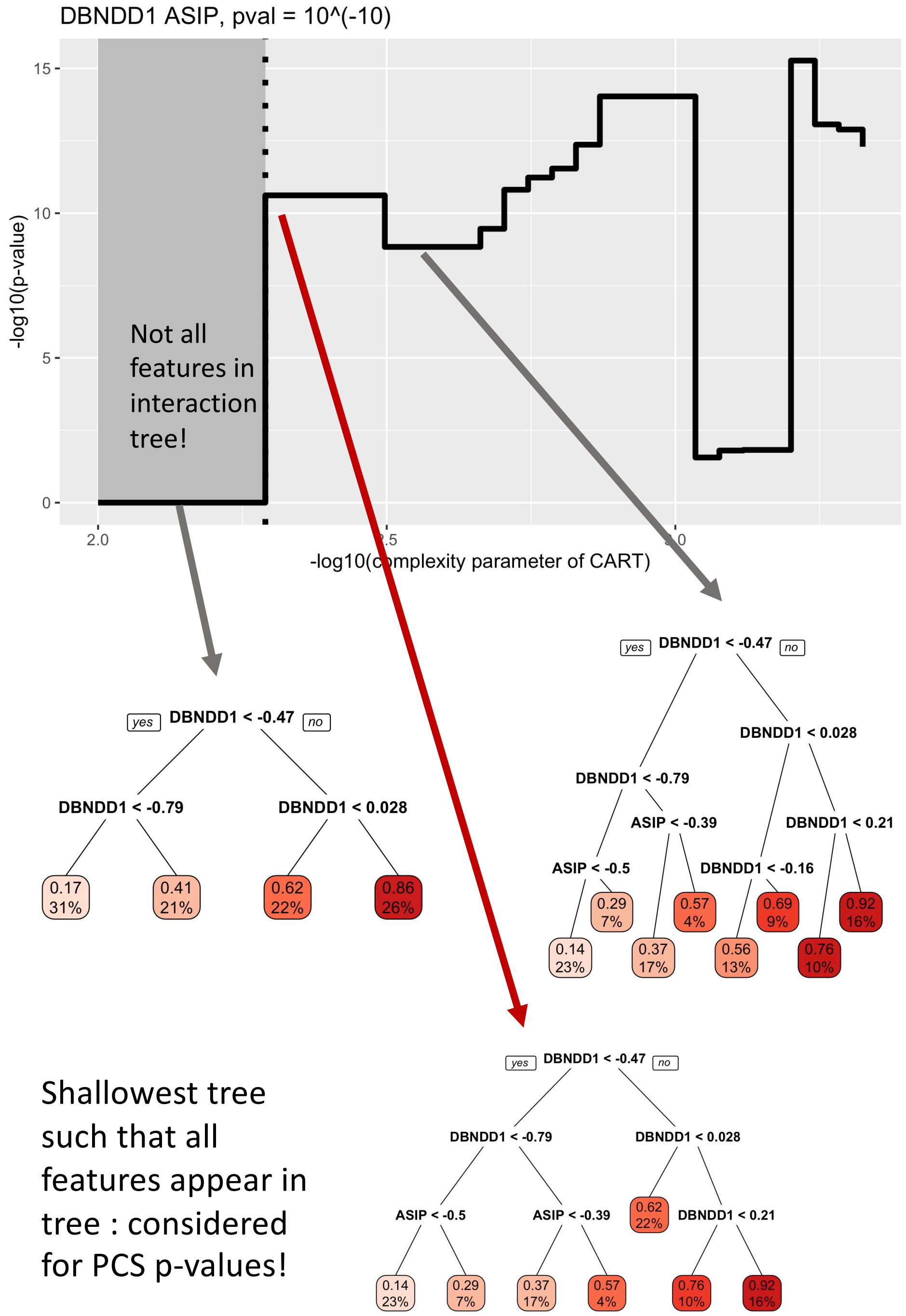

Figure S4: Obtained p-value (y-axis) for different choices of the complexity parameter `cp` (x-axis) in the CART implementation of `rpart` for the candidate interaction between ASIP and DBNDD1. The dashed line corresponds to the `cp` value which was considered for the PCS p-value and the gray highlighted area corresponds to those `cp` values where the resulting CART interaction model of ASIP and DBNDD1 does only split on DBNDD1 but not on ASIP. Three different trees, which correspond to the interaction model of some `cp` values are shown. Left: a tree for a large `cp` value, which only splits on DBNDD1 but not on ASIP and thus cannot capture interaction behaviour; middle: the shallowest tree such that both, DBNDD1 and ASIP, appear in the tree, this tree is considered for the PCS p-value; right: a deeper tree which corresponds to a smaller `cp` value.

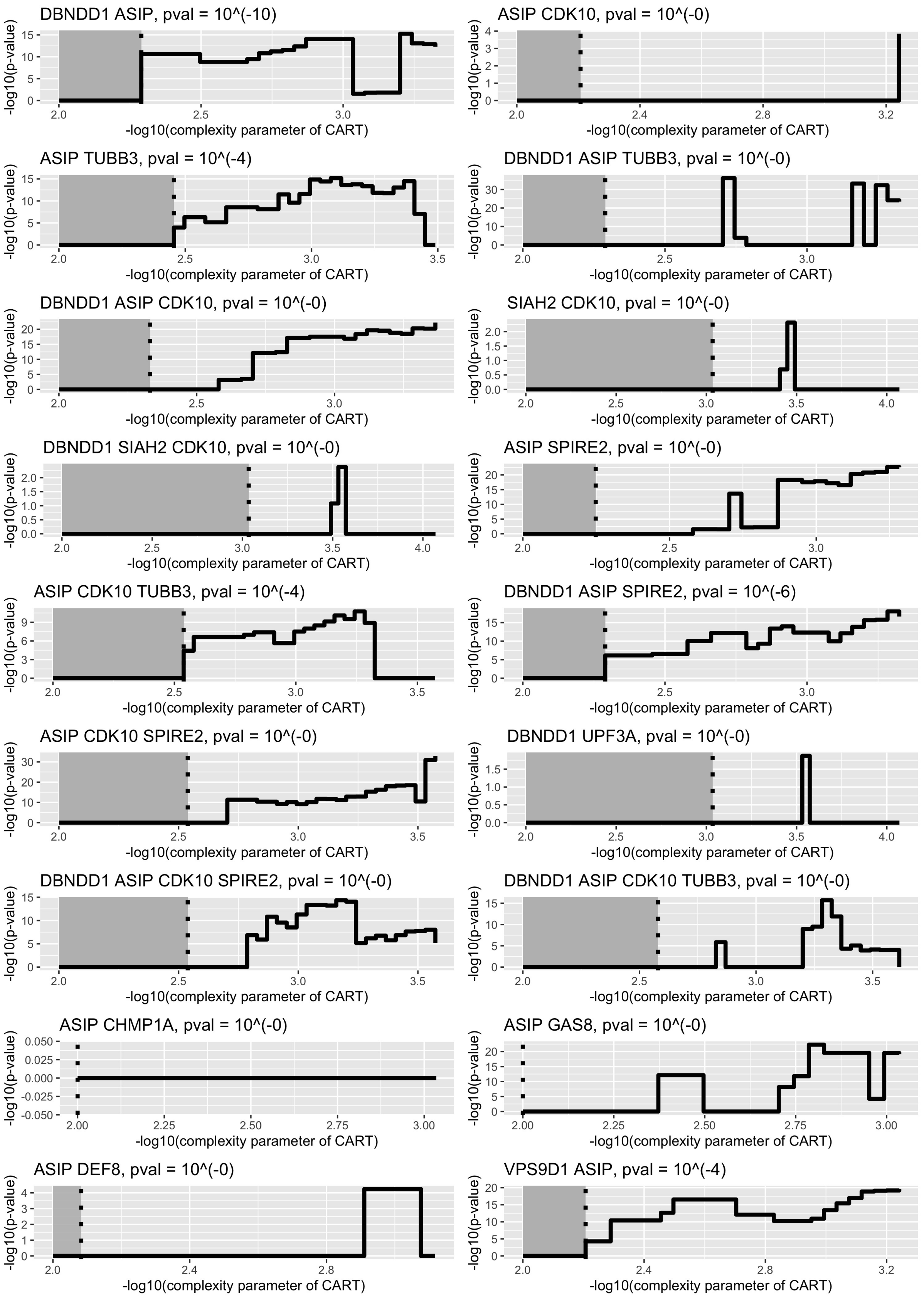

Figure S5: Obtained p-value (y-axis) for different choices of the complexity parameter `cp` (x-axis) in the CART implementation of `rpart`, analog as in Figure S4.

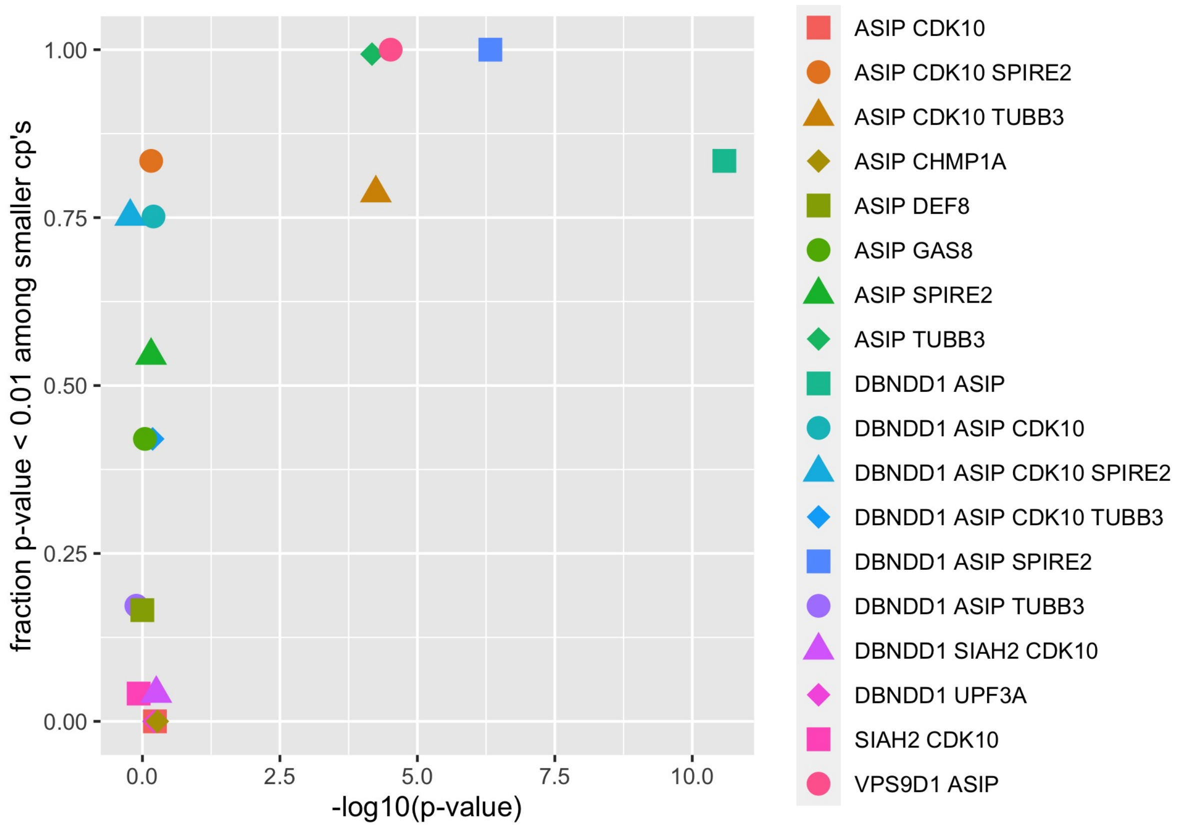

Figure S6: The x-axis shows the PCS p-value of the candidate interactions which were detected by the iRF screening procedure, as in Figure **??**. The y-axis shows the fraction of complexity parameters where the resulting p-value is smaller than 0.01. For this fraction we considered all complexity parameters `cp` on a log-scale up to one unit smaller than considered in the PCS p-value, that is, the range of `cp` values shown right of the dashed lines in Figure S5.

#### S1.6 Illustrative example of PCS p-value for simple linear regression with one predictor

Here we compare classical and PCS p-values using a toy example: a linear model with a single feature or predictor variable, given unit variance, and no intercept. In this simple setting, one can easily derive closed form solutions for the classical p-value and the PCS p-value. We find that in situations where the data is generated exactly from the hypothesized alternative model, the PCS p-value has reasonable detection power, but the classical p-value is superior with fewer false negatives than the PCS p-value. However, when models are misspecified, the PCS p-value is more robust, leading to fewer false positives under a misspecified null model and fewer false negatives under a misspecified alternative model. In most practical situations the hypothesised models are misspecified to some extent, especially in the big data era. This suggests that the PCS p-value is favorable for most real data applications.

We consider a test data set $\{(y_1, x_1), \ldots, (y_n, x_n)\}$ with response values $y_i \in \mathbb{R}$ and a single, fixed covariate $x_i \in \mathbb{R}$, along with a separate training data set $\{(\tilde{y}_1, \tilde{x}_1), \ldots, (\tilde{y}_n, \tilde{x}_n)\}$, where we have assumed for simplicity that the training and test data are of the same size. Further, to ease notation, in the following we assume that $\sum_{i=1}^n x_i^2 = \sum_{i=1}^n \tilde{x}_i^2 = 1$. Our goal is to obtain and compare classical and PCS p-values for the hypothesis testing problem

$$H_0 : \; y_i \overset{i.i.d.}{\sim} \mathcal{N}(0,1) \quad \text{vs.} \quad H_1 : \; y_i \overset{ind.}{\sim} \mathcal{N}(c\, x_i, 1) \text{ for some } c > 0. \tag{30}$$

**Test statistic:** We consider the test statistic as in a classical *z-test*, that is

$$T(Y) = \sum_{i=1}^n x_i y_i, \tag{31}$$

with $Y = (y_1, \ldots, y_n)^\top$. The classical p-value from the *z-test* is given by

$$\text{p-value } = \Phi\left(-T(Y)\right) = \Phi\left(-\sum_{i=1}^n x_i y_i\right), \tag{32}$$

where, again, $\Phi$ denotes the cumulative distribution function of the standard normal distribution. In the following examples we compare this classical p-value to the respective PCS p-value. For the PCS p-value we first obtain an estimate for both, the hypothesis model and the alternative model, from the training data. For the alternative $H_1$ we obtain an estimate for the coefficient $c$ from the training data $\{(\tilde{y}_i, \tilde{x}_i), i = 1, \ldots, n\}$ as

$$\hat{c} = \sum_{i=1}^n \tilde{x}_i \tilde{y}_i. \tag{33}$$

For the null hypothesis $H_0$ there are no unknown parameters that need to be estimated.

**PCS Prediction screening:** In the prediction screening step, we use the test data to evaluate the prediction error from the null model and the alternative model, where here we estimate the alternative model using the training data as in (33). For the loss function, we consider the negative log-likelihood, which equals the squared loss in the Gaussian case (30). More precisely, the prediction screening as in (8) yields

$$\text{PCS p-value } = 1 \text{ if } \sum_{i=1}^{n} y_i^2 \leq \sum_{i=1}^{n} (y_i - \hat{c}x_i)^2 \quad \Leftrightarrow \quad \sum_{i=1}^{n} x_i y_i \leq \frac{1}{2}\hat{c}. \tag{34}$$

**PCS Null perturbation:** When the prediction screening does not result in a failure to reject the null hypothesis, we simulate from the null perturbation. In this simple setting, the null hypothesis corresponds to i.i.d. Gaussian white noise. Therefore, we perturb our true responses $y_i$ as follows:

$$Y_0 = (y_{0,1}, \ldots, y_{0,n})^\top \sim \mathcal{N}((0, \ldots, 0)^\top, I_{n\times n}), \tag{35}$$

where $I_{n\times n}$ denotes the $n \times n$ identity matrix.

**PCS p-value:** We obtain a PCS p-value with test statistic $T(Y)$ as in (31) and null perturbation $Y_0$ as in (35) via bootstrap samples as in (11). Note that in this case we can approximate the PCS p-value as in (14) such that

$$\text{PCS p-value } \approx P\left(\sum_{i=1}^{n} x_i \left(y_i - y_{0,i}\right) < 0\right). \tag{36}$$

##### S1.6.1 Simulation studies

In the following simulations, we compare the classical p-value as in (32) and the PCS p-value as in (36) under different probabilistic generating models for the responses $y_i$ (and $\tilde{y}_i$, respectively). We note that the PCS p-value requires a separate training data set, which is not required by the classical p-value. In order to provide a fair comparison in all of our simulations studies , we use all available data as test data for the classical p-value and for the PCS p-value we randomly split all available data into a training and a test data sets of equal size. Below, we summarize our results. More details, as well as further examples, can be found in a supplementary R Markdown file, available at `https://github.com/merlebehr/epiTree/example_pcs_pvalues_linear_regression.R`.

**Classical setting – correctly specified model.** Here we compare PCS and classical p-values under the correctly specified model where data are generated exactly as indicated by (i) the null hypothesis $H_0$ and (ii) the alternative hypothesis $H_1$. In these settings, the classical p-value outperforms the PCS p-value. That is, the classical p-value has higher power under the alternative and maintains the correct

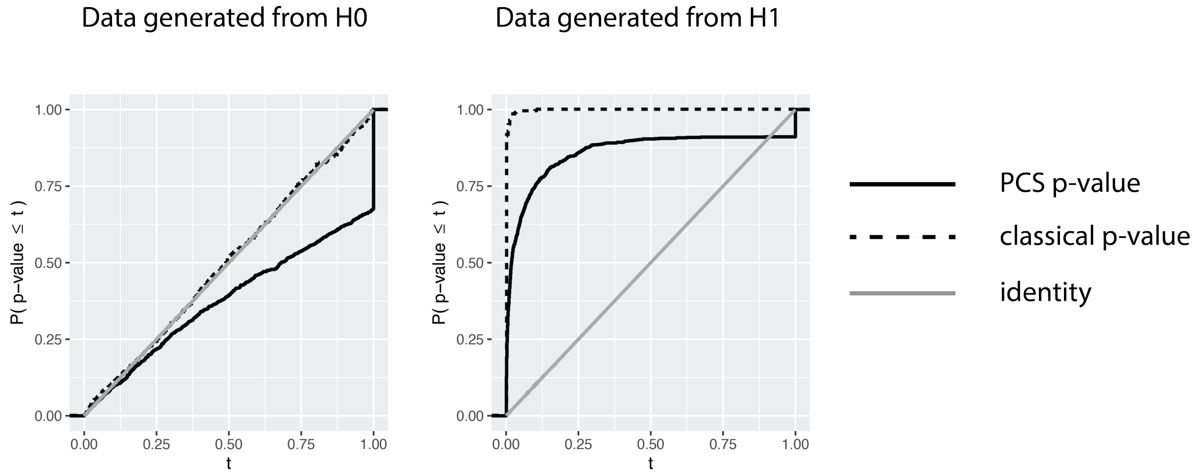

Figure S7: Distribution of classical p-value and PCS p-value for the hypothesis testing problem in (30) with test statistic as in (31) for $n = 1,000$ and a single dependent variables $x_i$ drawn independently from a uniform distribution on $[0, 1]$. The black solid line corresponds to the PCS p-value, the black dotted lines to the classical p-value, and the gray solid line to the identity. Results are obtained from $1,000$ Monte Carlo runs. Left: responses $y_i$ as in $H_0$ in (30); Right: responses $y_i$ as in $H_1$ in (30) with $c = 4$.

control over type I error. Of course, we note that in real data settings the "true" model is rarely known and thus this performance is not expected in practice.

First, we consider the setting where the responses are generated from the $H_0$ distribution as in (30) with $y_i \sim \mathcal{N}(0, 1)$ independent for all $i = 1, \ldots, n$. As seen in the left plot in Figure S7, both, the classical p-value and the PCS p-value follow a sub-uniform distribution, indicated by the fact that their cumulative distribution function (cdf) is smaller or equal to the identity line. The PCS p-value is a bit more conservative than the classical p-value under the exact null distribution, indicated by the fact that the cdf of the PCS p-value is smaller or equal to the cdf of the classical p-value. However, for smaller quantiles (that are of dominant interest in practice) the different between the PCS p-value and the classical p-value distributions is almost negligible under the exact null distribution. For example, in our Monte Carlo simulations we found that the PCS p-value is smaller than 0.05 in exactly 5% of cases, which coincides with the classical p-value.

Second, we consider the situation where the responses follow a model as in $H_1$ in (30) with $y_i \sim \mathcal{N}(c\, x_i, 1)$ independent for all $i = 1, \ldots, n$, for some fixed $c > 0$. The right plot in Figure S7 shows the respective p-value distributions for $c = 4$. In this case, we find that the classical p-value outperforms the PCS p-value, although the PCS p-value has high power. For example, at a nominal level of $\alpha = 0.05$ the classical p-value correctly rejects the null hypothesis in 99% of cases, but the PCS p-value rejects in only 64% of cases. Similar, at nominal level $\alpha = 0.1$ the classical p-value rejects in 99.7% of cases, but the PCS p-value in only 74.9% of cases. Intuitively, the reason for this is that exploring the full distribution of the test statistic $T(Y)$ via the bootstrap sampling results in an additional variance term, which leads to a weaker detection power when the observed responses exactly follow the specified alternative model $H_1$.

**Misspecified model** Here we compare PCS and classical p-values when data are generated under a model that deviates slightly from the null $H_0$ via a misspecified variance. That is, we assume that the variance of observations $Y_i$ in (30) is not 1 but $\sigma^2 > 1$. In this setting, we argue that it is preferable *not* to reject $H_0$. We find that the PCS p-value is robust to this model misspecification and behaves largely the same way as in the correctly specified setting. In contrast, the classical p-value is highly sensitive to this model misspecification, even in the simple linear model considered in our simulations.

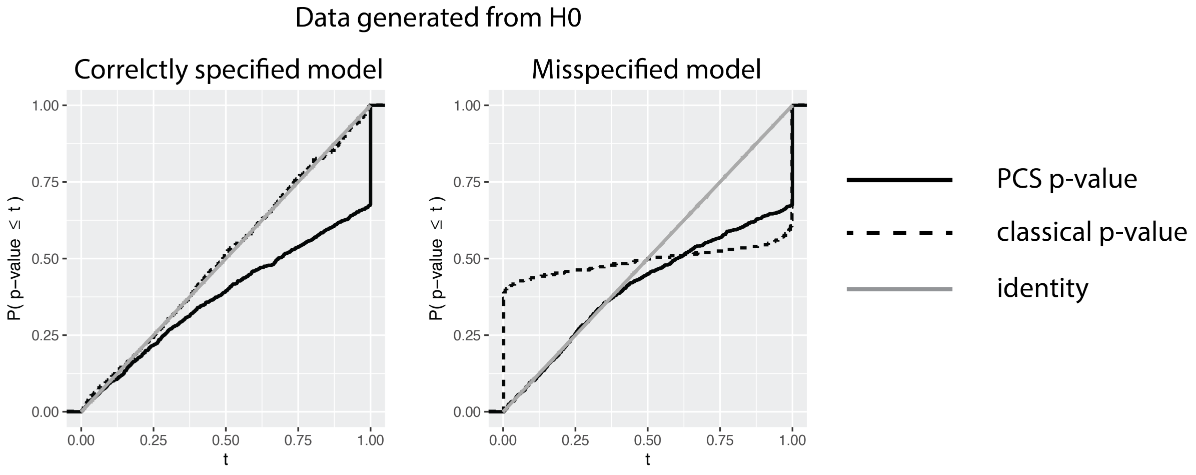

Figure S8: Same as Figure S7, but with the right plot such that responses $y_i \overset{i.i.d.}{\sim} \mathcal{N}(0, \sigma^2)$ with $\sigma = 10$.

We let responses follow the null distribution $H_0$ but with a larger variance $\sigma^2 \gg 1$. That is, $y_i \overset{i.i.d.}{\sim} \mathcal{N}(0, \sigma^2)$. The right plot in Figure S8 shows the respective p-value distributions when $\sigma = 10$ (results for different values of $\sigma$ are shown in the R markdown file at `https://github.com/merlebehr/epiTree/example_pcs_pvalues_linear_regression.R`). In this case, we find that the classical p-value yields severe false positives due to the model misspecification. For example, at a nominal level of $\alpha = 0.05$ the classical p-value falsely rejects the null hypothesis in 42% of cases. In contrast, the PCS p-value, via the prediction screening as well as the bootstrap sampling, is robust to this misspecification. E.g., at nominal level $\alpha = 0.05$ the PCS p-value falsely rejects the null hypothesis in only 5% of cases, as is expected under the null. Similar, at nominal level $\alpha = 0.01$ the PCS p-value rejects in %1 of cases, but the classical p-value still rejects in as much as 40% of cases. We stress that even when the null model is extremely misspecified with a very large $\sigma$ remarkably the PCS p-value is robust to this with no increase in the type 1 error (see the supplementary R markdown file at `https://github.com/merlebehr/epiTree/example_pcs_pvalues_linear_regression.R`).

In summary, in our simple simulation set-up, the PCS p-value is found to be more robust to a misspecified data generating process as it is almost always the case in practice, even though, as expected, the classical p-value is found to work better than the PCS p-value when the data generative models are specified correctly. In particular, the use of PCS p-values results in fewer false positives under the null. Moreover, as detailed in the supplementary R markdown file at `https://github.com/merlebehr/epiTree/example_pcs_pvalues_linear_regression.R`, the PCS p-value can result in fewer false negatives when the alternative is misspecified. Of course, our simulation set-up is very simple. We nevertheless believe that the lessons learned hold in general. It is our current research to provide more extensive evidence to support our belief that in practice the PCS p-value should be preferred over the classical p-value, unless the very precise forms of $H_0$ and $H_1$ are carefully backed up by domain knowledge and quantitative empirical evidence.

#### S1.7 Comparison between PCS and other inference methods

We compared the performance of epiTree (PCS epistasis p-values) against (i) logistic regression-based inference and (ii) permutation random forests (pRF) [19]. Logistic regression has been widely used to estimate epistasis (e.g. [15, 22, 30, 33]), but is formulated to detect multiplicative interactions after controlling for linear main effects. Permutation random forests (pRF) leverage random forests, as epiTree does, to detect interactions of arbitrary form. However, pRF is designed to detect the presence of pairwise, rather than higher-order interactions. We note that an extension of pRF to high-order interactions could be formulated in the context of PCS inference, with permutations serving as the null perturbation, but we do not explore this extension here.

We evaluated the performance of epiTree and pRF under four models with responses generated from the gene expression data used in our red hair case study (subset to the 50 genes with highest MDI importance). For each model, we generated Bernoulli responses from sets of active interactions $S_k$, $k = 1, 2$ (described below), identified candidate interactions using iRF, and computed p-values from each method for all candidate interactions. We note that each method uses a different inference technique for computing p-values (logistic regression: classical, epiTree: PCS, pRF: permutation-based). The probability of success $\pi$ for Bernoulli responses in the four models was given, respectively, by:

$$
\begin{aligned}
\pi^{(AND)} &= \pi_A \left( \prod_{j \in S_1} \mathbb{1}(x_j > t_j) \right) + \pi_I \left( 1 - \prod_{j \in S_1} \mathbb{1}(x_j > t_j) \right), \\
\pi^{(OR)} &= \pi_A \left( \max_{k=1,2} \prod_{j \in S_k} \mathbb{1}(x_j > t_j) \right) + \pi_I \left( 1 - \max_{k=1,2} \prod_{j \in S_k} \mathbb{1}(x_j > t_j) \right), \\
\pi^{(ADD)} &= \pi_A \left( \prod_{j \in S_1} \mathbb{1}(x_j > t_j) + \prod_{j \in S_2} \mathbb{1}(x_j > t_j) \right) \\
&+ \pi_I \cdot \mathbb{1} \left\{ \left( \prod_{j \in S_1} \mathbb{1}(x_j > t_j) + \prod_{j \in S_2} \mathbb{1}(x_j > t_j) \right) = 0 \right\}, \\
\pi^{(MULT)} &= \pi_A \cdot \sigma \left( \prod_{j \in S_k} x_j \right),
\end{aligned}
$$

where $\sigma$ denotes the sigmoid function and $\pi_A$, $\pi_I$ baseline success probabilities for active and inactive samples respectively. We set $\pi_A = 0.75$ and $\pi_I = 0.25 \cdot \{\text{proportion of active samples}\}$ for all models. We selected features in active interactions $S_k$ using an iterative sampling strategy to prevent high levels of correlation between interacting features. Specifically, the first feature was sampled with uniform probability and subsequent features with probability inversely proportional to their maximum absolute correlation over previously selected features. For all models, we set the size of each interaction component to $|S_k| = 3, k = 1, 2$ (i.e. an order-3 interaction).

We note that the different response generating models represent a range of interaction forms. The AND model ($\pi^{(AND)}$) corresponds to a single interaction that is either active or inactive for all observations. The OR model ($\pi^{(OR)}$) corresponds to two distinct interactions that can be viewed as heterogeneous mechanisms governing response behavior. The ADD model ($\pi^{(ADD)}$) corresponds to two interactions terms that do not interact with one another. Finally, the multiplicative model ($\pi^{(MULT)}$) corresponds to a multiplactive interactions.

For each model above, we generated a list of candidate interactions using iRF and computed p-values using logistic regression, pRF, and epiTree. To assess the variability in p-values, we repeated the inference step 25 times on new responses generated from the same set of active interactions. Figure S21 reports the results from each response generating model. We report the distribution of $-\log_{10}(p)$ for the following groups: (i) active interactions — the true response generating interaction (i.e. $S = S_1$ or $S = S_2$) (ii) active interaction subsets — a strict subset of the true response generating interaction (i.e. $S \subset S_k$) (iii) active features — features used in active interactions but not all part of the same interaction term (i.e. $S \subseteq S_1 \cup S_2, S \nsubseteq S_1, S \nsubseteq S_2$) (iv) inactive interaction — includes features that do not appear in any interaction term (i.e. $S \cap (S_1 \cup S_2) \neq \emptyset$). We note that the resolution of p-values for pRF depends on the number of permutations run, which we set to 1000 in our simulations. For consistent comparison across all methods, we threshold reported p-values below at $10^{-4}$. Below we summarize several key findings.

- **AND/OR models:** All methods consistently detect (at significance level 0.05) active interactions. epiTree and logistic regression also detects subsets of active interactions while pRF generally does not. We find that active interactions fall at the top of the p-value ranked list for epiTree and pRF but not for logistic regression. In other words, the interactions deemed most important (with respect to p-value) by epiTree and pRF included only features that were truly interacting.

- **AND model:** pRF consistently detects false positives in the AND model (average p-value of inactive interactions $\sim 0.025$). We find that this is due to the fact that pRF p-values are generally low for interactions that contain both an active interaction and other non-interacting features. In contrast, epiTree and LR rarely detect these false positive interactions. The average p-values for epiTree and LR are consistently $> 0.25$.

- **ADD model:** epiTree is the only method that consistently detects active interactions (and their subsets) in the ADD model. We find that active interactions are at the top of the p-value ranked list for epiTree in the ADD model but not for pRF or logistic regression.

- **MULT model:** Logistic regression is the only method that consistently detects subsets of multiplicative interactions. We note that no method computes p-values for the active interaction since it was not detected by iRF, and thus not contained in the set of candidate interactions. While epiTree does not detect subsets of the active multiplicative interaction at a significance level of 0.05, we find that such subsets still fall at the top of the p-value ranked list for epiTree.

In summary, epiTree performs comparably to or better than competing methods under the AND, OR, and ADD models. Specifically, epiTree consistently recovers true interactions with generally lower false positive rates than pRF or logistic regression. While logistic regression slightly outperforms epiTree under the multiplicative model, all methods perform poorly in this setting.

#### S1.8 Further details on the multiple sclerosis case study

For the multiple sclerosis case study we considered a total of 502,369 individuals from the UK Biobank cohort, aged between 40-69 at recruitment (MS status defined using ICD-10 code G35). We searched for gene-level interactions in a balanced, random sample of cases and controls (2083 MS individuals and 2083 controls). To help assess the generalizability of our results to unobserved data, we performed a random sample split into training and test sets with 3333 training samples and 833 test samples. We modeled MS using ∼62,000,000 common variants (MAF $> 0.01$) imputed to the Haplotype Reference Consortium (HRC) and UK10K reference panels from ∼800,000 directly genotyped variants, which were obtained by UK Biobank using one of two similar arrays [5]. Details regarding the ascertainment and quality control of these genotypes have been previously described [5]. We estimated gene expression levels in brain cortex tissue from the individual SNP data using CTIMP [1].

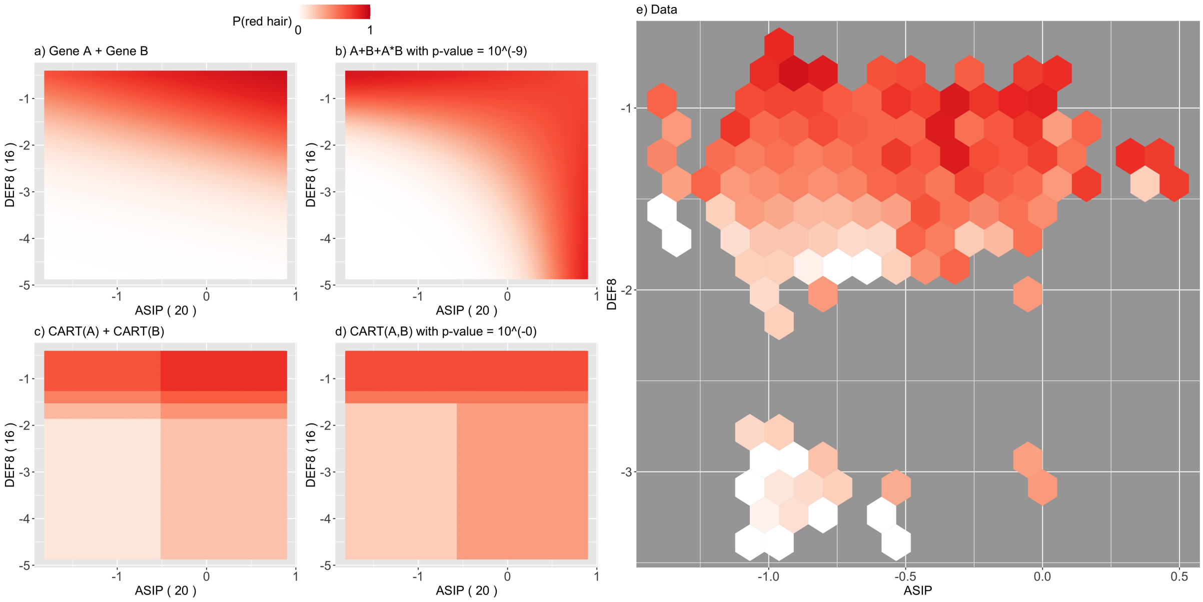

Figure S9: Response surface for *ASIP* - *DEF8*, otherwise as Figure ??

### S2 Supplemental Figures

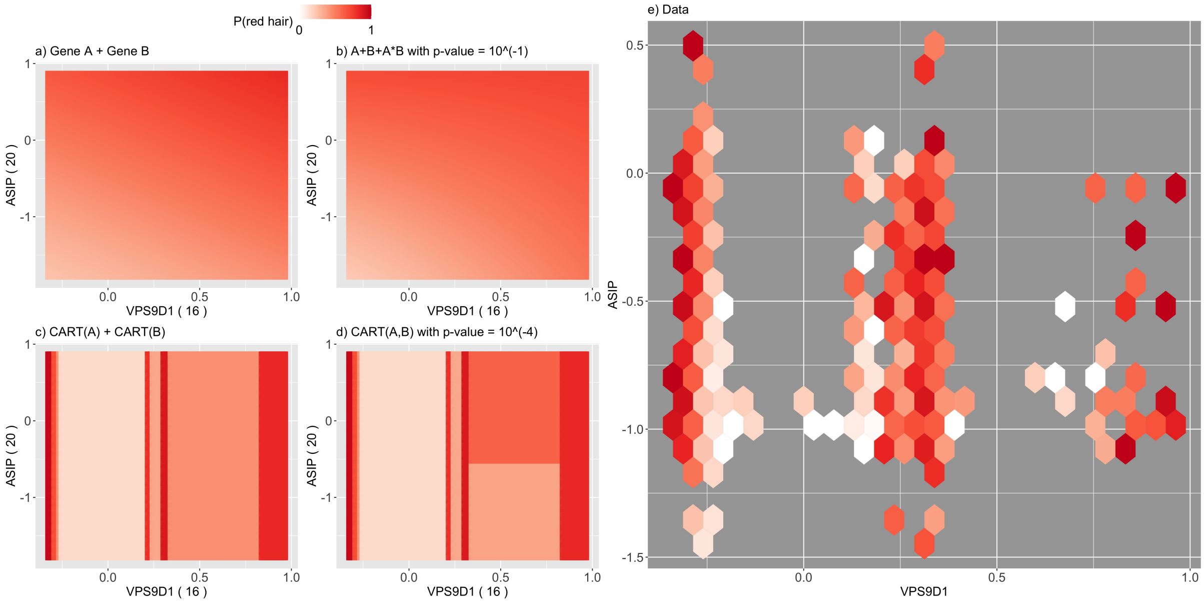

Figure S10: Response surface for *ASIP* - *VPS9D1*, otherwise as Figure ??

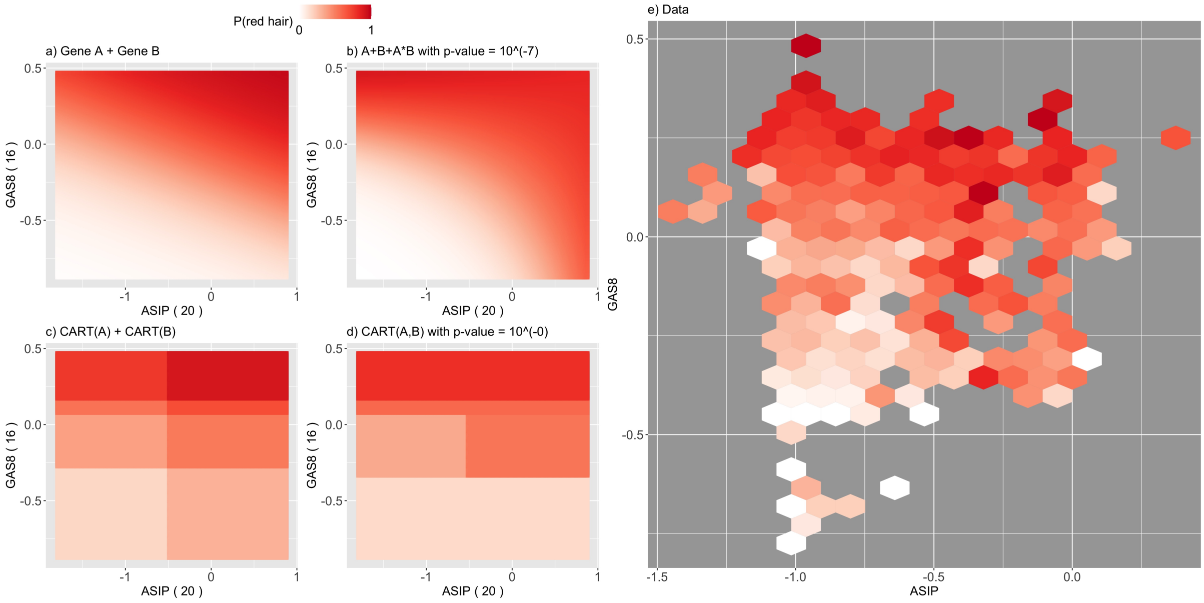

Figure S11: Response surface for *ASIP* - *GAS8*, otherwise as Figure ??

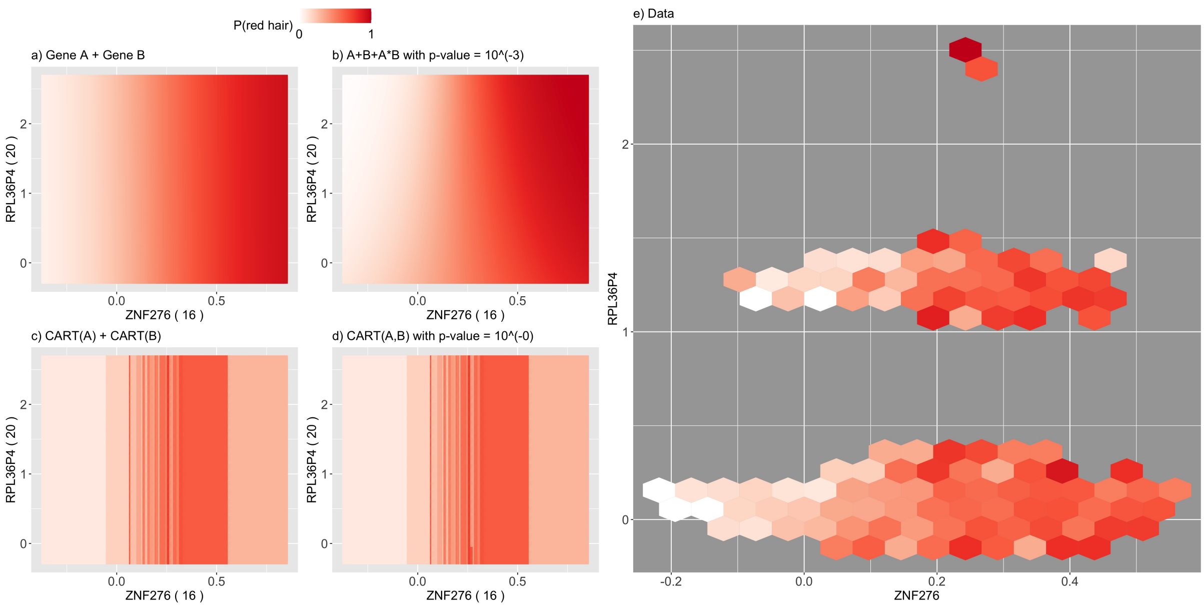

Figure S12: Response surface for *ZNF276* - *RPL36P4*, otherwise as Figure **??**

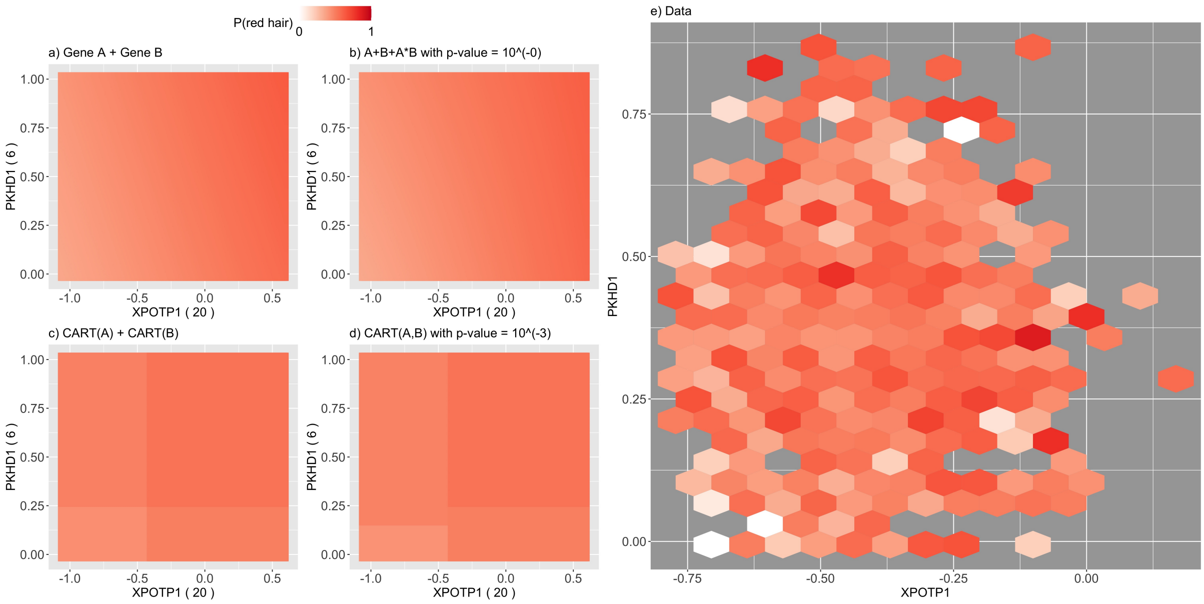

Figure S13: Response surface for *XPOTP1* - *PKHD1*, otherwise as Figure **??**

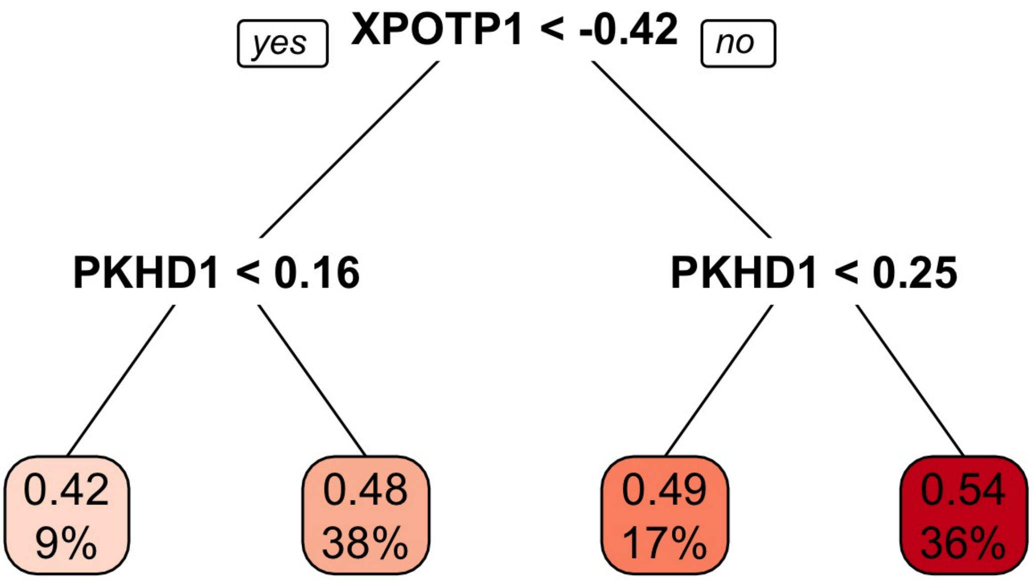

Figure S14: Decision trees for *XPOTP1* - *PKHD1* interaction. The decimal digits at the tip nodes correspond to the predicted probability of red hair. The percentage at the tip nodes corresponds to the percentage of training observations falling into this tip node.

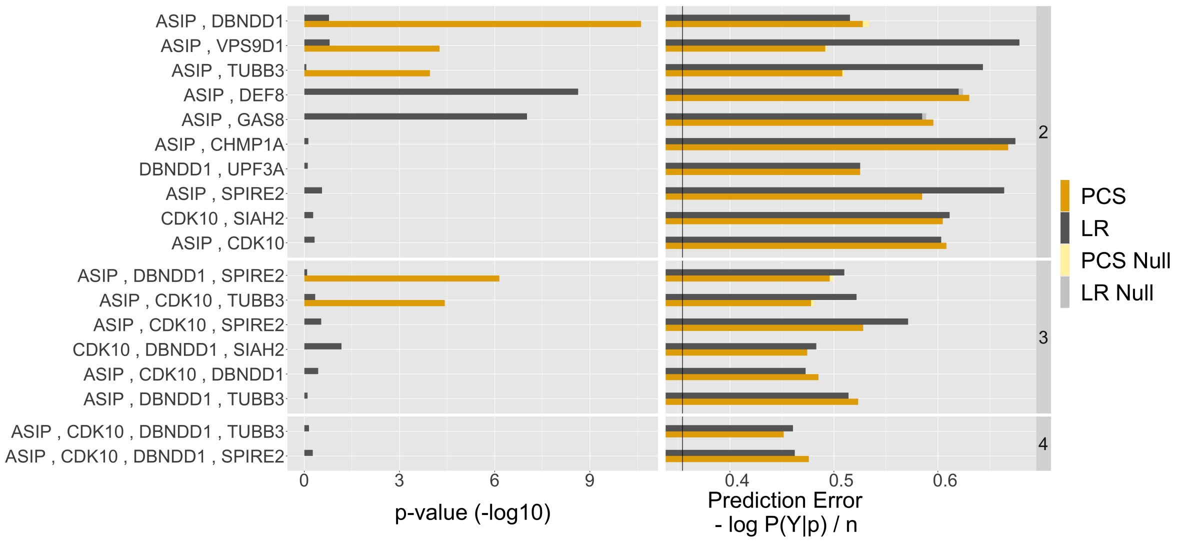

Figure S15: List of stable gene level interactions found by iRF (stability score > 0.5). The first column shows the PCS p-value (orange) and p-value from logistic regression (gray) on a -log10 scale. The second column shows the prediction error (cross-entropy) on the test data of the learned CART (orange) and logistic regression (gray) models for both, no-epistasis (light color) and epistasis (dark color). The black vertical line shows the prediction error achieved by iRF using all the gene features simultaneously.

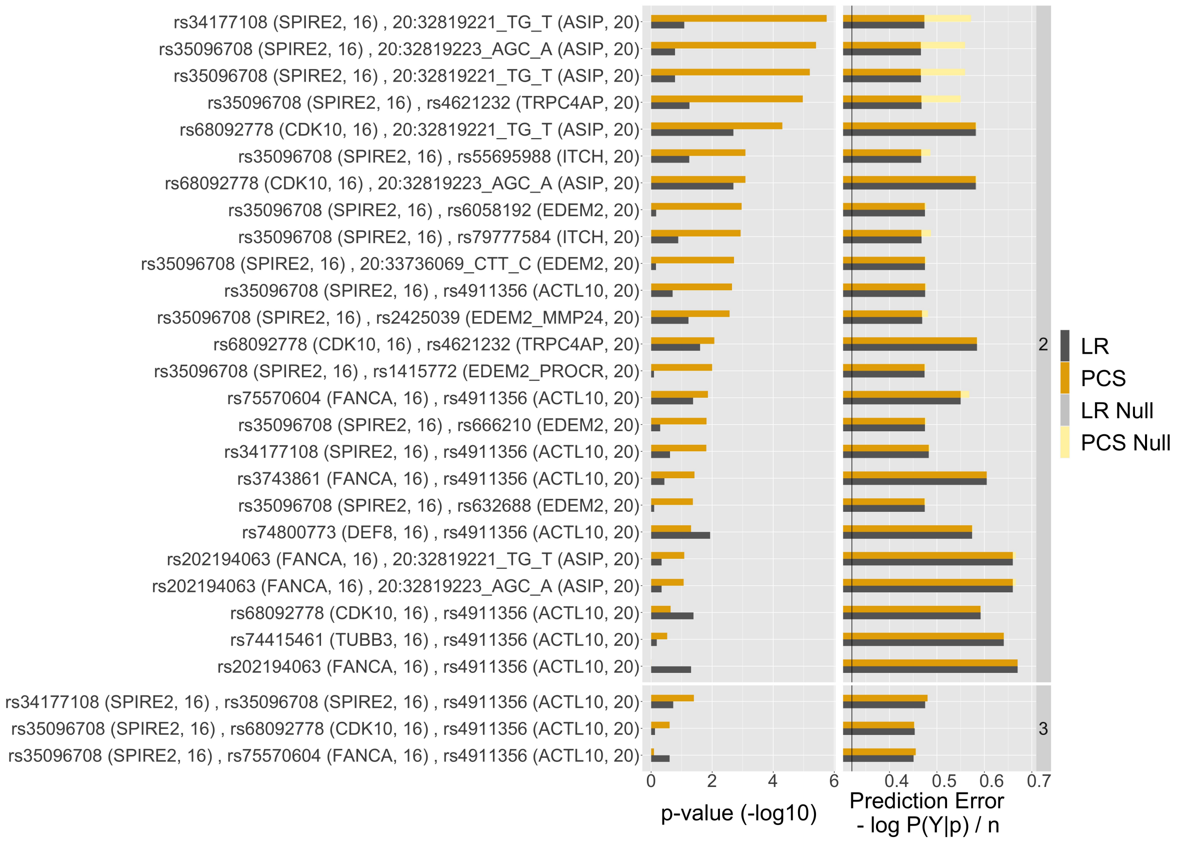

Figure S16: Same as Figure S15, but for the variant level.

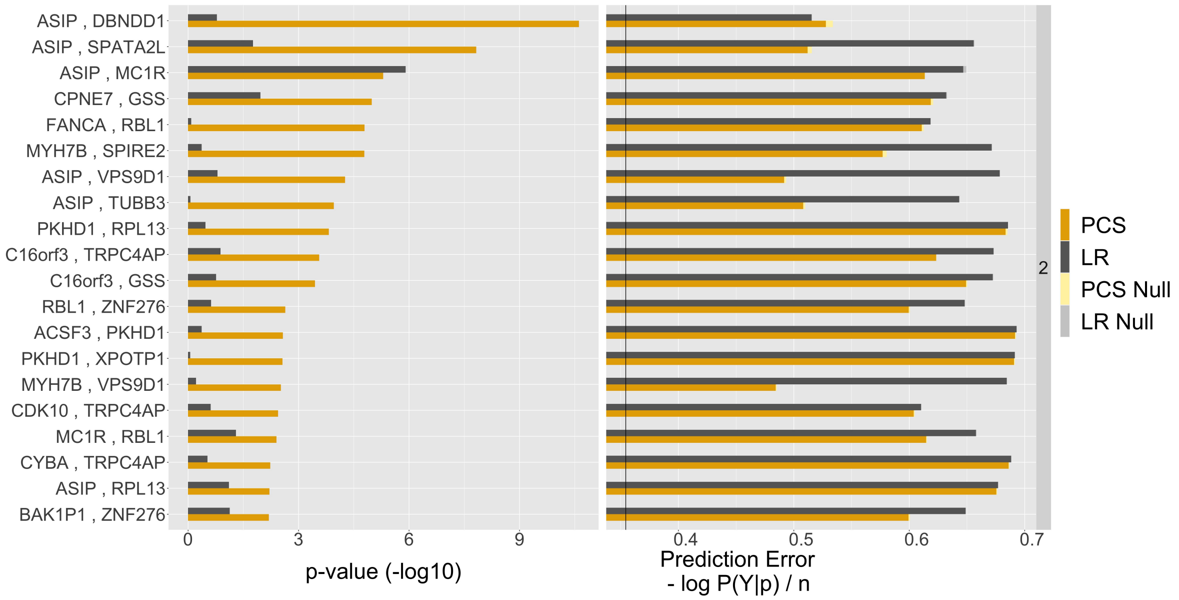

Figure S17: Same as Figure S15, but for the top 20 PCS p-values among inter chromosome pairwise brute force search of top 50 iRF genes (in terms of Gini importance).

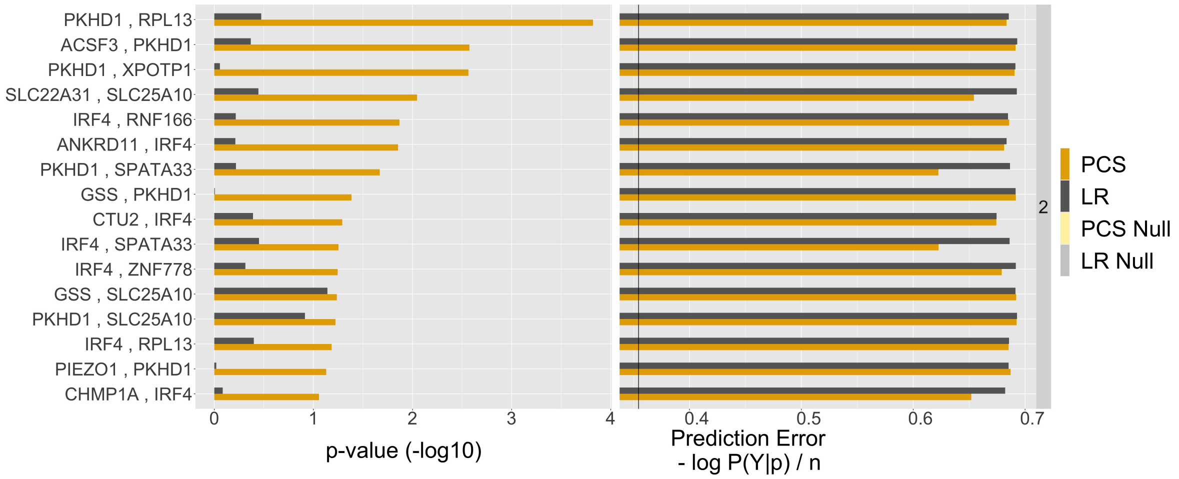

Figure S18: Same as Figure S17 but restricted those interactions which are not between chromosome 16 and chromosome 20 genes.

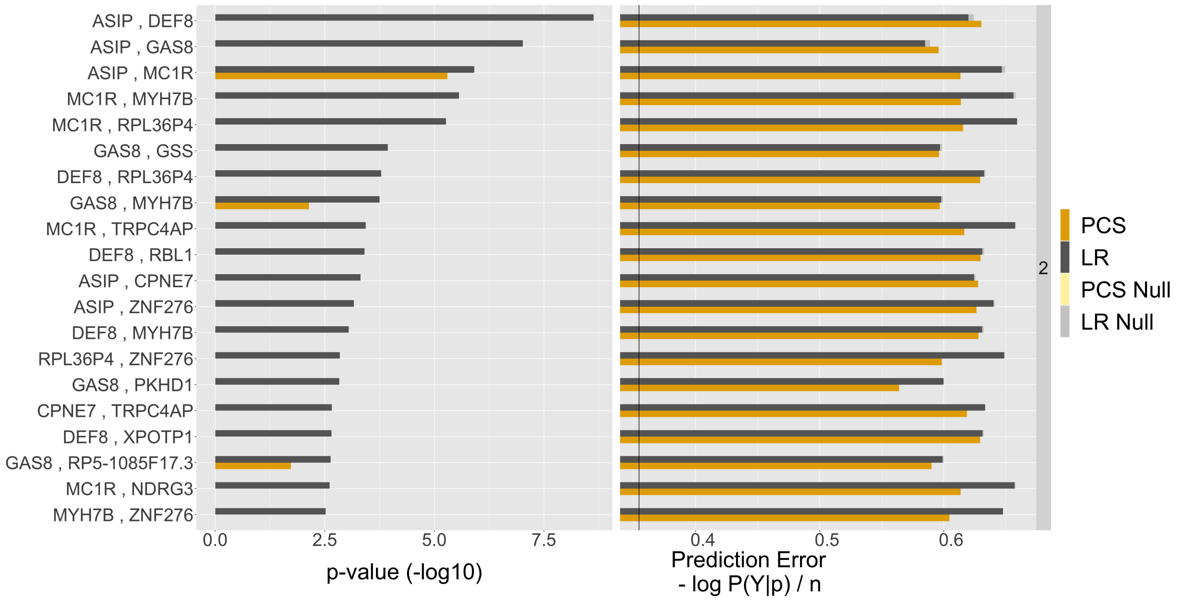

Figure S19: Same as Figure S17 but for top p-values from logistic regression.

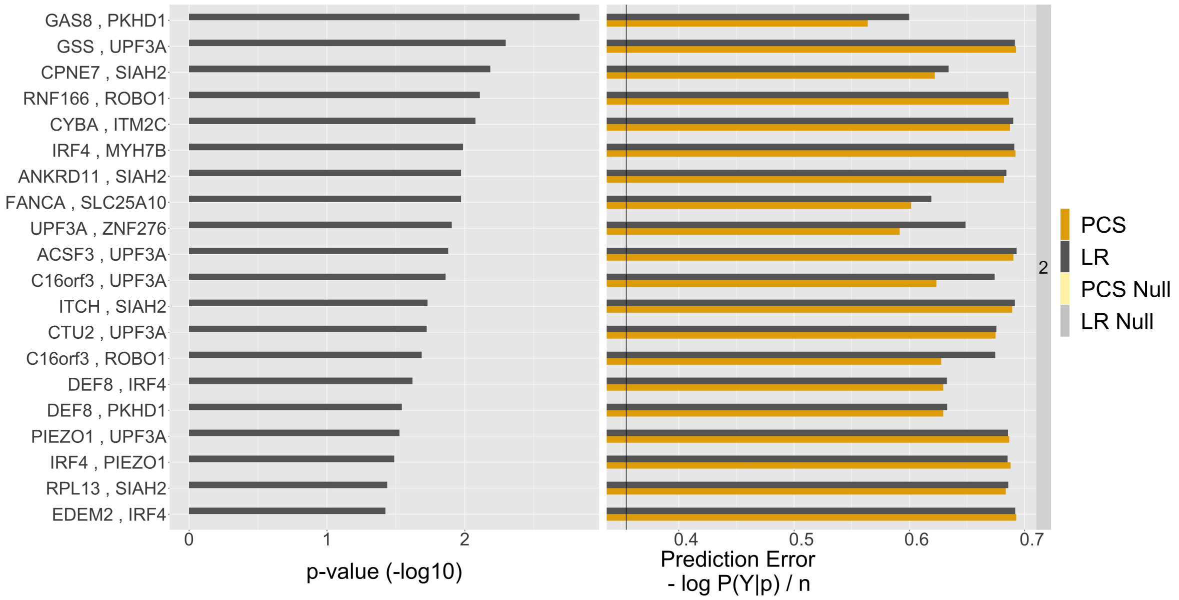

Figure S20: Same as Figure S18 but for top p-values from logistic regression.

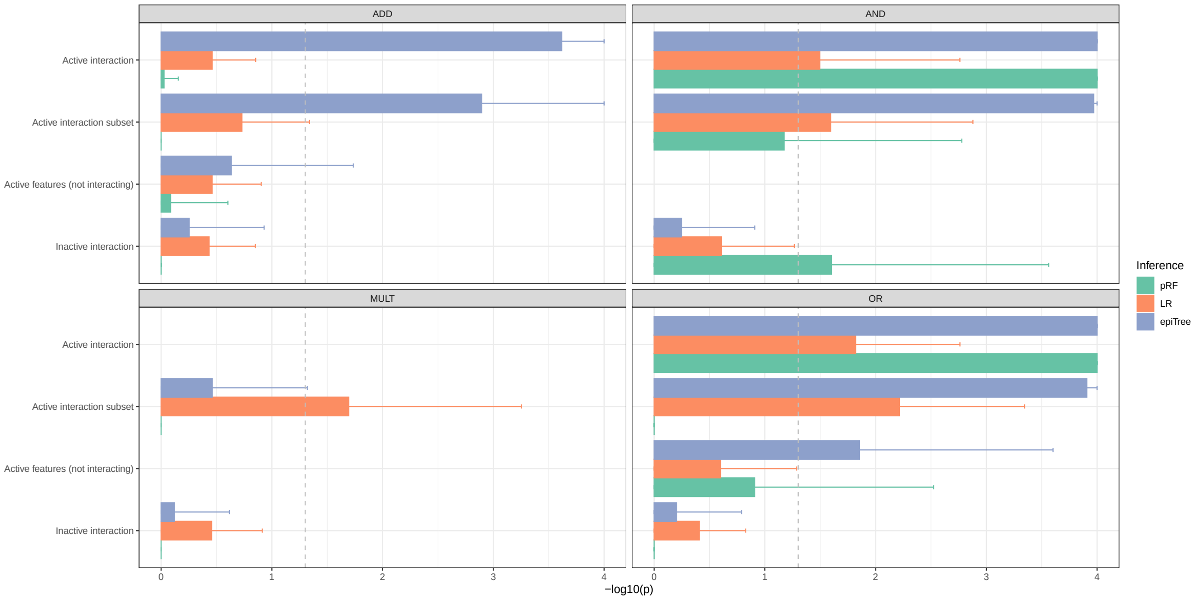

Figure S21: Summary of average p-values across 25 simulation replicates by response model and inference method (pRF: permutation random forests, LR: logistic regression, epiTree: PCS epistasis p-values). Inference was conducted on interactions recovered by iRF. Note: iRF did not detect the full active interaction for the multiplicative model. Candidate interactions are grouped by whether they correspond to (i) a true interaction (ii) a subset of the true interaction (iii) a non-active interaction that includes only active features (iv) a non-active interaction that includes inactive features. Error bars show 1 standard deviation. The grey dashed line corresponds to a significance threshold of 0.05.

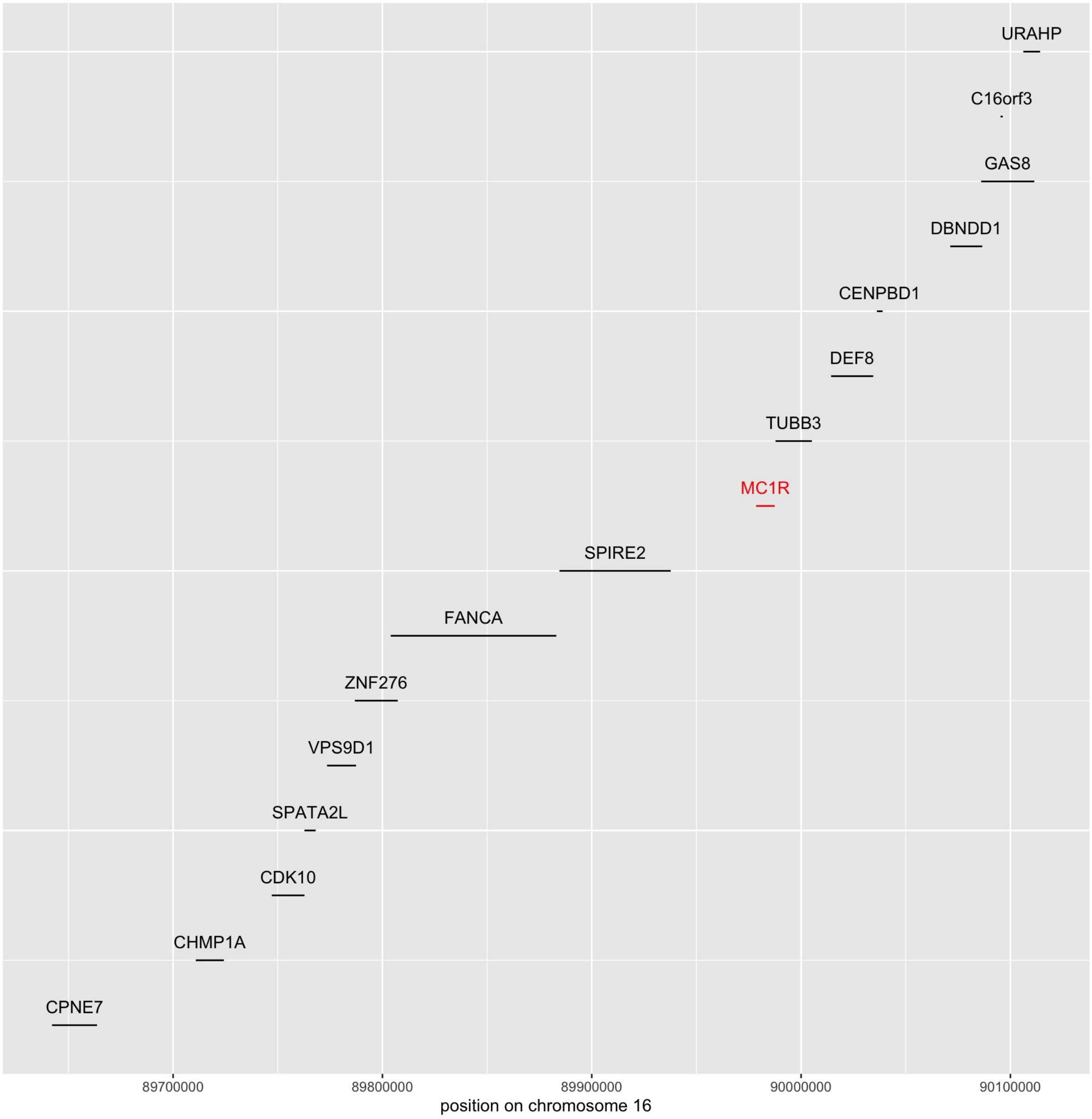

Figure S22: Location of the coding region for the 16 genes which appear in the stable gene level interactions found by iRF that contain only genes on chromosome 16, together with the location of the MC1R gene.

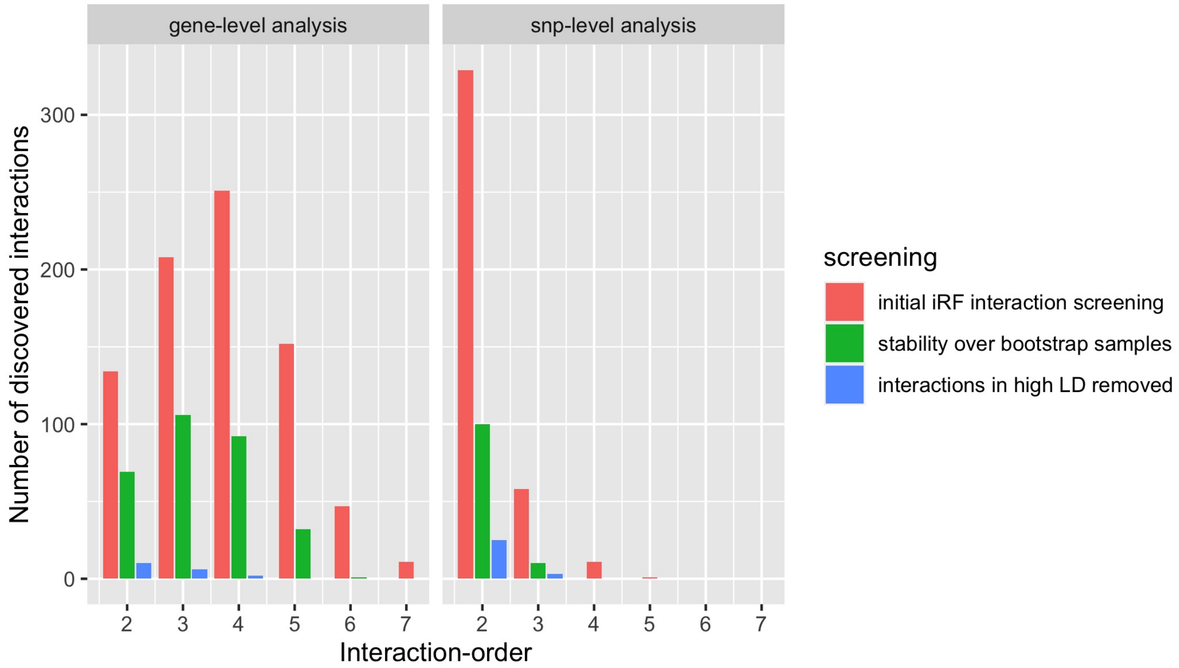

Figure S23: Number of interactions per interaction-order discovered by iRF within individual screening steps (red hair case study). The left plot corresponds to the gene-level analysis and the right plot to the SNP-level analysis. The x-axis shows the order of the interactions and the y-axis the number of discovered interactions for a specific order. Red: Interactions from the initial iRF screening. Green: From those, interactions which are stable among bootstrap replicates. Blue: From those, interactions which are not in high linkage disequilibrium.

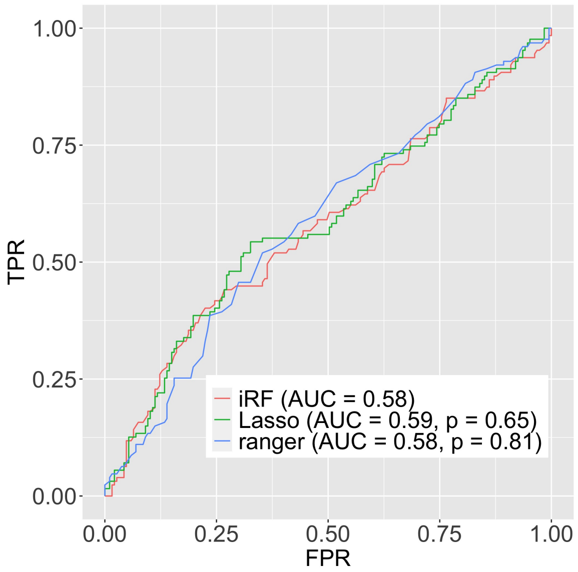

(a) Results for analysis restricted to female subjects.

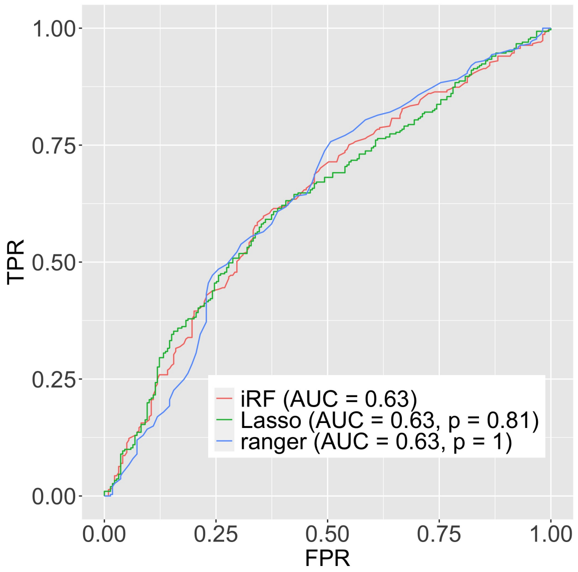

(b) Results for analysis restricted to male subjects.

Figure S24: Analog as Figure ?? A., but for the iRF prediction model and competitors trained on the data from the multiple sclerosis case study in Section S1.8.

| | Interaction | PCS p-value |
|---|---|---|
| 1 | HLA-DRB1 + ZNF771 | 0.081 |
| 2 | HLA-DRB1 + SLC1A6 | 0.094 |

(a) Results for analysis restricted to female subjects.

| | Interaction | PCS p-value |
|---|---|---|
| 1 | HLA-DRB1 + NDUFS2 | 0.0035 |
| 2 | HLA-DRB1 + RP11-54O7.17 | 0.0230 |
| 3 | HLA-DRB5 + AC005224.2 | 0.0340 |
| 4 | HLA-DRB1 + STEAP3 | 0.0340 |
| 5 | HLA-DRB5 + WDR18 + POC5 | 0.0970 |

(b) Results for analysis restricted to male subjects.

Figure S25: List of stable gene level interactions found by iRF (stability score $> 0.5$) in the multiple sclerosis case study from Section S1.8, together with PCS p-value. Only interactions with PCS p-value $< 0.1$ are shown.

### References

[1] A. N. Barbeira, et al. Exploiting the gtex resources to decipher the mechanisms at gwas loci. *Genome biology*, 22:1–24, 2021.

[2] T. M. Beasley, S. Erickson, and D. B. Allison. Rank-based inverse normal transformations are increasingly used, but are they merited? *Behavior genetics*, 39(5):580–595, 2009.

[3] R. Berk, et al. Misspecified mean function regression: Making good use of regression models that are wrong. *Sociological Methods & Research*, 43(3):422–451, 2014.

[4] L. Breiman, et al. *Classification and Regression Trees*. Chapman and Hall, New York, 1984.

[5] C. Bycroft, et al. The UK Biobank resource with deep phenotyping and genomic data. *Nature*, 562(7726):203–209, 2018.

[6] H. J. Cordell. Epistasis: What it means, what it doesn't mean, and statistical methods to detect it in humans. *Human Molecular Genetics*, 11(20):2463–2468, 2002.

[7] H. J. Cordell, et al. Statistical modeling of interlocus interactions in a complex disease: Rejection of the multiplicative model of epistasis in type 1 diabetes. *Genetics*, 158(1):357–367, 2001.

[8] A. Dumitriu, et al. Integrative analyses of proteomics and RNA transcriptomics implicate mitochondrial processes, protein folding pathways and GWAS loci in Parkinson disease. *BMC Medical Genomics*, 9(1):5, December 2015.

[9] J. J. Faraway. Does data splitting improve prediction? *Statistics and Computing*, 26(1-2):49–60, 2016.

[10] R. Foraita, K. Bammann, and I. Pigeot. Modeling gene-gene interactions using graphical chain models. *Human Heredity*, 65(1):47–56, 2008.

[11] F. Girosi and T. Poggio. Representation properties of networks: Kolmogorov's theorem is irrelevant. *Neural Computation*, 1(4):465–469, 1989.

[12] I. B. Hallgrímsdóttir and D. S. Yuster. A complete classification of epistatic two-locus models. *BMC Genetics*, 9(1), 2008.

[13] T. Hastie and R. Tibshirani. Generalized additive models. *Statistical Science*, 1(3):297–318, 1986.

[14] J. Hathaway, et al. Diagnostic yield of genetic testing in a heterogeneous cohort of 1376 HCM patients. *BMC Cardiovascular Disorders*, 21(1):126, December 2021.

[15] Y. Huang, S. Wuchty, and T. M. Przytycka. eQTL epistasis – challenges and computational approaches. *Frontiers in Genetics*, 4:51, 2013.

[16] O. Kobiler, et al. Quantitative kinetic analysis of the bacteriophage genetic network. *Proceedings of the National Academy of Sciences*, 102(12):4470–4475, 2005.

[17] H. Leeb. Conditional predictive inference post model selection. *The Annals of Statistics*, 37(5B):2838–2876, 2009.

[18] E. Levine and T. Hwa. Small RNAs establish gene expression thresholds. *Current Opinion in Microbiology*, 11(6):574–579, 2008.

[19] J. Li, et al. Detecting gene-gene interactions using a permutation-based random forest method. *BioData mining*, 9(1):1–17, 2016.

[20] J. W. Little. Threshold effects in gene regulation: When some is not enough. *Proceedings of the National Academy of Sciences*, 102(15):5310–5311, 2005.

[21] J. W. Little, D. P. Shepley, and D. W. Wert. Robustness of a gene regulatory circuit. *The EMBO Journal*, 18(15):4299–4307, 1999.

[22] M. D. Morgan, et al. Genome-wide study of hair colour in UK Biobank explains most of the SNP heritability. *Nature Communications*, 9:5271, 2018.

[23] S. V. Naoaev. Some limit theorems for large deviation. *Theory of Probability and its Applications*, 10(2):214–235, 1965.

[24] B. V. North, D. Curtis, and P. C. Sham. Application of logistic regression to case-control association studies involving two causative loci. *Human Heredity*, 59(2):79–87, 2005.

[25] S. Pudewell, et al. Accessory proteins of the RAS-MAPK pathway: Moving from the side line to the front line. *Communications Biology*, 4(1):696, December 2021.

[26] Z. R. Sailer and M. J. Harms. Detecting high-order epistasis in nonlinear genotype-phenotype maps. *Genetics*, 205(3):1079–1088, 2017.

[27] S. H. Spiezio, et al. Genetic divergence and the genetic architecture of complex traits in chromosome substitution strains of mice. *BMC Genetics*, 13(1):38, December 2012.

[28] C. Tcheandjieu, et al. A phenome-wide association study of 26 mendelian genes reveals phenotypic expressivity of common and rare variants within the general population. *PLOS Genetics*, 16(11):e1008802, November 2020.

[29] A. T. Timberlake, et al. Two locus inheritance of non-syndromic midline craniosynostosis via rare SMAD6 and common BMP2 alleles. *eLife*, 5:e20125, September 2016.

[30] M. Ueki and H. J. Cordell. Improved statistics for genome-wide interaction analysis. *PLOS Genetics*, 8(4):e1002625, 2012.

[31] G. Wahba. Bayesian "confidence intervals" for the cross-validated smoothing spline. *Journal of the Royal Statistical Society: Series B (Methodological)*, 45(1):133–150, 1983.

[32] L. Wasserman, A. Ramdas, and S. Balakrishnan. Universal inference. *Proceedings of the National Academy of Sciences*, 117(29):16880–16890, 2020.

[33] X. Wu, et al. A novel statistic for genome-wide interaction analysis. *PLoS Genetics*, 6(9):e1001131, 2010.

[34] B. Yu and K. Kumbier. Veridical data science. *Proceedings of the National Academy of Sciences*, 117(8):3920–3929, 2020.
